# Structural basis of human antibody recognition of linear and conformational epitopes on the peanut allergen Ara h 2

**DOI:** 10.64898/2026.11.24.768991

**Authors:** Alesandro Civljak, Weslley Fernandes-Braga, Bruno Bonnettaz, Mariana Miranda-Waldetario, Iden A. Sapse, Galina Grishina, Hugh A. Sampson, Maria A. Curotto de Lafaille, Goran Bajic

## Abstract

IgE-mediated food allergy requires allergens to crosslink FcεRI-bound IgE on mast cells, yet how individual allergen epitopes organize pathogenic IgE complexes remains poorly understood. We defined the structural basis of antibody recognition of Ara h 2, a major peanut allergen, using three human monoclonal antibodies isolated from IgG1 memory B cells of peanut-allergic children. As IgE, the DPYSPS repeat-specific antibody P4EC10 alone induced degranulation in vitro and passive cutaneous anaphylaxis in vivo, whereas P1BC6 and P1EC5 elicited both responses only when combined. P4EC10 IgG1 inhibited a mean of 46% of Ara h 2 binding to plasma IgE in the highest Ara h 2-specific IgE stratum and 24% in the lowest. Cryo-EM revealed 2:1 and 3:1 P4EC10 Fab–Ara h 2 complexes. In the 2:1 structure, two Fabs bound adjacent linear DPYSPS repeats within the flexible loop and contacted one another. P1BC6 and P1EC5 each bound Ara h 2 with 1:1 stoichiometry and simultaneously engaged distinct conformational epitopes. Solution binding measurements supported these stoichiometries and indicated that proline hydroxylation enhanced P4EC10 binding. These findings show that Ara h 2 can drive effector-cell activation by recruiting a single IgE specificity to repeated epitopes or by engaging distinct IgE specificities at separate conformational epitopes.

**Summary:** Cryo-EM resolved human antibodies bound to linear loop epitopes and conformational epitopes on Ara h 2. Repeat-specific IgE alone supported degranulation, while IgG1 pairs targeting both epitope types most effectively inhibited degranulation in cells sensitized with pooled patient plasma.

## INTRODUCTION

Food allergies affect one in ten persons in the U.S. population. Four out of the ten affected individuals are at risk of severe disease and report a history of life-threatening anaphylaxis^1^. Peanut allergy can persist beyond childhood and cause severe immediate reactions. Ara h 2, a major peanut 2S albumin, is an important target of IgE and a potent elicitor of allergic effector-cell activation ^2–4^. Its disulfide-stabilized helical core and flexible repeat-containing loop present multiple antibody-binding surfaces. Understanding how antibodies assemble on these surfaces can connect molecular recognition with the functional capacity of allergen-specific IgE ^5–9^.

Immediate allergic reactions occur when an allergen engages allergen-specific IgE molecules prebound to the high-affinity IgE receptor FcεRI on mast cells and basophils. Productive signaling requires allergen-mediated cross-linking of IgE-FcεRI complexes and therefore depends on the capacity of an allergen to engage multiple receptor-bound IgE molecules^10–14^. Such functional polyvalency may arise from polyclonal IgE responses directed to distinct epitopes on a single allergen, from a monoclonal IgE specificity directed against repeated determinants within one allergen molecule, such as the hydroxyproline-containing repeat loop of Ara h 2, or from repeated epitope display by oligomeric allergens. Thus, the geometry, copy number, and spatial organization of allergen-bound IgE molecules are likely to be major determinants of effector-cell activation.

Recent single-cell analyses have begun to define the cellular and molecular basis of this response. Human monoclonal IgE antibodies isolated from unrelated peanut-allergic individuals show striking convergent evolution toward a recurrent Ara h 2 epitope centered on the repeated DPYSPS-containing sequence^15–17^. These antibodies bind Ara h 2 with high affinity and, remarkably, some individual monoclonal IgE antibodies can drive allergen-dependent mast-cell degranulation and systemic anaphylaxis in humanized mice. Here we used biological assays and physical methods to resolve how individual pathogenic antibodies physically engage a native peanut allergen, Ara h 2, to generate receptor-cross-linking assemblies.

Type 2-associated IgG memory B-cell populations with the capacity to contribute to IgE responses have been described independently by Ota et al.^18^ and Koenig et al.^19^. Ota et al. identified high-affinity Ara h 2-specific clones within CD23-positive IgG1 memory B cells of peanut-allergic children, providing the source of the antibody variable regions examined here. Public clonal families of Ara h 2-specific antibodies containing IgE and IgG members in different individuals were identified, evidence of strong convergence of the antibody to this molecule (refs).

The immunodominant Ara h 2 region is not solely a linear peptide target. Studies using chimeric 2S albumins localized a dominant conformational IgE-binding region to residues 33-81, overlapping the hydroxyproline-containing repeat loop and identifying this segment as critical for mast-cell degranulation^5,6^. Previous studies established that repeat-directed monoclonal IgE can trigger effector-cell activation and that antibodies recognizing complementary Ara h 2 epitopes can support or inhibit allergic responses^7,16,17,20^. A more specific question is how neighboring antibodies pack on the repeat loop and how that arrangement compares with simultaneous occupancy of separate surfaces on the same allergen. We addressed this question using P4EC10, P1BC6, and P1EC5, three antibodies originally isolated from IgG1 memory B cells. We combined Fab–allergen cryo-EM structures with solution binding measurements, functional assays of recombinant IgE versions, and inhibition assays using individual and pooled patient plasma. To our knowledge, these are the first cryo-EM structures of Ara h 2 in complex with human antibody Fabs. They show how two P4EC10 Fabs contact one another on adjacent DPYSPS repeats and how P1BC6 and P1EC5 simultaneously engage distinct surfaces of the folded allergen. These arrangements provide structural context for the functional capacities of the corresponding recombinant IgEs.

## RESULTS

### Recombinant IgE versions of three memory B cell antibodies show distinct effector activities

We previously identified Ara h 2-specific IgG1 memory B cells, including some belonging to ‘public’ clonal specificities shared among peanut-allergic individuals^18^. Members of two clonal families recognized distinct linear peptides of Ara h 2 which are also main IgE-epitopes. Additional clones targeted epitopes not represented in a linear Ara h 2 peptide array. To examine how these distinct modes of antigen recognition influence IgE function, we selected three representative antibodies: P4EC10, which recognizes the DPYSPS repeat-containing region; P1BC6, which recognizes an N-terminal linear peptide; and P1EC5, which binds full-length Ara h 2 but not linear peptides of Ara h 2. To assess their effector potential, we expressed the three IgG1 memory B cell-derived variable-region pairs as human IgE. P4EC10 belongs to the PA13P1E10-related convergent antibody family, which includes antibodies recovered from naturally occurring IgE-producing plasma cells in peanut-allergic individuals^15,18^. Related repeat-directed IgE antibodies have also been shown to induce mast-cell degranulation and anaphylaxis^16^. Our P4EC10 experiments therefore characterize the effector capacity of a representative of an antibody family already documented in the human IgE response.

All three IgE mAbs bound peanut extract and both natural and recombinant Ara h 2 (nAra h 2 and rAra h 2, respectively; **Fig. 1A**). In addition, P4EC10 recognized synthetic peptides containing the sequence RDEDSYGRDPYSPSQDPYSPSQDPDRRDPYSPSPYD, which includes three copies of the DPYSPS motif, in both its non-hydroxylated and hydroxylated forms (DPYSPS and DPYSP^(OH)^S, respectively), whereas P1EC5 and P1BC6 showed no detectable binding to these peptides. None of the three mAbs bound natural Ara h 1 (nAra h 1). P4EC10 showed only minimal binding to natural Ara h 3 (nAra h 3), while all three antibodies exhibited low-level reactivity to natural Ara h 6 (nAra h 6).

**Figure 1.**
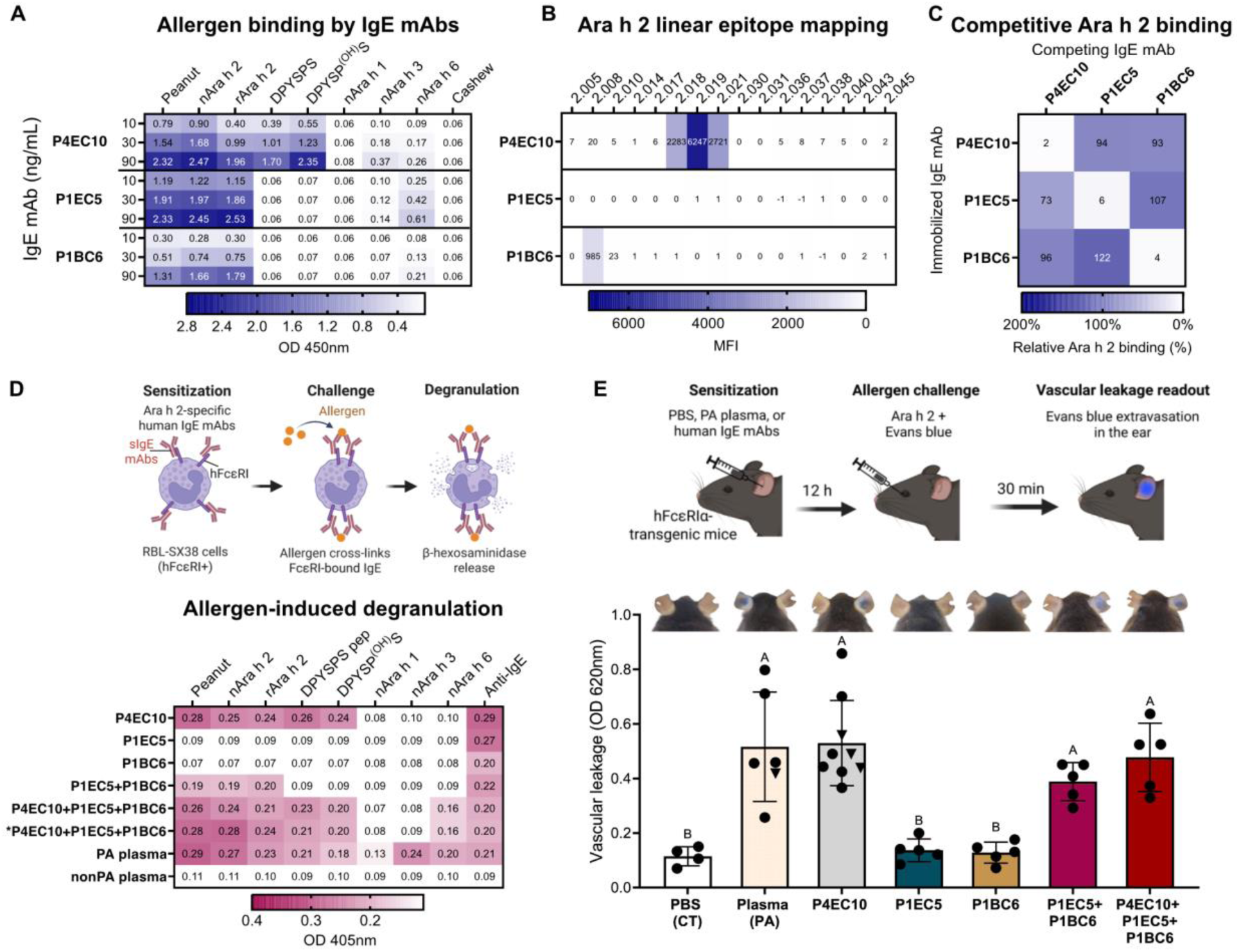
Epitope characterization and functional activity of recombinant human Ara h 2-specific monoclonal antibodies. **(A)** ELISA analysis of recombinant human IgE monoclonal antibody binding to peanut extract, natural and recombinant Ara h 2, natural Ara h 1, Ara h 3, Ara h 6, cashew extract, and Ara h 2-derived synthetic peptides. IgE mAbs were tested at 10, 30, and 90 ng/mL. Heatmap values represent OD 450nm. **(B)** Linear epitope mapping of recombinant human IgE mAbs using an overlapping Ara h 2 peptide library. Heatmap values represent mean fluorescence intensity (MFI). **(C)** Competitive ELISA measuring Ara h 2 binding to immobilized recombinant human IgE mAbs in the presence of homologous or heterologous competing antibodies. The heatmap shows relative Ara h 2 binding in the presence of a 3 μg/mL competing IgE mAb, normalized to binding in the presence of an irrelevant human monoclonal antibody. Complete inhibition curves are shown in Fig. S2. **(D)** β-hexosaminidase release from human FcεRI-expressing RBL-SX38 cells sensitized with individual or combined recombinant human IgE mAbs and challenged with the indicated peanut allergens or Ara h 2-derived peptides (0.5 μg/mL). IgE mAbs were used at 0.5 µg/mL, except in the asterisk-marked condition, in which P4EC10 was maintained at 0.5 µg/mL and P1EC5 and P1BC6 were each added at 0.25 µg/mL. Heatmap values represent OD450. **(E)** Passive cutaneous anaphylaxis in hFcεRIα-transgenic mice sensitized intradermally with the indicated recombinant human IgE mAbs and challenged intravenously with Ara h 2. Circles indicate natural Ara h 2 and triangles indicate recombinant Ara h 2. Representative ears are shown above. Evans blue extravasation was quantified by absorbance at 620 nm. Each symbol represents one sensitized ear from an individual mouse. Data are presented as mean ± SEM. Statistical analysis was performed by one-way ANOVA followed by Tukey’s multiple-comparison test. Groups not sharing a common letter are significantly different (*P* < 0.05).

Linear Ara h 2 epitope mapping of the three IgE mAbs was consistent with the binding profiles previously described for the corresponding IgG antibodies^18^. P4EC10 bound predominantly to peptides spanning the DPYSPS repeat-containing region, spanning peptides Ara h 2.018, 2.019, and 2.021 (RDDSYERDPYSPSQ, DSYERDPYSPSQDPY and PYSPSQDPYSPSPYD respectively) (**Fig. 1B**). In contrast, P1BC6 bound selectively to the N-terminus Ara h 2.008 peptide (WELQGDRRCQSQLER), showing no detectable reactivity to peptides spanning the repeat-containing region. P1EC5 did not bind any peptide in the array, consistent with recognition of a conformational epitope. None of the three IgE mAbs showed substantial binding to linear peptides from Ara h 1, Ara h 3, or Ara h 6 (**Fig. S1**).

We used competition ELISA to determine whether the three antibodies recognized overlapping antigenic regions. Each IgE mAb efficiently inhibited Ara h 2 binding to its homologous immobilized antibody, whereas heterologous competitors had little effect on binding (**Fig. 1C**). Concentration-dependent competition assays confirmed complete self-competition but minimal competition between antibodies recognizing different epitopes (**Fig. S2**). These assays, therefore, distinguish three largely non-competing antibody specificities.

We next evaluated the functional consequences of these distinct binding profiles in basophil activation assays using the RBL-SX38 cell line expressing human FcεRI. P4EC10-sensitized basophils underwent robust degranulation following stimulation with peanut extract, nAra h 2, rAra h 2, or hydroxylated or non-hydroxylated peptides containing the repeated DPYSPS motif (**Fig. 1D**). In contrast, P1EC5 or P1BC6 alone failed to induce detectable degranulation despite efficient binding to Ara h 2. Co-sensitization with P1EC5 and P1BC6 induced allergen-dependent degranulation, indicating that simultaneous engagement of two distinct Ara h 2 epitopes is sufficient to crosslink FcεRI and activate effector cells. Co-sensitizing basophils with equimolar amounts of P4EC10, P1EC5, and P1BC6 produced a degranulation profile similar to that observed with P4EC10 alone. Neither Ara h 1 nor Ara h 3 induced detectable degranulation under these conditions. Ara h 6 induced low-level degranulation in basophils sensitized with all three IgE mAbs (**Fig. 1D**), likely reflecting the combined weak reactivity of the three antibodies with Ara h 6 observed in the binding assays (**Fig. 1A**).

We next evaluated the in vivo relevance of these findings using passive cutaneous anaphylaxis (PCA). We sensitized human FcεRIα-transgenic (hFcεRIα) mice by intradermal injection in the ear of individual IgE mAbs or the indicated antibody combinations, followed by intravenous challenge with Ara h 2 and the Evans blue dye (**Fig. 1E**). Consistent with the basophil degranulation assays, P4EC10 alone induced marked vascular leakage (evidenced by Evans blue extravasation) when challenged with nAra h 2 or rAra h 2, whereas P1EC5 and P1BC6 individually induced minimal vascular leakage. Co-sensitization with P1EC5 and P1BC6 induced a robust PCA response, and sensitization with all three IgE mAbs produced a similar robust response. Thus, the relative activities of the individual and combined IgE antibodies were consistent across the *in vitro* and *in vivo* assays.

Under the conditions tested, P4EC10 alone and the P1BC6–P1EC5 combination supported RBL-SX38 degranulation and passive cutaneous anaphylaxis. These results are consistent with previously described activation by repeat-directed or complementary IgE specificities (refs) and establish functional activity for the antibody sequences analyzed structurally here.

### Ara h 2-specific IgG antibodies produce heterogeneous plasma IgE inhibition profiles

Having identified distinct Ara h 2 epitope configurations capable of supporting IgE-mediated effector-cell activation, we next examined the contribution of these antigenic epitopes to polyclonal IgE recognition in peanut-allergic individuals. IgG1 versions of P4EC10, P1EC5, and P1BC6 were used as epitope-specific competitors to inhibit plasma IgE binding to Ara h 2. Plasma samples from 24 peanut-allergic individuals were stratified according to Ara h 2-specific IgE (sIgE) levels (>100, 40–60, 10–20, and 4–8 kU_A_/L; n = 6 per group), and IgE binding to Ara h 2 was measured in the presence of the individual IgG1 mAbs or the indicated combinations (**Fig. S3** and **Fig. 2A–C**). Inhibition in this assay measures reduced allergen binding to captured plasma IgE. Because the competitors are intact IgG1 antibodies, it cannot be ruled out that in addition to the direct epitope competition, there is steric interference with nearby binding sites, thus the assay does not directly quantify the proportion of IgE directed to the identical epitope.

**Figure 2.**
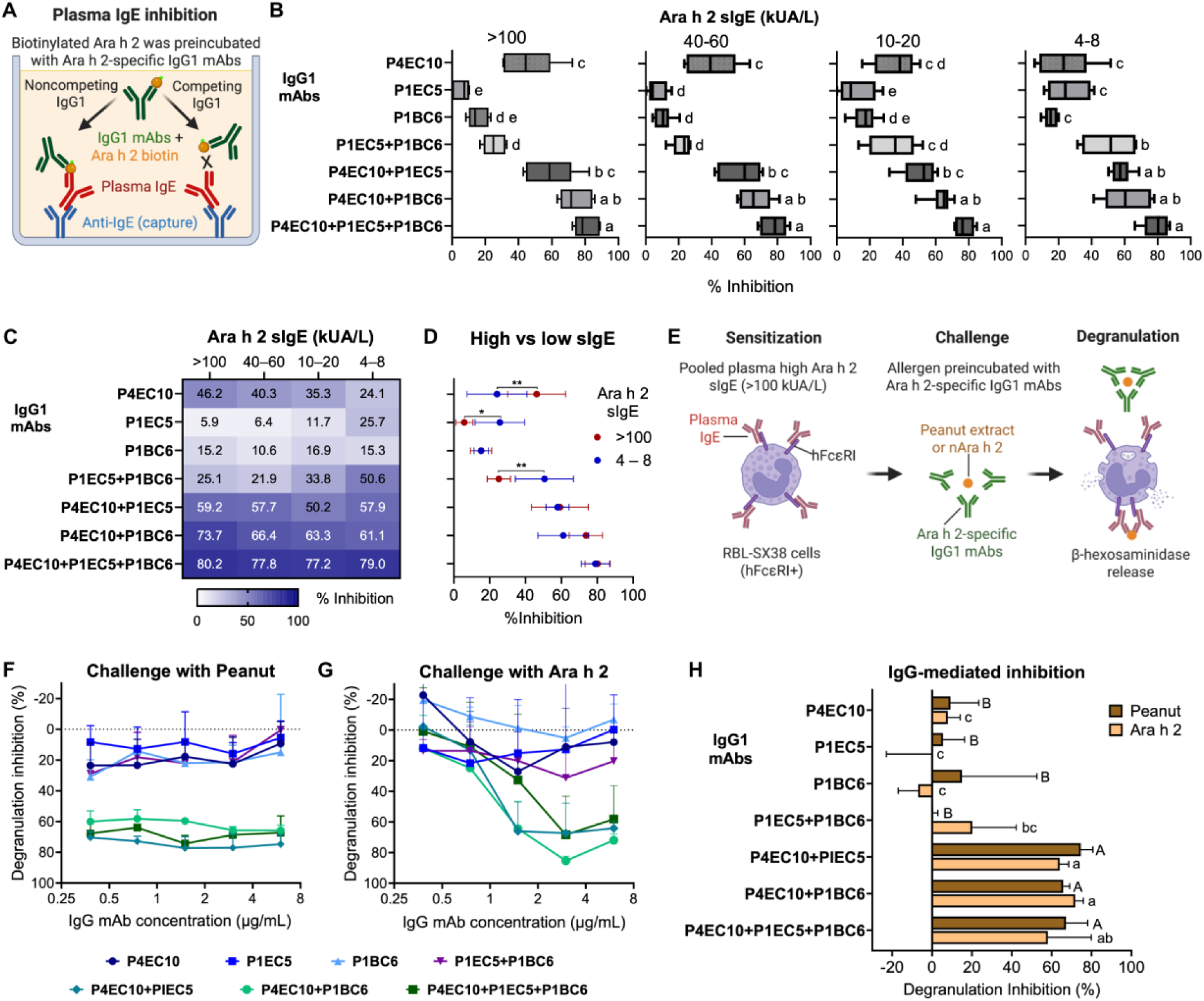
Epitope-specific competition reveals differences in polyclonal Ara h 2 IgE recognition and effector-cell activation. **(A)** Schematic representation of the ELISA used to measure the inhibition of plasma IgE binding to Ara h 2 by recombinant human IgG monoclonal antibodies. **(B)** Distribution of IgE-binding inhibition for the six individuals within each Ara h 2 sIgE group. IgG conditions were compared separately within each sIgE group by ordinary one-way ANOVA followed by Tukey’s multiple-comparisons test. Different letters indicate statistically significant differences among IgG conditions (P < 0.05); conditions sharing at least one letter are not significantly different. **(C)** Mean inhibition of plasma IgE binding to Ara h 2 by recombinant human IgG1 versions of P4EC10, P1EC5, and P1BC6, alone or in the indicated combinations; schema of the assay is shown in the top panel and experimental values in the bottom panel. Plasma samples from peanut-allergic individuals were stratified by Ara h 2-specific IgE (sIgE) level (>100, 40–60, 10–20, and 4–8 kUA/L; n = 6 per group). IgG1 mAbs were used at a total concentration of 3 μg/mL. Heatmap values indicate mean percent inhibition within each sIgE group. **(D)** Direct comparison of IgE-binding inhibition between individuals with Ara h 2 sIgE >100 kUA/L and 4–8 kUA/L. Statistical analysis was performed by ordinary two-way ANOVA followed by Šídák’s multiple-comparisons test comparing the two sIgE groups within each IgG condition. *P < 0.05; **P < 0.01. **(E)** Schematic of the IgG-mediated degranulation-inhibition assay. RBL-SX38 cells expressing human FcεRI were sensitized with pooled plasma from peanut-allergic individuals with Ara h 2 sIgE >100 kUA/L. Peanut extract or natural Ara h 2 was preincubated with Ara h 2-specific IgG1 mAbs before challenge, and degranulation was measured by β-hexosaminidase release. **(F, G)** Concentration-dependent inhibition of degranulation induced by peanut extract **(F)** or natural Ara h 2 **(G)** after preincubation with the indicated IgG1 mAbs or combinations. For antibody combinations, the indicated concentration represents the total IgG1 concentration, divided equally among the component mAbs. **(H)** Degranulation inhibition at 6 μg/mL total IgG1 following challenge with peanut extract or natural Ara h 2. Data in (**F**–**H**) are shown as mean ± SD from three independent experiments. In **H**, IgG1 treatments were compared within each allergen challenge by ordinary two-way ANOVA followed by Tukey’s multiple-comparisons test. Groups that do not share a letter differ significantly (P < 0.05). Uppercase letters denote comparisons within the peanut extract challenge and lowercase letters denote comparisons within the Ara h 2 challenge.

The three-antibody mixture inhibited mean plasma IgE binding by 77.2–80.2% across the four sIgE strata (**Fig. 2B,C**). In the highest and lowest strata, mean inhibition by P4EC10 was 46.2% and 24.1%, respectively. Corresponding values were 5.9% and 25.7% for P1EC5 and 25.1% and 50.6% for the P1EC5–P1BC6 combination **(Fig. 2C,D**). These comparisons should be interpreted with caution because each stratum included only six participants and plasma was tested at a different dilution in each stratum.

We next examined whether IgG1-blocking of these epitopes could interfere with polyclonal IgE-mediated effector-cell activation. RBL-SX38 cells expressing human FcεRI were sensitized with pooled plasma from peanut-allergic individuals with Ara h 2 sIgE >100 kU_A_/L. The cells were then challenged with peanut extract or nAra h 2 preincubated with increasing concentrations of individual IgG1 mAbs or the indicated mAb combinations (**Fig. 2E**). Despite inhibiting a substantial fraction of plasma IgE binding in this group (**Fig. 2A-C**), P4EC10 IgG1 alone had little effect on basophil degranulation mediated by patient plasma IgE. P1EC5 and P1BC6 were similarly ineffective individually, and their combination produced only limited inhibition (**Fig. 2F,G**).

In contrast, combining P4EC10 with either P1EC5 or P1BC6 IgG1 mAbs markedly inhibited degranulation induced by both peanut extract and nAra h 2 (**Fig. 2F,G**). Combining all three IgG1 mAbs produced a similar level of inhibition, with no substantial increase over the P4EC10-containing pairs. At 6 µg/mL of total IgG1, the P4EC10-containing combinations strongly inhibited responses to both peanut extract and Ara h 2, whereas individual antibodies and the P1EC5–P1BC6 combination showed substantially less activity (**Fig. 2H**).

While P4EC10 IgG1 was necessary to inhibit degranulation in these assays, it was not sufficient, and a second non-competing blocking antibody was necessary. Thus, inhibition of cell-free IgE binding and inhibition of IgE-mediated cellular activation were not interchangeable readouts. Inhibition of responses to peanut extract by blocking Ara h 2 epitopes is consistent with the contribution of Ara h 2 to peanut allergenicity(refs); however it does not establish a universal requirement for these particular epitope combinations or exclude the contribution of cross-reactivity with Ara h 6.

### Repeated Ara h 2 epitopes support multivalent P4EC10 binding

We asked how the repeat-containing region of Ara h 2 supports engagement by a single antibody specificity. Based on the functional activity of P4EC10 and its specific recognition of peptides containing the repeated DPYSPS motif, we hypothesized that P4EC10 could engage multiple copies of this motif within the extended Ara h 2 loop. To define the structural basis of this recognition, we determined a cryo-EM structure of Ara h 2 in complex with P4EC10 Fab fragments at 3.66 Å resolution. The P4EC10–Ara h 2 complex adopted two distinct binding stoichiometries, corresponding to 3:1 and 2:1 Fab:Ara h 2 complexes (**Fig. 3A, S4, S5, Table S2**). In the 3:1 state, the α-helical core of Ara h 2 adopts an upright orientation, with its principal axis approximately aligned with the threefold axis defined by the three bound Fabs. In contrast, in the 2:1 complex, the globular core of Ara h 2 lies approximately within the plane defined by the two bound Fabs (**Fig. 3A, B**). The 3:1 particles exhibited pronounced preferred orientation and were represented by a relatively small particle population, precluding reconstruction of this state at sufficient resolution for atomic model building. In contrast, the 2:1 complex could be reconstructed to 3.66 Å resolution, enabling determination of an atomic model (**Fig. 3C, D**).

**Figure 3.**
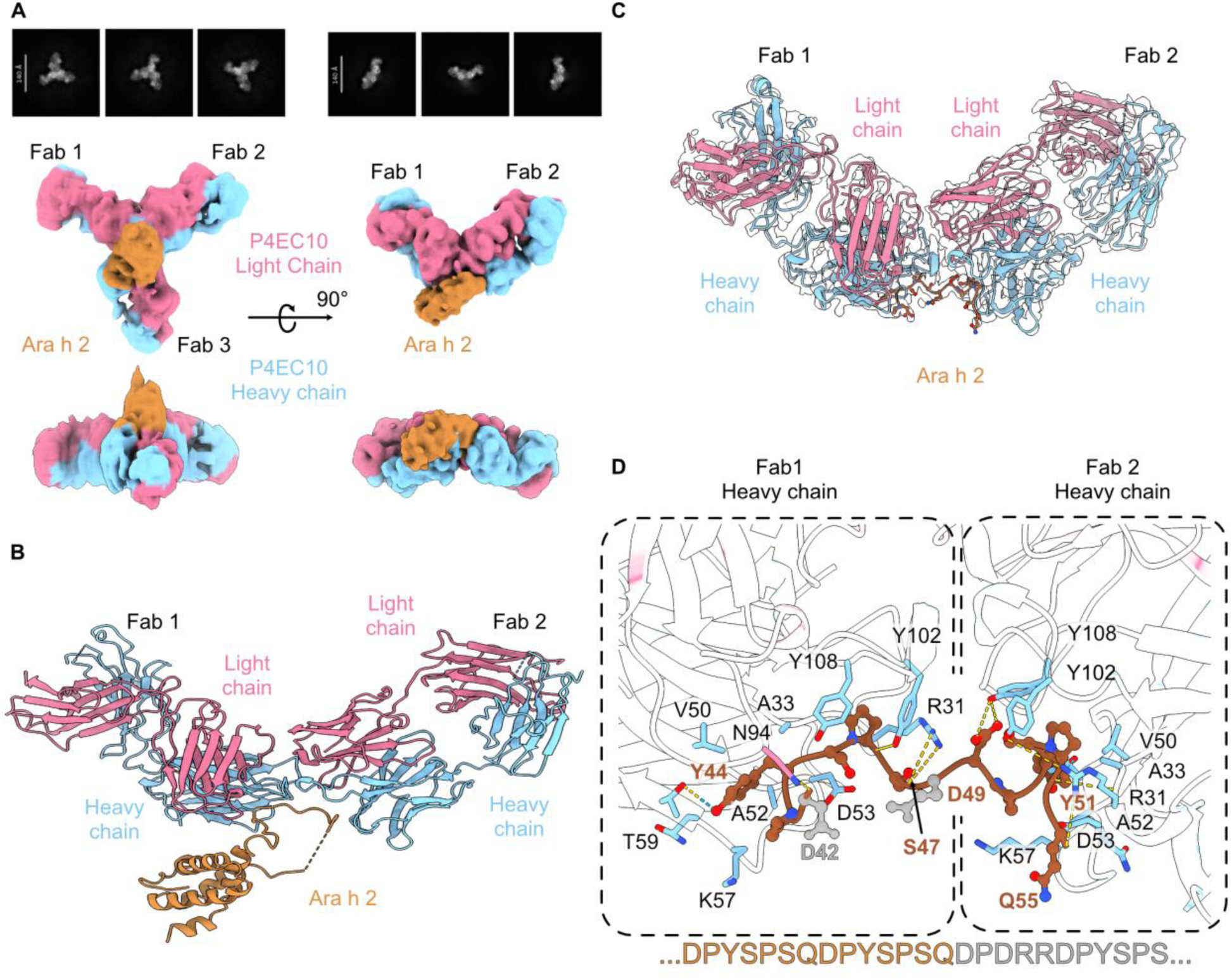
P4EC10 recognizes Ara h 2 through multivalent binding. **(A)** 2D class averages and overall cryo-EM maps for the P4EC10-Ara h 2 3:1 (left) and 2:1 (right) complexes. Ara h 2 is shown in light brown. The heavy and light chains of P4EC10 are shown in light blue and pink, respectively. (**B**) Model of the full Ara h 2 molecule bound with two P4EC10 Fabs. Model of the complete complex was generated by rigid-body fitting of the Ara h 2 structure from PDB ID 8SJ6 to the resolved DPYSPS repeat-containing loop, followed by flexible refinement of the remaining Ara h 2 structure. (**C**) Map-to-model fit. The atomic model of the ternary complex shown in cartoon representation and in the same orientation as in the 2:1 complex and colored as in (A). The Ara h 2 engaged loop peptide is shown in sticks. (**D**) Detailed view of the P4EC10-Ara h 2 interface with two Fabs engaging the DPYSPS-repeated epitope. Interacting residues are shown as sticks and hydrogen bonds and salt bridges indicated by yellow dashed lines. The amino acid sequence of the entire Ara h 2 loop is shown below, with the peptide residues resolved in cryo-EM colored brown.

In the resolved structure, two P4EC10 Fabs engage adjacent, structurally mirrored epitopes within the Ara h 2 loop, spanning residues 42–47 and 49–55. The cryo-EM reconstruction resolved the repeat-containing loop at sufficient resolution for atomic model building, whereas density corresponding to the remainder of Ara h 2 was poorly defined, likely reflecting substantial conformational flexibility. To approximate the position of the unresolved α-helical core, we generated a model of the complete complex by rigid-body fitting of the Ara h 2 structure from PDB ID 8SJ6 to the resolved DPYSPS repeat-containing loop, followed by flexible refinement of the remaining Ara h 2 structure (**Fig. 3D**). This model provides a plausible estimate of the position and orientation of the Ara h 2 α-helical core relative to the bound P4EC10 Fabs.

Within the resolved DPYSPS repeat-containing loop, each binding site corresponds to a copy of the DPYSPS repeat and is anchored at its N terminus by interactions between HCDR1 R31 and the Ara h 2 backbone. A hydrophobic pocket formed at the interface between the antibody heavy and light chains accommodates the central Tyr residue of each repeat, providing an additional conserved interaction (**Fig. 3D**). The heavy chain contributes most contacts with Ara h 2, although the light chain also participates in epitope recognition. Notably, LCDR3 N94 contacts the C-terminal region of the repeat motif in one Fab, whereas in the adjacent Fab it contributes to the interface between the two bound Fabs.

The two bound Fab molecules also form direct contacts through their CDR loops. Specifically, N94 and Y33 from one Fab form hydrogen-bonding interactions with D101 and Y102 in HCDR3 of the neighboring Fab (**Fig. S6**). Notably, N94 is somatically mutated rather than germline encoded, suggesting that affinity maturation introduced a residue that contributes both to antigen recognition and to stabilization of the inter-Fab interface.

Together, these structural observations demonstrate that a single Ara h 2 molecule can accommodate up to three P4EC10 Fabs through repeated recognition of the DPYSPS motif within its extended loop. The close spacing and continuity of these repeated epitopes, together with direct contacts between neighboring Fabs, provide a structural basis for direct contacts between adjacent repeat-bound Fabs. This unusual binding architecture provides a mechanistic explanation for the ability of a single IgE specificity to trigger FcεRI-mediated effector-cell activation *in vitro* and passive cutaneous anaphylaxis *in vivo* (**Fig. 1D,E**).

### Distinct Ara h 2 epitopes support simultaneous binding by P1EC5 and P1BC6

We next asked how Ara h 2 supports cooperative engagement by distinct antibody specificities. Because P1EC5 and P1BC6 required combined recognition to induce effector-cell activation, we examined whether Fabs from both antibodies could simultaneously engage a single Ara h 2 molecule. Cryo-EM analysis revealed that P1EC5 and P1BC6 bound simultaneously to a single Ara h 2 molecule, demonstrating that the two antibodies recognize spatially distinct, nonoverlapping epitopes on the allergen surface (**Fig. 4A,B, S7, Table S2**), consistent with our peptide-array and competition ELISA data (**Fig. 1B,C**). P1EC5 and P1BC6 engage distinct surfaces on the allergen formed by exposed helical and loop regions.

**Figure 4.**
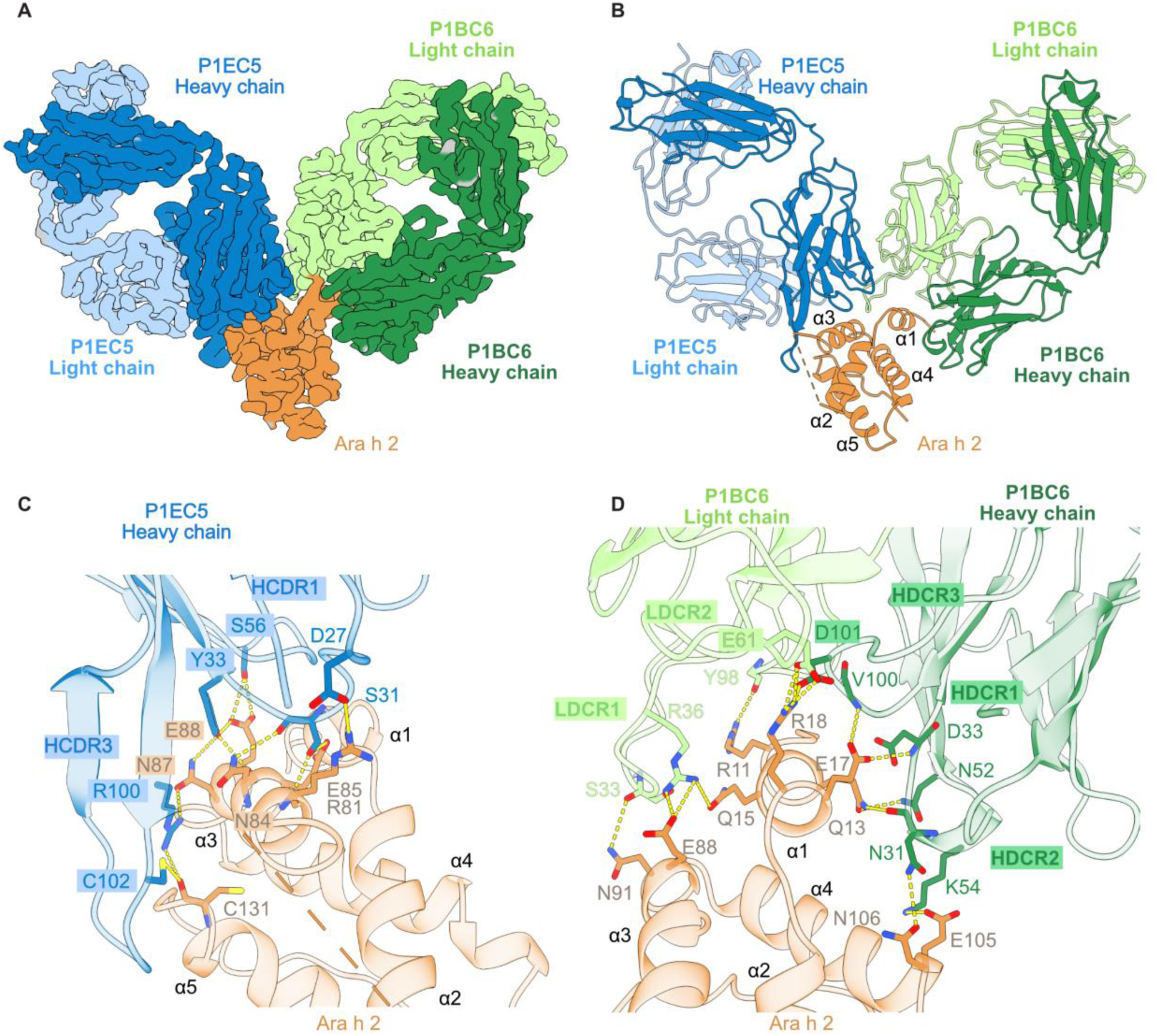
P1BC6 and P1EC5 simultaneously engage distinct, nonoverlapping epitopes on Ara h 2. **(A)** Overall cryo-EM map of the P1EC5–P1BC6–Ara h 2 ternary complex. Ara h 2 is shown in light brown. The heavy and light chains of P1EC5 are shown in dark and light blue, respectively, and those of P1BC6 in dark and light green, respectively. (**B**) The atomic model of the ternary complex shown in cartoon representation and in the same orientation and colored as in (A). Ara h 2 protein α-helices α1–α5 are indicated. (**C**) Detailed view of the P1EC5-Ara h 2 interface, with interacting residues shown as sticks and hydrogen bonds and salt bridges indicated by yellow dashed lines. Major interacting antibody CDR loops are labeled. Ara h 2 α-helices α1–α5 are indicated. (**D**) Detailed view of the P1BC6-Ara h 2 interface, with interacting residues shown as sticks and hydrogen bonds and salt bridges indicated by yellow dashed lines. Major interacting antibody CDR loops are labeled. Ara h 2 α-helices α1–α5 are indicated.

P1EC5 binds the α3-centered surface that overlaps the 13T5 footprint in bin 2 from Min et al.^21^ (**Fig. 4C, S4, S10A**). Its footprint includes Ara h 2 residues R81, N84, E85, N87, E88, C131 with most contacts concentrated around the helix α3 surface. The P1EC5 heavy chain dominates the interaction with Ara h 2. Heavy-chain residues D27, S31, Y33, S56, R100, and C102 contribute to an extended interface comprising polar interactions and close contacts with charged and polar residues on the allergen surface.

P1BC6 recognizes a distinct composite epitope spanning the N-terminal region and additional structural elements of Ara h 2 (**Fig. 4D**). Its footprint includes Ara h 2 residues R11, Q13, Q15, E17, R18, E88, N91, E105, and N106. In contrast to P1EC5, both the heavy and light chains of P1BC6 contribute substantially to the interface. Light-chain residues S33, R36, E61, and Y98 and heavy-chain residues N31, D33, K54, V100, and D101 contact Ara h 2 through a combination of polar interactions and close packing contacts. These contacts create a composite interface that extends across multiple regions of the allergen surface.

The P1EC5–P1BC6 structure demonstrates simultaneous Fab occupancy of distinct surfaces on one Ara h 2 molecule. The two approaches are compatible with co-binding in the soluble complex and provide structural context for activation by the corresponding recombinant IgE pair. The structure does not, however, resolve intact receptor-bound IgE or establish how approach angle affects signaling. P1EC5 primarily targets the helix α3 (**Fig. S4**), whereas P1BC6 engages a separate composite epitope spanning multiple Ara h 2 structural elements. Their simultaneous engagement of Ara h 2 provides a structural basis for the cooperative activity observed in our functional assays: neither antibody alone efficiently induced FcεRI-mediated effector-cell activation *in vitro* or passive cutaneous anaphylaxis *in vivo*, whereas their combination triggered robust allergen-dependent effector cell activation *in vitro* and *in vivo*. Thus, the structure directly links the spatial organization of distinct Ara h 2 epitopes to the capacity of different IgE specificities to cooperate in FcεRI crosslinking and allergic effector responses. These findings establish a molecular framework for understanding how the epitope architecture of Ara h 2 governs cooperative IgE-mediated allergen recognition and activation.

### Isothermal titration calorimetry (ITC) supports distinct Fab binding stoichiometries

The interactions between Ara h 2-specific mAbs and their respective epitopes were quantified using isothermal titration calorimetry (ITC). P4EC10 binds the repeat-containing loop of Ara h 2, and we therefore used ITC to determine the affinity and binding stoichiometry of P4EC10 for hydroxylated and non-hydroxylated loop peptides in solution. P4EC10 bound both peptides, with apparent dissociation constants (K_D_) of 34 nM for the non-hydroxylated peptide and 1 nM for the hydroxyproline-containing peptide (**Fig. 5A, B, S8A,B**). The corresponding Fab stoichiometries were 2.5 and 2.6, respectively. These values indicate that P4EC10 engages multiple repeat units within the Ara h 2 loop, with an apparent stoichiometry intermediate between two and three Fabs per loop, consistent with the multivalent binding mode observed by cryo-EM.

**Figure 5.**
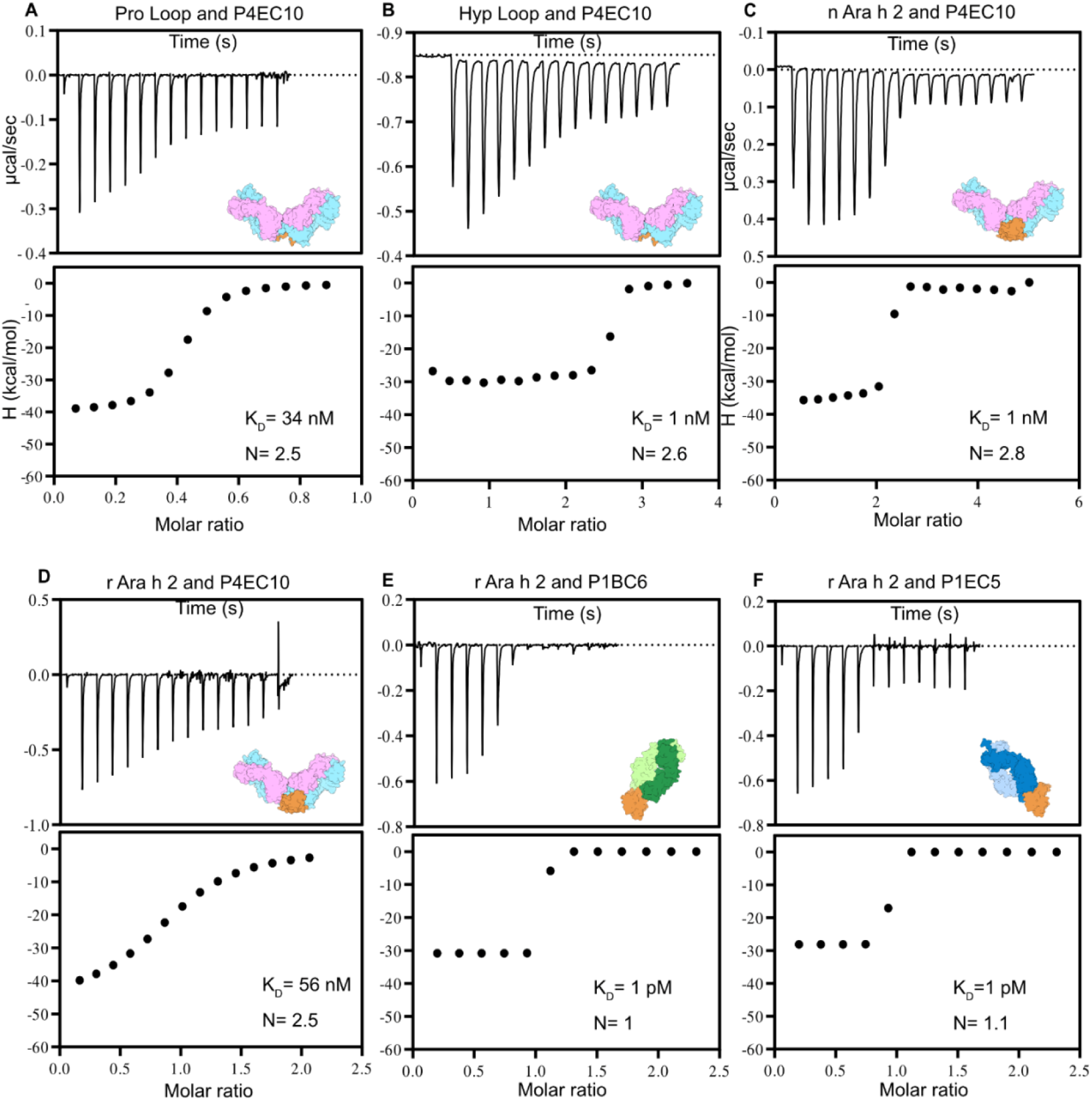
P1BC6, P1EC5, and P4EC10 bind Ara h 2 with distinct stoichiometries and affinities. Isothermal titration calorimetry (ITC) measurements of Ara h 2-specific Fab binding. Representative thermograms are shown above the corresponding isotherms (normalized heat per injection plotted as a function of molar ratio). (**A**) P4EC10 binding to the non-hydroxylated (Pro) Ara h 2 repeat-loop peptide. (**B**) P4EC10 binding to the hydroxyproline-containing (Hyp) repeat-loop peptide. (**C**) P4EC10 binding to native Ara h 2 (nAra h 2). (**D**) P4EC10 binding to recombinant Ara h 2 (rAra h 2). (**E**) P1BC6 binding to rAra h 2. (**F**) P1EC5 binding to rAra h 2. Dissociation constants calculated according to a simple independent binding site model are displayed.

We next measured P4EC10 binding to native Ara h 2 (nAra h 2) and recombinant Ara h 2 (rAra h 2). P4EC10 bound nAra h 2 with an apparent K_D_ of 1 nM and a stoichiometry of 2.8 (**Fig. 5C**, **S8D**). In contrast, P4EC10 bound rAra h 2 with an apparent K_D_ of 56 nM and a Fab:Ara h 2 stoichiometry of 2.5 (**Fig. 5D**, **S8C**). The binding stoichiometries measured for the non-hydroxylated loop peptide, hydroxyproline-containing loop peptide, rAra h 2, and nAra h 2 were highly consistent, ranging from 2.5 to 2.8 Fabs per antigen. These values support a common binding model in which P4EC10 engages two to three copies of the repeated DPYSPS-containing motif within a single Ara h 2 molecule.

Because our ITC measurements suggested that proline hydroxylation within the Ara h 2 repeat-containing loop may contribute to P4EC10 binding, we modeled hydroxyproline residues into the cryo-EM structure (**Fig. S9**). Substitution of the corresponding prolines with hydroxyprolines introduced three additional potential hydrogen bonds at the antibody-allergen interface (**Fig. S9**). The hydroxyl group at Ara h 2 residue 46 formed potential hydrogen-bonding interactions with P4EC10 residues T99 and Y108 in Fab1, whereas the hydroxyl group at residue 53 formed an additional potential interaction with T99. These modeled contacts provide a structural rationale for the higher apparent affinity of P4EC10 for the hydroxyproline-containing peptide as well as nAra h 2 observed by ITC.

We also characterized the binding of P1BC6 and P1EC5 to rAra h 2. Both antibodies exhibited very high-affinity binding, with fitted K_D_ values of approximately 1 pM (**Fig. 5E,F, S8E,F**). The approximately 1 pM fitted dissociation constants are not interpreted as reliable affinity measurements because the titrations do not adequately resolve binding in this range. The corresponding Fab h 2 stoichiometries were 1.0 for P1BC6 and 1.1 for P1EC5, consistent with one binding site per antibody on each Ara h 2 molecule.

Together, these ITC measurements reveal distinct binding profiles among the Ara h 2-specific antibodies. P4EC10 exhibits multivalent binding, with stoichiometries of approximately 2.5-2.8 across loop peptides and intact Ara h 2 and shows higher apparent affinity for hydroxylated and native forms of the allergen. In contrast, P1BC6 and P1EC5 bind rAra h 2 with sub-nanomolar affinity and near 1:1 stoichiometry. These solution-based measurements are consistent with the binding architectures observed by cryo-EM.

### Ara h 2 accommodates both convergent and distinct modes of antibody recognition

We compared the footprints of P1EC5, P1BC6, and P4EC10 with those of previously characterized Ara h 2-specific antibodies to assess the extent of epitope overlap and structural convergence (**Fig. S10**). P1EC5 showed substantial overlap with the footprints of 13T5^7^ and, to a lesser extent with the Ara h 6-specific antibody 8F3^20^, with α2 and α3 forming the core of the shared epitope. The extensive overlap between P1EC5 and 13T5 is consistent with convergent recognition of the same conformational surface on the folded Ara h 2 core (**Fig. S4, S10A**). P1BC6 which primarily targets the N-terminal α1 helix, has an extended footprint toward α4 and partially overlapped with that of 13T1^7^ (**Fig. S4, S10B**).

In contrast, P4EC10 recognized a distinct epitope within the flexible, repeat-containing loop. Among previously structurally characterized antibodies, only 38B7^20^ also targets the DPYSPS repeat. The binding of one 38B7 Fab to Ara h 2 was characterized by crystallography^20^. Despite the shared specificity, the two antibodies engage the loop differently, with P4EC10 binding exclusively to the repeat motif and Ara h 2 accommodating up to three P4EC10 paratopes (**Fig. 3C, D**), whereas 38B7 Fab binds to one repeat and additionally contacts the adjacent structured region (**Fig. S10C, D**). Indeed, in the crystal structure reported by Smith *et al.*, one 38B7 Fab bound a DPYSPS repeat and was positioned close enough to Arg102 on helix 5 to suggest an additional contact (**Fig. S10D**). Thus, while Smith *et al*. resolved one 38B7 Fab bound to recombinant Ara h 2, they proposed that the repeated, hydroxylated DPYSPS motifs in native Ara h 2 allow a single allergen molecule to crosslink receptor-bound 38B7 IgE, accounting for the anaphylaxis observed in mice sensitized with 38B7 IgE alone^17^.

Overall, Ara h 2 presents a highly immunogenic surface and contains both recurrent antigenic domains, targeted through convergent binding modes, and regions for which distinct antibody solutions can be accommodated.

## DISCUSSION

We previously identified a population of CD23^+^IgG1^+^ memory B cells in children with peanut allergy that includes high-affinity Ara h 2-specific clones and is poised to generate pathogenic IgE^18^. Within this population, we noted Ara h 2-specific clones with B-cell receptor sequences and specificities shared across individuals, suggesting that recurrent solutions to Ara h 2 recognition are maintained within the memory B-cell compartment. Here, we examined three Ara h 2-specific antibodies from this population. When expressed as IgE, P4EC10 alone supported Ara h 2-induced degranulation, whereas P1BC6 and P1EC5 were active only when combined. These experiments establish the effector activity that the three variable-region pairs can confer when expressed as IgE. Whether the corresponding clones produced IgE in the children from whom they were isolated remains unknown, however P4EC10 belongs to a public clone with IgE members^15,18^. To our knowledge, these are the first cryo-EM structures of Ara h 2 in complex with human antibody Fabs. They show how two P4EC10 Fabs contact one another on adjacent copies of a linear DPYSPS epitope and how P1BC6 and P1EC5 simultaneously engage distinct conformational epitopes on the same allergen molecule.

Previous studies showed that a single IgE specificity directed to the DPYSPS repeats of Ara h 2 can trigger mast-cell degranulation and anaphylaxis^16,17^. Parkkinen *et al.* resolved, by crystallography, three D08 Fabs bound to the repeat-containing loop and showed that neighboring Fabs contact one another^22^. We observed both 2:1 and 3:1 P4EC10 Fab–Ara h 2 complexes by cryo-EM, consistent with the stoichiometries of 2.5–2.8 Fabs per antigen measured by ITC. In the 2:1 structure, two P4EC10 Fabs bind adjacent DPYSPS motifs and contact one another through their CDR loops. Of particular interest is the somatically mutated LCDR3 residue N94. In one Fab, N94 contacts the C-terminal end of the DPYSPS motif, whereas in the adjacent Fab, the same residue contributes to the Fab-Fab interface. Somatic mutations can stabilize contacts between neighboring antibodies bound to repeats in the *Plasmodium falciparum* circumsporozoite protein^23,24^. Whether N94 similarly stabilizes adjacent P4EC10 Fabs on Ara h 2 remains an open question.

Proline hydroxylation increased the apparent affinity of P4EC10 for the repeat loop. P4EC10 bound the hydroxylated peptide with an apparent K_D_ of 1 nM and the unmodified peptide with an apparent K_D_ of 34 nM, with similar Fab stoichiometries for the two peptides. P4EC10 also bound native Ara h 2 more tightly than recombinant Ara h 2. Modeling hydroxyproline at positions 46 and 53 suggested additional hydrogen bonds at the antibody–allergen interface that could account for the peptide affinity difference. The magnitude of this effect varies among repeat-directed antibodies. Whereas 38B7 binding depends strongly on hydroxylation of the second proline in the DPYSPS motif,^20^ P4EC10 retained nanomolar affinity for the unmodified peptide, and both peptides triggered degranulation of P4EC10 IgE-sensitized cells.

P1BC6 and P1EC5 reveal a second route to FcεRI crosslinking. Both antibodies bind Ara h 2 with very high affinity and 1:1 stoichiometry, yet neither efficiently activated effector cells when tested individually. Their lack of individual activity therefore separates binding affinity from pathogenic potency. Cryo-EM showed that the two Fabs can bind simultaneously to spatially distinct, nonoverlapping surfaces of a single Ara h 2 molecule. When expressed as IgE and combined, P1BC6 and P1EC5 induced robust degranulation *in vitro* and passive cutaneous anaphylaxis *in vivo*. These observations provide a direct structural link between multispecific recognition of an allergen and effector-cell activation: a single Ara h 2 molecule can serve as a bridge between FcεRI-bound IgEs of different specificities even when neither specificity encounters a repeated epitope. Previous structural studies have mapped several distinct conformational antibody epitopes across the compact Ara h 2 surface^7,20,25^, highlighting the antigenic density of this small allergen. Our findings demonstrate the functional consequence of that density by showing that simultaneous occupation of separate epitopes can generate an activating antibody arrangement. The 2025 study by Marini-Rapoport *et al.* further showed that structurally convergent recognition of Ara h 2 can arise from multiple germline rearrangements, emphasizing that stereotyped antibody solutions to this allergen can be widespread in humans^25^.

Keswani *et al.* showed that inhibition of serum IgE binding to Ara h 2 varied with the pair of conformational epitopes targeted^26^. Here, we compared antibodies to two conformational epitopes with an antibody to the linear DPYSPS repeat. Across the 24 plasma samples stratified by Ara h 2-specific IgE, the three IgG1 antibodies together inhibited a mean of 77–80% of Ara h 2 binding to plasma IgE in each of the four Ara h 2-specific IgE strata. The individual antibodies differed in their inhibition profiles. P4EC10 inhibited a mean of 46% of binding in the highest stratum and 24% in the lowest. P1EC5 showed the opposite pattern (6% and 26%, respectively), whereas P1BC6 inhibited approximately 15% in both groups. An intact IgG1 bound to Ara h 2 can obstruct IgE access to sites adjacent to its footprint. The fraction of binding inhibited by P4EC10 therefore cannot be equated with the fraction of plasma IgE directed to the DPYSPS repeats. The comparison across strata also warrants caution because each group contained six individuals and the plasma dilutions differed among groups.

Previous work examined how Ara h 2-specific IgG antibodies inhibit both plasma IgE binding and allergen-induced degranulation^27^. In our experiments, P4EC10 IgG1 alone had little effect on Ara h 2-induced degranulation of RBL-SX38 cells sensitized with pooled plasma from three donors with high Ara h 2-specific IgE. At the same total IgG1 concentration, pairing P4EC10 with either P1BC6 or P1EC5 substantially inhibited degranulation in response to both purified Ara h 2 and peanut extract. The P1BC6–P1EC5 pair was less effective, and adding the third antibody gave little further inhibition over either P4EC10-containing pair. Thus, although the repeat loop alone could support activation of cells sensitized with P4EC10 IgE, inhibition in the plasma-sensitized cells was greatest when the antibodies targeted both the loop and a second surface on Ara h 2.

The cryo-EM structures resolve the contacts that allow two Fabs to bind simultaneously to Ara h 2. The number and orientation of intact, receptor-bound IgE molecules in the signaling complex remain unknown. Similarly, although the three antibodies supported effector activity when expressed as IgE, we isolated their variable regions from IgG1 memory B cells. Whether these clones contributed to circulating IgE in the donors is unknown. We measured inhibition of IgE binding in 24 individual plasma samples and tested inhibition of degranulation using a single pool from three donors. Pooling therefore precluded a donor-by-donor comparison of the two readouts. In this pool, P4EC10 illustrates why the functional assay matters when selecting blocking antibodies: its activity in the binding assay gave little indication of the gain in inhibition achieved by adding a second specificity.

## MATERIALS AND METHODS

### Selection and production of Ara h 2-specific monoclonal antibodies

Ara h 2-specific monoclonal antibodies P4EC10, P1EC5, and P1BC6 were selected from IgG1 memory B-cell clones previously identified in peanut-allergic individuals, as described by Ota et al. (24). P4EC10 and P1BC6 belong to the PA13P1E10-related and 4C5G1 convergent BCR families, respectively. Paired heavy- and light-chain variable-region sequences were cloned into expression vectors containing human IgG1 or IgE heavy-chain constant regions and the corresponding κ or λ light-chain constant regions. Recombinant full-length IgG1 and IgE mAbs were produced by Expi293 cells and purified using Protein A or IMAC affinity resins, respectively. Antibody concentrations were determined by absorbance at 280nm. Fab fragments used for structural studies were produced as described below.

### Cloning, expression, and purification of IgE antibodies

The genes for P1EC5 kappa light chain and the soluble isoform of the IgE P1EC5 heavy chain were introduced into pTRIOZ vector via restriction site pairs PmeI-BsIWI and AgeI-AvrII, respectively. The construct encoding the membrane isoform of IgE P1EC5 was modified to encode the his-tagged soluble isoform by inverse PCR. Parallel methods were used to generate to construct encoding IgE P4EC10. The construct encoding IgE P1BC6 was generated by restriction cloning of the oligonucleotide encoding IgE P1BC6 heavy chain (TwistBio) and light chain (digested from preexisting construct) into pTRIOZ vector via restriction site pairs AgeI-AvrII and PmeI-BsIWI, respectively.

pTRIOZ constructs encoding IgE clones were linearized by digestion with PacI restriction endonuclease (NEB). The linearized plasmid was purified using Quiaquick TM PCR purification kit (Quiagen), according to manufacturer protocol. Linearized DNA encoding the IgE clones was transfected in complex with PEI (1:3 ratio; linear 25kDa PEI; Polysciences) into Expi293 cells at a rate of 1 ug of DNA per mL of transfected culture. The resultant transfectants were cultured for 48 hours, prior to commencing selection under 50 ug/mL zeocin. Zeocin selection at 50 ug/mL was performed for 4 weeks. The resultant stable transfectants were further selected at elevated zeocin concentration (250 ug/mL) for 2 weeks.

His-tagged IgE was purified from stable transfectant supernatant by a standard affinity chromatography pipeline. Briefly, clarified supernatant was applied to a gravity column containing 5 mL of Talon TM cobalt resin equilibrated in buffer consisting of 20 mM HEPES (pH 7.5) and 150 mM NaCl (‘binding buffer’). Resin was washed with 6 column volumes of binding buffer, followed by 6 columns of wash buffer (20 mM HEPES, 500 mM NaCl, 5 mM imidazole). His-tagged IgE was eluted in buffer consisting of 20 mM HEPES (pH 7.5), 150 mM NaCl, and 400 mM imidazole. Crude eluate was concentrated in a 50 kDa MWCO PES centrifugal device (Amicon) and further purified by size exclusion chromatography on a Superdex200 (Cytiva) column equilibrated in binding buffer.

### Cloning, expression, and purification of Ara h 2 and Fabs

Recombinant Ara h 2.0201 and the P4EC10, P1BC6, and P1EC5 Fabs were expressed in Expi293 cells (Thermo Fisher Scientific, A14527). Protein constructs were cloned into the pVRC mammalian expression vector. Five days after transient transfection, culture supernatants were collected, clarified by centrifugation at 3,500 × g for 30 min, and filtered through a 0.45-µm filter. Recombinant proteins were purified from the filtered supernatants using 5 mL of TALON® metal affinity resin (Takara Bio, 635504) and eluted as previously described. Eluted proteins were concentrated using 10-kDa Vivaspin® 20 centrifugal concentrators (Sartorius, VS2002) and further purified by size-exclusion chromatography using an ÄKTA system for size separation and buffer exchange.

### Allergen-binding ELISA

High-binding 96-well Immulon 4 HBX plates (Thermo Scientific, cat. no. 3855) were coated overnight at 4°C with peanut extract (Stallergenes Greer, item no. XPF171D3A-25; lot 303179), cashew extract (prepared in-house), natural Ara h 1, Ara h 2, and Ara h 6 (AllerTeQ; batches 2014-01, 2012014-01, and 2014-02, respectively), natural Ara h 3 (InBio, cat. no. NA-AH3-1; lot 45129), recombinant Ara h 2 (produced as described above), or non-hydroxylated and hydroxyproline-containing synthetic Ara h 2 loop-derived peptides (Peptide 2.0), at 2 μg/mL in DPBS (Gibco, cat. no. 14190-144). Plates were washed with PBS containing 0.05% Tween-20 (PBST) and blocked with 1% BSA in PBST for 1 h at room temperature. Recombinant human IgE mAbs were added at 10, 30, or 90 ng/mL in 1% BSA/PBST and incubated overnight at 4°C. After washing, bound IgE was detected with biotinylated anti-human IgE mAb 182 (MABTECH, cat. no. 3810-6; 1:1500) for 1 h at room temperature, followed by streptavidin-HRP (BD Pharmingen, cat. no. 554066; 1:2000) for 30 min. Plates were developed with TMB substrate (BioLegend, cat. no. 421101), the reaction was stopped with 0.5 M H₂SO₄ (LabChem, cat. no. LC257704), and absorbance was measured at 450 nm using an EnVision 2105 Multimode Plate Reader (PerkinElmer).

### Bead-based linear epitope mapping

Linear epitope recognition by recombinant human IgE mAbs was assessed using a previously described bead-based epitope assay^28^. Peptides representing sequential epitopes of Ara h 1, Ara h 2 and Ara h 3 were coupled to LumAvidin xMAP microspheres (Luminex Corporation). Recombinant IgE mAbs were added to the master mix (MM) of Ara h 1, Ara h 2 and Ara h 3 peptides at 0.1 ng/μL and incubated for 2 h at room temperature with shaking at 400 rpm. After washing, bound IgE was detected with PE-conjugated mouse anti-human IgE (Thermo Fisher Scientific, cat. no. MA1-10375; 2 μg/mL) for 30 min at room temperature. Median fluorescence intensity (MFI) was measured using a Luminex 200 instrument (Luminex Corporation) and quantified with the xPONENT software. Bead-based epitope assay buffer-only wells were included to assess nonspecific fluorescence.

### Competitive Ara h 2-binding ELISA

High-binding 96-well Immulon 4 HBX plates (Thermo Scientific, cat. no. 3855) were coated overnight at 4°C with recombinant human IgE mAbs P4EC10, P1EC5, or P1BC6 (1 μg/mL; 50 μL/well) in DPBS. Plates were washed with PBST and blocked with 2% BSA in PBST for 2 h at room temperature. In a separate plate, biotinylated natural Ara h 2 was mixed 1:1 with serial dilutions of P4EC10, P1EC5, P1BC6, or an irrelevant human mAb to final concentrations of 0.03 μg/mL Ara h 2 and 0.1875–6 μg/mL competing mAb and preincubated for 2 h at room temperature. Allergen–antibody mixtures were transferred to the IgE-coated plates and incubated for 2 h at room temperature. After washing, bound biotinylated Ara h 2 was detected with streptavidin-HRP (BD Pharmingen, cat. no. 554066; 1:2000) for 30 min. Plates were developed with TMB substrate (BioLegend, cat. no. 421101), the reaction was stopped with 0.5 M H₂SO₄ (LabChem, cat. no. LC257704), and absorbance was measured at 450 nm using an EnVision 2105 Multimode Plate Reader (PerkinElmer). Relative Ara h 2 binding was calculated after background subtraction and expressed relative to the no-competition control, set to 100%.

### RBL-SX38 cell culture

RBL-SX38 rat basophilic leukemia cells expressing the human FcεRI α-chain were maintained in RPMI 1640 medium supplemented with GlutaMAX (Gibco, cat. no. 61870-036), 10% heat-inactivated fetal bovine serum (HyClone, cat. no. SH30071.03), 1% penicillin-streptomycin (Gibco, cat. no. 15140-122), 1 mM sodium pyruvate (Gibco, cat. no. 11360-070), 1× MEM non-essential amino acids (Gibco, cat. no. 11140-050), 15 mM HEPES, and 1.2 mg/mL Geneticin (G418; Gibco, cat. no. 10131-035). Cells were maintained at 37°C in a humidified atmosphere with 5% CO₂ and passaged approximately every 2 days at 80–90% confluence using TrypLE Express (Gibco, cat. no. 12604-013). Cells between passages 6 and 12 were used for degranulation experiments.

### IgE sensitization and degranulation assay

RBL-SX38 cells were seeded in flat-bottom 96-well plates at 5 × 10⁴ cells/well in 100 μL of complete medium without Geneticin and incubated overnight at 37°C and 5% CO₂. The following day, the medium was replaced with sensitization medium containing 0.5% FBS and recombinant human IgE mAbs P4EC10, P1EC5, and P1BC6, alone or in the indicated combinations, at a total IgE concentration of 0.5 μg/mL. Cells were incubated overnight at 37°C and 5% CO₂. For mAb combinations, each antibody contributed equally to the total IgE concentration. Pooled plasma from peanut-allergic (PA) and non-peanut-allergic (non-PA) individuals (Table S1) was included at a 1:50 dilution as controls. After sensitization, cells were washed twice with assay buffer (HBSS containing Ca²⁺ and Mg²⁺ supplemented with 0.5% BSA) and challenged for 1 h at 37°C with peanut extract, natural Ara h 1, Ara h 2, Ara h 3, or Ara h 6, recombinant Ara h 2.0201, or hydroxylated and non-hydroxylated synthetic Ara h 2 loop-derived peptides, each at 0.5 μg/mL in 100 μL of assay buffer. Rabbit anti-human IgE (Bethyl Laboratories/Fortis Life Sciences, cat. no. A80-109A; 0.5 μg/mL) was used as a positive control. Degranulation was quantified by β-hexosaminidase release. Culture supernatants (50 μL) were incubated with an equal volume of p-nitrophenyl-N-acetyl-β-D-glucosaminide (Sigma-Aldrich, cat. no. N9376) in 0.2 M citrate buffer (pH 4.5) for 1 h at 37°C. Reactions were stopped with 100 μL of 0.2 M glycine buffer (pH 10.7), and absorbance was measured at 450 nm using an EnVision 2105 Multimode Plate Reader (PerkinElmer).

### Passive cutaneous anaphylaxis

Human FcεRIα-transgenic mice (B6.Cg-*Fcer1a^tm1Knt^* Tg(FCER1A)1Bhk/J; JAX #010506) were used for passive cutaneous anaphylaxis (PCA) experiments. Mice were maintained under specific pathogen-free conditions at the Icahn School of Medicine at Mount Sinai, and all procedures were performed in accordance with NIH guidelines and protocols approved by the Institutional Animal Care and Use Committee (IACUC protocol 2019-0040). For PCA, one ear per mouse was sensitized intradermally with recombinant human IgE mAbs P4EC10, P1EC5, and P1BC6 (0.5 μg total/ear in 10 μL), alone or in the indicated combinations, or with pooled peanut-allergic plasma (Table S1). PBS was used as a negative control. After 12 h, mice were challenged intravenously with 40 μg of natural or recombinant Ara h 2, as indicated, in 80 μL containing 1% Evans blue. Thirty minutes after challenge, mice were euthanized, and sensitized ears were collected, minced, and incubated individually in 500 μL of formamide (Fisher Scientific, cat. no. BP227-100) at 65°C with shaking. Evans blue extravasation was quantified by measuring absorbance at 620 nm.

### Plasma IgE inhibition assay

High-binding 96-well Immulon 4 HBX plates (Thermo Scientific, cat. no. 3855) were coated overnight at 4°C with anti-human IgE mAb 107 (code 3810-3; 1:500 in PBS). Plates were washed with PBST and blocked with 2% BSA in PBST for 2 h at room temperature. Plasma samples from peanut-allergic individuals were diluted according to Ara h 2 sIgE levels (>100 kUA/L, 1:250; 40–60 kUA/L, 1:25; 10–20 kUA/L, 1:10; and 4–8 kUA/L, 1:5) and incubated overnight at 4°C. Biotinylated natural Ara h 2 (0.03 μg/mL final concentration) was preincubated for 2 h at room temperature with recombinant human IgG1 mAbs P4EC10, P1EC5, and P1BC6, alone or in the indicated combinations, at a total IgG1 concentration of 3 μg/mL. For antibody combinations, the total IgG1 concentration was divided equally among the component antibodies. The Ara h 2–IgG1 mixtures were then added to the plasma IgE-captured wells and bound biotinylated Ara h 2 was detected with streptavidin-HRP and developed as described above. Percent inhibition was calculated after background subtraction relative to Ara h 2 binding in the absence of competing IgG1. Plasma samples from 24 peanut-allergic individuals were stratified according to Ara h 2 sIgE levels (>100, 40–60, 10–20, and 4–8 kUA/L; n = 6 per group).

### IgG-mediated inhibition of degranulation

RBL-SX38 cells were sensitized overnight with pooled plasma from three peanut-allergic individuals (Table S1) diluted 1:100. Peanut extract or natural Ara h 2 was preincubated for 40 min at 37°C with recombinant human IgG1 mAbs P4EC10, P1EC5, and P1BC6, alone or in the indicated combinations. Sensitized cells were challenged with the allergen–IgG1 mixtures at final concentrations of 0.1 μg/mL allergen and 0.1875–6 μg/mL total IgG1. For mAb combinations, each antibody contributed equally to the total IgG1 concentration. An irrelevant human IgG mAb was included as a control. Degranulation was quantified by β-hexosaminidase release as described above. Percent inhibition was calculated after background subtraction relative to allergen-induced degranulation in the absence of IgG1.

### Statistical analysis

Statistical analyses were performed using GraphPad Prism version 11.0.0 (GraphPad Software). Data are presented as mean ± SD or SEM, as indicated in the figure legends. Statistical significance was assessed using one-way or two-way analysis of variance (ANOVA), as appropriate, followed by Tukey’s or Šídák’s multiple-comparisons test as specified in the figure legends.. Technical replicates were averaged before statistical analysis. P values < 0.05 were considered statistically significant.

### Cryo-EM sample preparation

For preparation of the Ara h 2.0201-P4EC10 complex, Ara h 2.0201 was incubated with P4EC10 Fab and CaptureSelect™ LC-kappa (Human) Affinity Ligand (Thermo Fisher Scientific, 1033270500) at a final protein concentration of 1 mg/mL, using a 1.2-fold molar excess of both Fab and anti-kappa nanobody relative to Ara h 2.0201. The mixture was incubated for 20 min at room temperature, and the resulting complex was isolated by size-exclusion chromatography using an ÄKTA system.

The purified Ara h 2.0201–P4EC10 complex was applied to UltrAuFoil R1.2/1.3 gold grids (Ted Pella Inc., 688-300-AU). Before sample application, grids were glow-discharged using a PELCO easiGlow device at an air pressure of 0.39 mbar and a current of 28 mA for 25 s. Immediately before grid preparation, amphipol (Anatrace, A835) was added to the preformed complex to a final concentration of 0.07% (w/v). A 3-µL aliquot of the complex was applied to each grid, blotted for 3.5 s at 10°C and 100% humidity, and plunge-frozen in liquid ethane using a Leica EM GP.

For preparation of the Ara h 2.0201-P1BC6-P1EC5 complex, Ara h 2.0201 was expressed as an N-terminal fusion with the BRIL domain and purified as described above. Ara h 2.0201–BRIL was incubated with P1BC6 and P1EC5 Fabs and CaptureSelect™ LC-kappa (Human) Affinity Ligand (Thermo Fisher Scientific, 1033270500) at a final protein concentration of 2.3 mg/mL. Each Fab and the anti-kappa nanobody were added at a 1.2-fold molar excess relative to Ara h 2.0201. The mixture was incubated for 20 min at room temperature, and the resulting complex was isolated by size-exclusion chromatography using an ÄKTA system.

The purified Ara h 2.0201-P1BC6-P1EC5 complex was applied to Ultrathin Carbon on Quantifoil® R1.2/1.3 orthogonal-array grids (Ted Pella Inc.). Before sample application, grids were glow-discharged using a PELCO easiGlow device at an air pressure of 0.39 mbar and a current of 28 mA for 25 s. Immediately before grid preparation, n-octyl-β-D-glucoside (Anatrace) was added to the preformed complex to a final concentration of 0.1% (w/v). A 3-µL aliquot of the complex was applied to each grid, blotted for 5.5 s at 10°C and 100% humidity, and plunge-frozen in liquid ethane using a Vitrobot Mark IV (Thermo Fisher Scientific).

### Cryo-EM Data Processing

#### P4EC10 complex

A total of 11,546 movies were aligned and dose-weighted using Patch Motion Correction implemented in cryoSPARC v5.0.3^29^. Contrast transfer function (CTF) parameters were estimated using Patch CTF, and particles were identified using the cryoSPARC blob picker followed by template picker. Particles were extracted and subjected to two successive rounds of 2D classification, yielding 480,718 particles for initial model generation. Following initial model generation, 147,181 particles corresponding to the Ara h 2–P4EC10 complex were selected and subjected to heterogeneous refinement, 3D classification, and ab initio reconstruction with three classes (K = 3). A selected class containing 88,413 particles was used for subsequent processing.

Particles from this class were used to train Topaz^30^ for particle picking and extraction. The resulting particles were subjected to 2D classification, ab initio reconstruction (K = 3), homogeneous refinement, 3D classification, a second round of homogeneous refinement, and non-uniform (NU) refinement. Particles were subsequently re-extracted and subjected to an additional round of NU-refinement followed by ab initio reconstruction with ten classes (K = 10). A selected particle subset containing 13,494 particles was then used for an additional round of Topaz training and particle extraction.

Topaz-picked particles were subjected to two rounds of 2D classification followed by ab initio reconstruction with three classes (K = 3), yielding a selected population of 113,815 particles. These particles were further classified by ab initio reconstruction with two classes (K = 2). The selected particles were subjected sequentially to local refinement, reference-based motion correction, local refinement, local CTF refinement, and a final round of local refinement. The final reconstruction contained 62,331 particles and reached a nominal global resolution of 3.66 Å, as determined using the gold-standard Fourier shell correlation (FSC) 0.143 criterion.

#### P1BC6 and P1EC5 complex

A total of 6,700 movies were aligned and dose-weighted using Patch Motion Correction implemented in cryoSPARC v5.0.3^29^. Contrast transfer function (CTF) parameters were estimated using Patch CTF, and particles were identified using the cryoSPARC template picker. Particles were extracted and subjected to two successive rounds of 2D classification, yielding 245,884 particles for initial model generation. Three classes containing 66,446, 72,580, and 76,880 particles, respectively, were selected for further processing.

The selected particles were subjected to heterogeneous refinement, non-uniform (NU) refinement, particle re-extraction, a second round of NU-refinement, and 3D classification. Classes corresponding to the intact Ara h 2–P1EC5–P1BC6 complex were combined, yielding 213,308 particles, and subjected to NU-refinement. Particles were subsequently re-extracted and further refined by sequential NU-refinement, reference-based motion correction, NU-refinement, global CTF refinement, and a final round of NU-refinement. The resulting reconstruction reached a nominal global resolution of 3.52 Å, as determined using the gold-standard Fourier shell correlation (FSC) 0.143 criterion.

#### Model Building and Refinement

DeepEMhancer^31^-sharpened maps were used for atomic model building. Initial models were generated using ModelAngelo^32^ and manually adjusted in Coot^33^. All models were refined by real-space refinement in Phenix^34^ and validated using MolProbity^35^. Structural biology software and computing environments used in this study were compiled and configured by the SBGrid Consortium^36^.

#### Hydroxyproline Structural Modeling

The hydroxyproline-containing peptide was manually modeled in Coot based on the structure of recombinant Ara h 2. Proline residues were substituted with hydroxyproline, and the resulting model was subjected to one round of geometry minimization in Phenix, using the recombinant Ara h 2 structure as a reference model for restraints.

#### Isothermal Titration Calorimetry

The non-hydroxylated Ara h 2 loop peptide (Pro loop; RDEDSYGRDPYSPSQDPYSPSQDPDRRDPYSPSPYD) and the corresponding hydroxyproline-containing peptide (Hyp loop; RDEDSYGRDPYS-Hyp-SQDPYS-Hyp-SQDPDRRDPYS-Hyp-SPYD) were synthesized by Peptide 2.0.

Isothermal titration calorimetry (ITC) measurements were performed at 25°C using either a MicroCal PEAQ-ITC or Nano ITC instrument (TA Instruments). All experiments were conducted in 10 mM Tris, pH 7.5, 150 mM NaCl. Antigens, comprising the loop peptides or Ara h 2 protein, were loaded into the sample cell, and the corresponding antibody Fab was loaded into the injection syringe. Titrations of P4EC10 with native Ara h 2 (nAra h 2) or the Hyp loop peptide were performed using the Nano ITC. Titrations of P4EC10 with the Pro loop peptide or recombinant Ara h 2 (rAra h 2), as well as titrations of P1BC6 and P1EC5 with rAra h 2, were performed using the MicroCal PEAQ-ITC.

For MicroCal PEAQ-ITC experiments, 300 µL of antigen was loaded into the sample cell and 100 µL of Fab into the syringe. Titrations consisted of 15 sequential injections: an initial 0.4-µL injection followed by 14 injections of 2.0 µL each, for a total injected volume of 28.4 µL. Injections were separated by 150 s to allow the signal to return to baseline.

For Nano ITC experiments, 350 µL of antigen was loaded into the sample cell and 70 µL of Fab into the syringe. Titrations consisted of 16 sequential injections: an initial 0.4-µL injection followed by 15 injections of 2.0 µL each. Injections were separated by 150 s.

All ITC binding data were analyzed by fitting the integrated binding isotherms to a simple independent binding-site model using software provided with the instrument, with binding affinity, stoichiometry, and enthalpy allowed to vary during fitting.

## Supporting information

PDB validation reports

## Acknowledgments

We thank the clinical personnel of the Jaffe Food Allergy Institute and General Pediatrics at ISMMS for patient recruitment.

## Author contributions

Conceptualization: MACL, GB; Methodology: AC, WFB, BB; Investigation: AC, WFB, BB, IAS; Visualization: AC, WFB, GB; Funding acquisition: MACL, GB; Supervision: MACL, GB; Writing – original draft: AC, WFB, BB, MACL, GB; Writing – review & editing: AC, WFB, BB, IAS, MACL, GB, HAS.

## Data and materials availability

The EM maps have been deposited in the Electron Microscopy Data Bank (EMDB) under accession code EMD-76196 and EMD-77097, for the P4EC10 and P1EC5-P1BC6 complexes, respectively and the accompanying atomic coordinates in the Protein Data Bank (PDB) under accession code 11YS and 13JR. The raw movies were deposited under EMPIAR-13385 and EMPIAR-13578

