## Supplementary Materials Fig S1 to S10, Tables S1 and S2 for "Structural basis of human antibody recognition of linear and conformational epitopes on the peanut allergen Ara h 2"

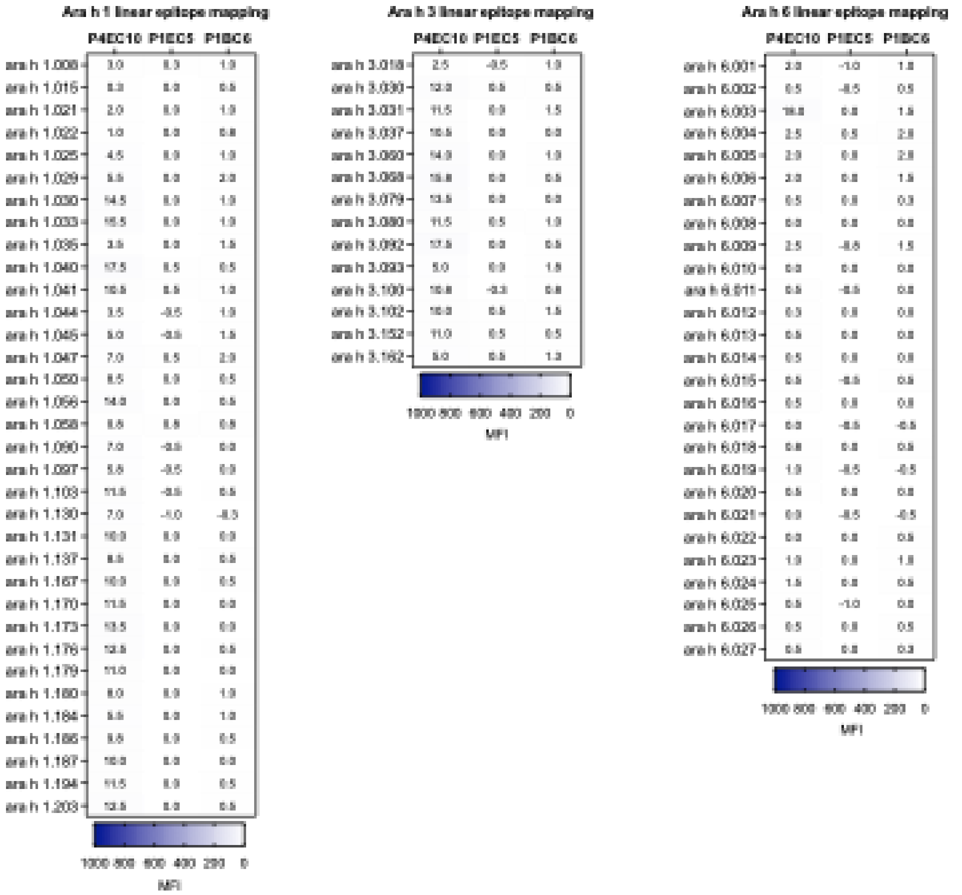


**Fig. S1. Linear epitope mapping of Ara h 2-specific IgE mAbs.** Binding of P4EC10, P1EC5, and P1BC6 to overlapping linear peptides spanning **(A)** Ara h 1, **(B)** Ara h 3, and **(C)** Ara h 6 was assessed by peptide-bead epitope mapping. Heatmaps show antibody binding to individual peptides, expressed as median fluorescence intensity (MFI). No substantial binding to the tested linear peptides was detected for any of the three antibodies.

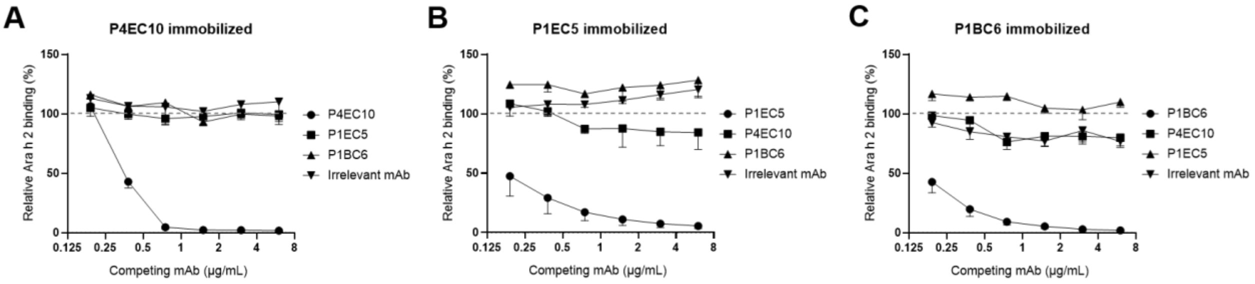


**Fig. S2. Competitive binding of Ara h 2-specific monoclonal antibodies**. Competitive ELISA was performed with P4EC10 **(A)**, P1EC5 **(B)**, or P1BC6 **(C)** immobilized on the plate. Ara h 2 binding was assessed in the presence of increasing concentrations of the indicated competing IgE mAbs. An irrelevant human mAb was included as a non-competing control. Binding was normalized to the corresponding condition without competing antibody, set to 100%. Data are presented as mean ± SD from three independent experiments.

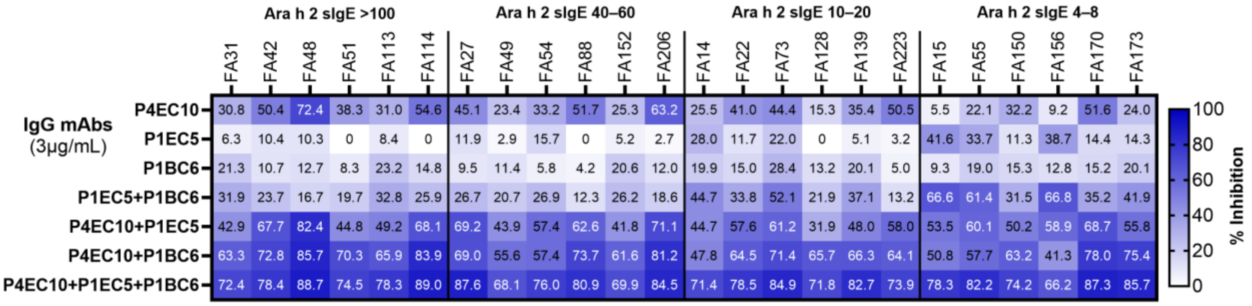


**Fig. S3. Individual plasma IgE inhibition profiles across Ara h 2 sIgE groups.** Heatmap showing inhibition of plasma IgE binding to Ara h 2 by recombinant human IgG1 versions of P4EC10, P1EC5, and P1BC6, alone or in the indicated combinations, for 24 peanut-allergic individuals. Individuals were stratified according to Ara h 2-specific IgE levels (>100, 40–60, 10–20, and 4–8 kUA/L; n = 6 per group). Plasma was tested at dilutions of 1:250, 1:25, 1:10, and 1:5, respectively. IgG1 mAbs were used at a total concentration of 3 µg/mL; for combinations, the total concentration was divided equally among the component antibodies. Values indicate percent inhibition relative to the corresponding no-IgG control.

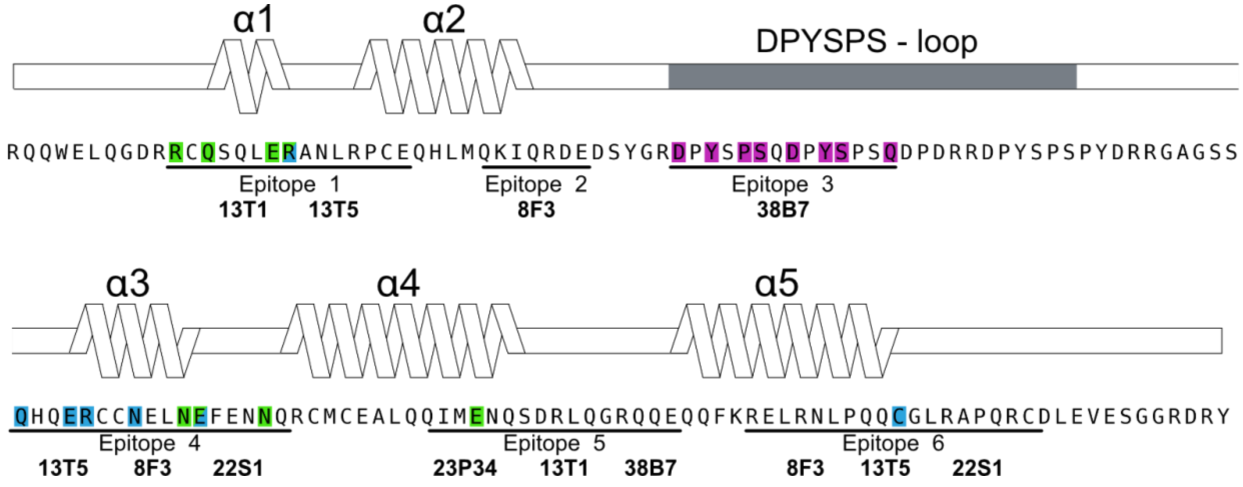


**Fig. S4. Sequence and secondary-structure representation of Ara h 2 epitopes recognized by recombinant human monoclonal antibodies.** Schematic representation of the Ara h 2 amino acid sequence and secondary structure, including α-helices α1–α5 and the DPYSPS-containing loop. The six antibody epitopes identified in previous studies are underlined and labeled Epitope 1–6. Recombinant human monoclonal antibodies assigned to each epitope are indicated below the corresponding sequence. Ara h 2 residues contacted by human monoclonal antibodies from this study, P4EC10, P1EC5, and P1BC6, are highlighted in magenta, blue, and green, respectively.

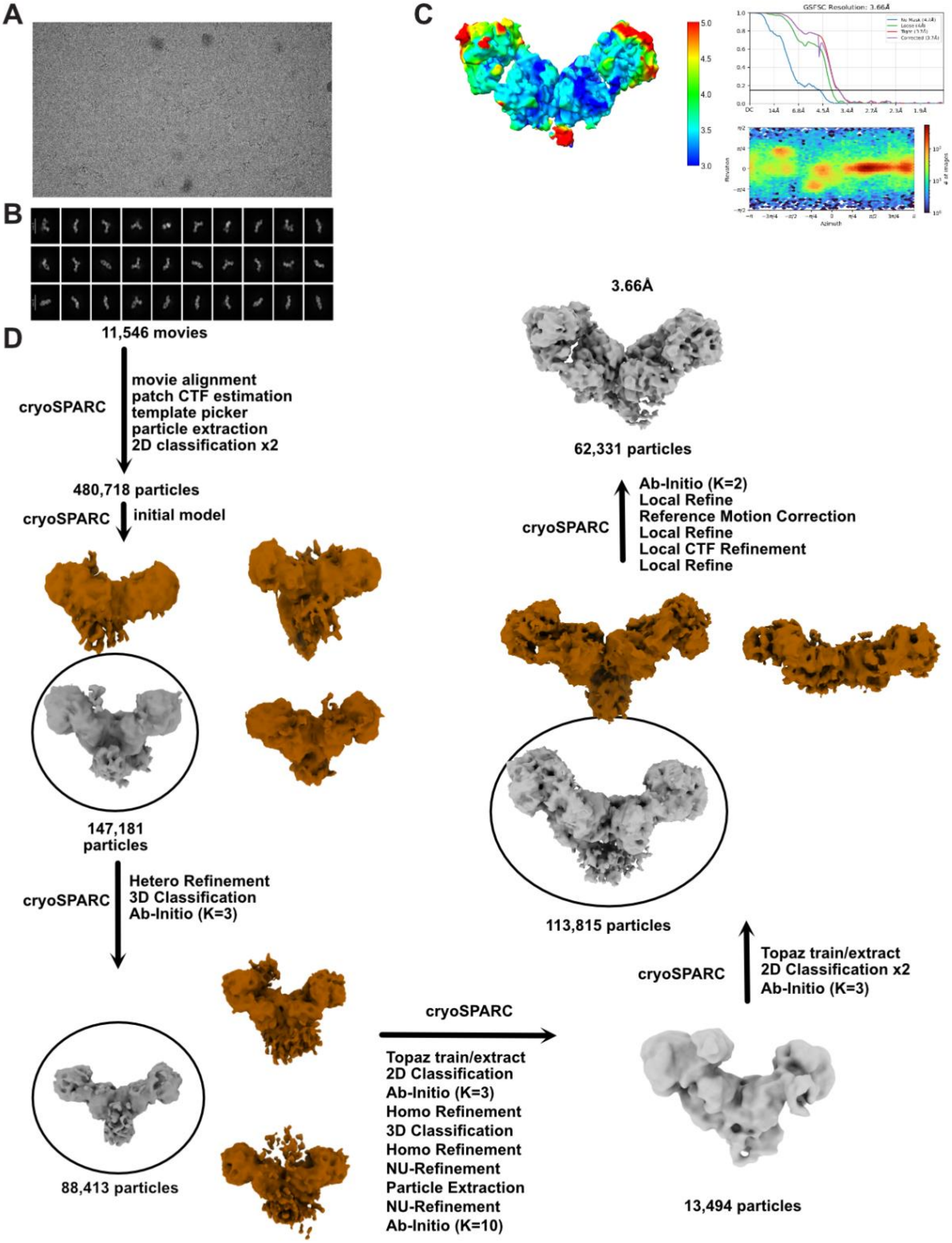


**Fig. S5. Cryo-EM data processing pipeline for determining the structure of P4EC10 Fab in complex with Ara h 2. (A)** Representative electron micrograph. **(B)** 2D class average obtained for Ara h 2 in complex with two P4EC10 Fabs. **(C)** Local resolution, gold-standard Fourier shell correlation curves and angular distribution plot, generated with cryoSPARC v5.0.3 **(D)** An outline of cryo-EM data processing workflow.

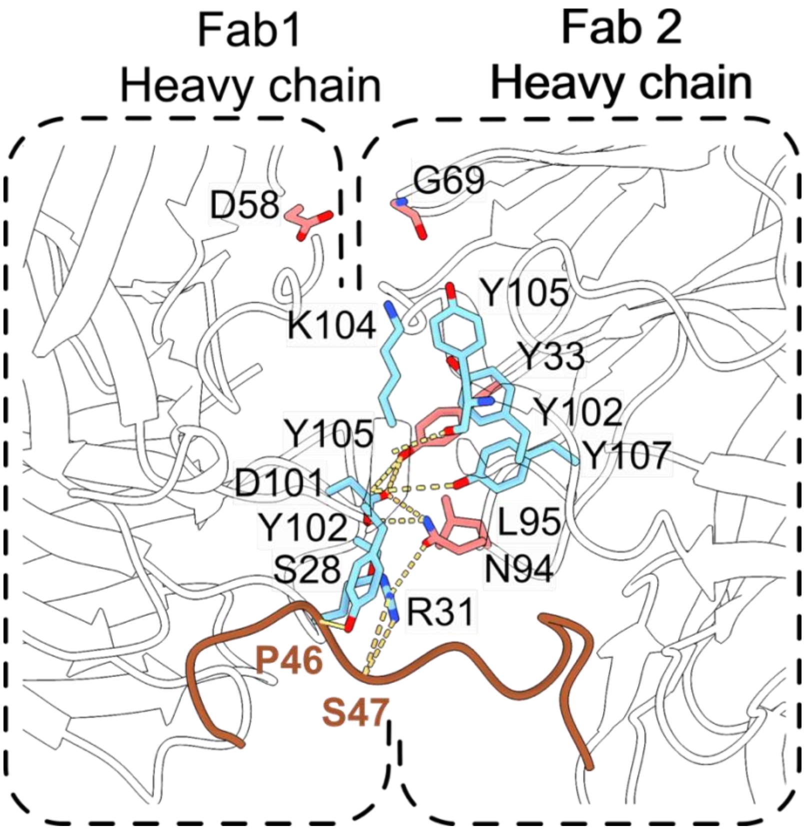


**Fig. S6. Zoom in on the P4EC10-Ara h 2 interface highlighting the homotypic Fab-Fab interactions.** N94 and Y33 from one Fab form hydrogen-bonding interactions with D101 and Y102 in HCDR3 of the neighboring Fab

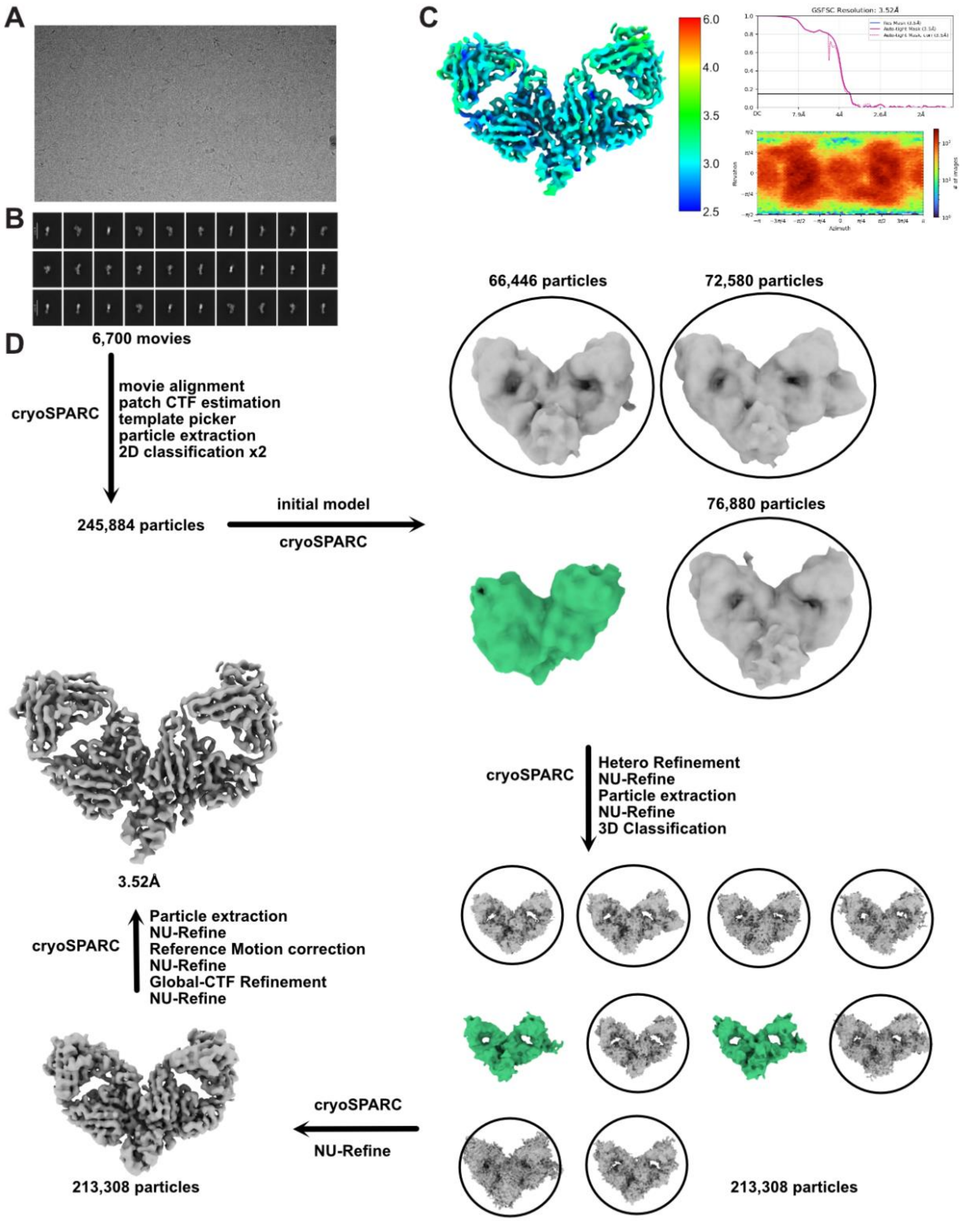


**Fig. S7. Cryo-EM data processing pipeline for determining the structure of P1EC5 and P1BC6 Fabs in complex with Ara h 2. (A)** Representative electron micrograph. **(B)** 2D class average obtained for Ara h 2 in complex with P1BC6 and P1EC5 Fabs. **(C)** Local resolution, gold-standard Fourier shell correlation curves and angular distribution plot, generated with cryoSPARC v5.0.3 **(D)** An outline of cryo-EM data processing workflow.

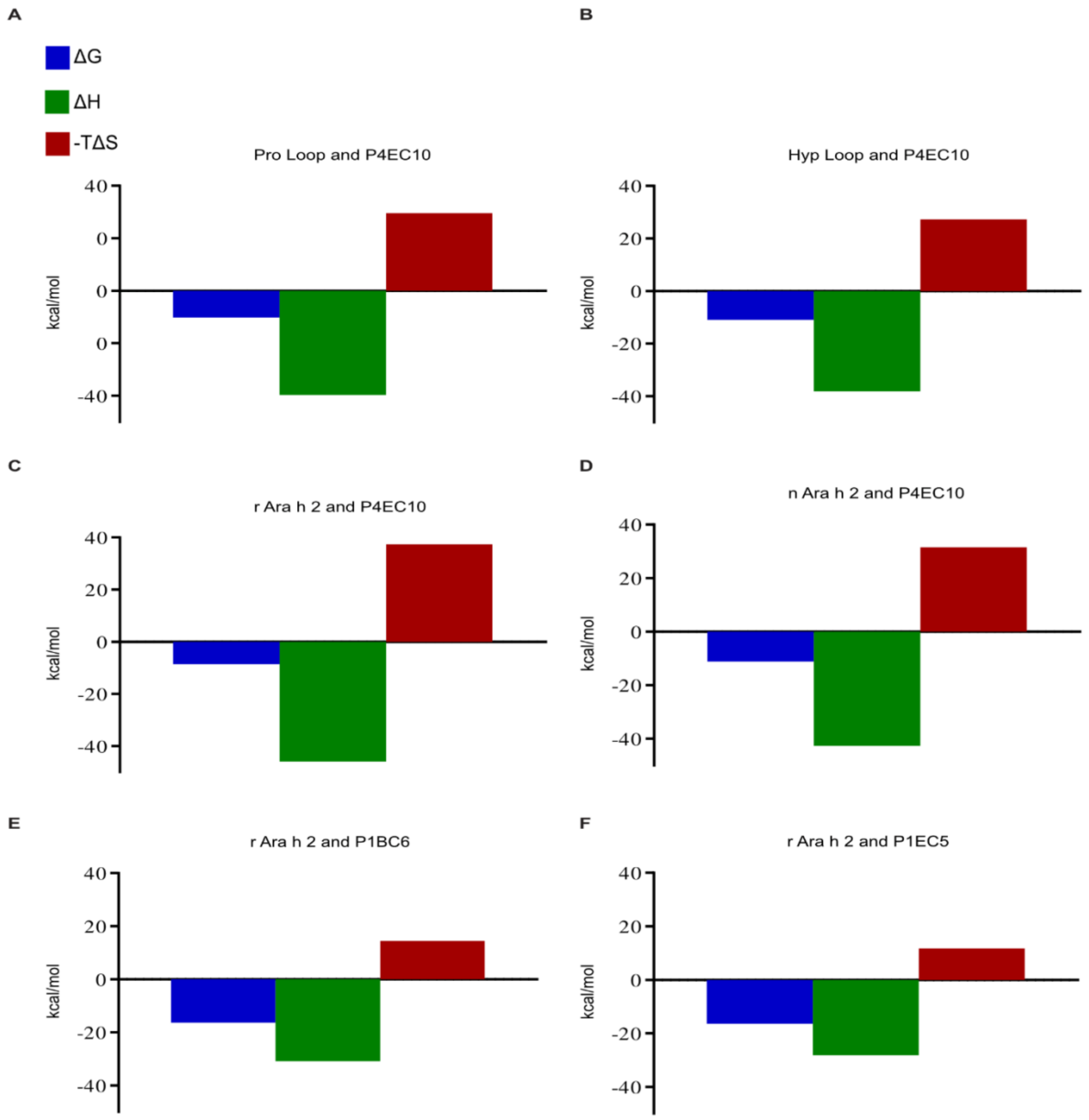


**Fig. S8. Thermodynamic signatures of Ara h 2-specific antibody binding.** Thermodynamic parameters derived from ITC measurements are shown for binding of P4EC10 to the non-hydroxylated (Pro) Ara h 2 repeat-loop peptide (A), the hydroxyproline-containing (Hyp) repeat-loop peptide (B), recombinant Ara h 2 (rAra h 2; C), and native Ara h 2 (nAra h 2; D), and for binding of P1BC6 (E) and P1EC5 (F) to rAra h 2. Gibbs free energy (ΔG), enthalpy (ΔH), and the entropic contribution (−TΔS) are shown in blue, green, and red, respectively, in kcal/mol.

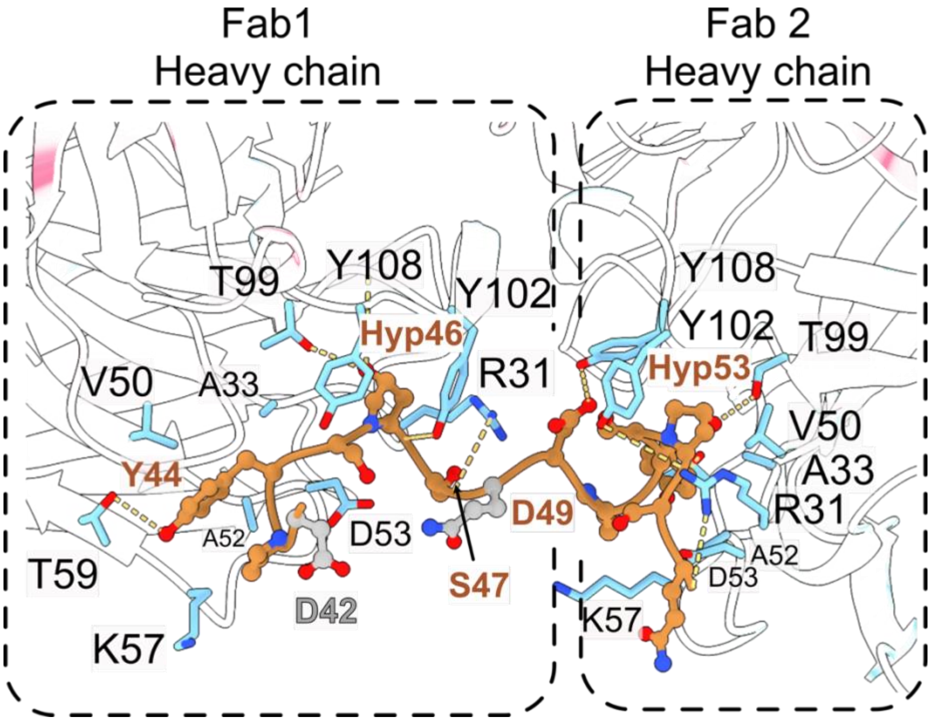


**Fig. S9. Modeling of hydroxyproline residues in the P4EC10–Ara h 2 interface.** Hydroxyproline residues were modeled into the P4EC10-bound Ara h 2 repeat-containing loop based on the cryo-EM structure of the recombinant complex. Substitution of P46 and P53 with hydroxyproline introduced additional potential hydrogen-bonding interactions at the antibody-allergen interface. The hydroxyl group at Ara h 2 residue 46 forms potential interactions with P4EC10 residues T99 and Y108, whereas the hydroxyl group at residue 53 forms a potential interaction with T99. Hydrogen bonds are indicated by yellow dashed lines.

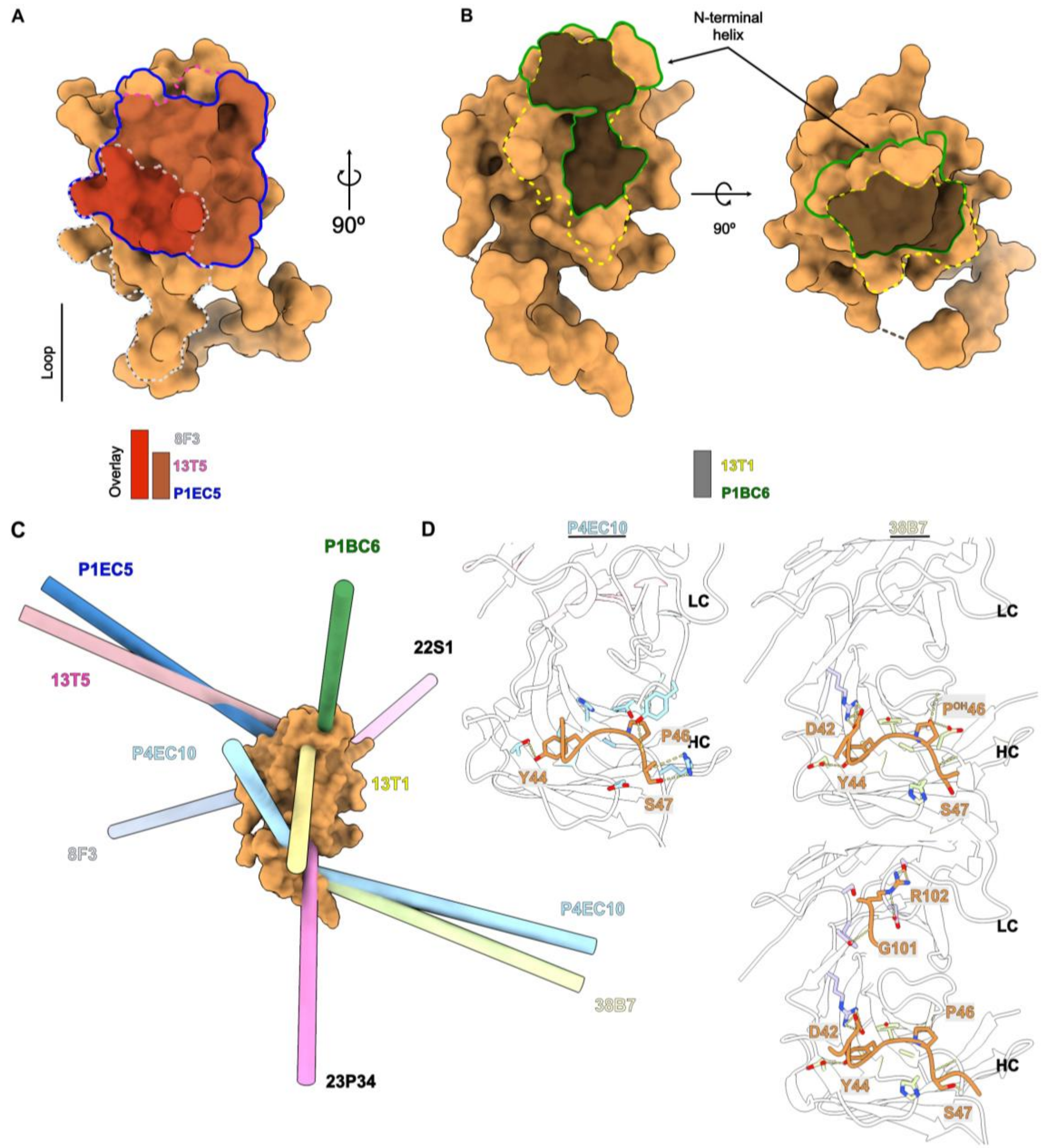


**Fig. S10. Comparison of P4EC10, P1EC5, and P1BC6 with previously characterized Ara h 2-specific antibodies.** Ara h 2 is shown in light-brown representation (PDB: 8SJ6). (**A**) Footprints of antibodies 8F3 (PDB: 8SJ6), 13T5 (PDB: 8G4P), and P1EC5 (PDB: 13JR) are outlined in grey, magenta, and blue, respectively. (**B**) Footprints of antibodies 13T1 (PDB: 8G4P) and P1BC6 (PDB: 13JR) are outlined in yellow and green, respectively. Overlapping regions between antibody footprints are highlighted. (**C**) Comparison of approach orientations of structurally characterized Ara h 2-specific antibodies. Each antibody is represented by a cylinder indicating its principal axis relative to the antigen. Structures shown are 22S1 (PDB: 8DB4), 23P34 (PDB: 9CMC), 38B7 (PDB: 8SJ6),8F3, 13T1, 13T5, P1EC5, P1BC6, P4EC10. (**D**) Comparison of P4EC10 bound to recombinant Ara h 2 with 38B7 bound to hydroxyproline containing peptide (upper panel) and full-length recombinant Ara h 2 (lower panel). Antibody contact residues are shown as sticks and colored cyan (P4EC10) and lime (38B7), while Ara h 2 is colored light brown.

**TABLE S1. Characteristics of plasma donors used in this study**

| Subject ID | Group | Sex | Age (years) | Peanut status | Total IgE (kU/L) | Peanut sIgE (kUA/L) | Ara h 1 sIgE (kUA/L) | Ara h 2 sIgE (kUA/L) | Ara h 3 sIgE (kUA/L) | Ara h 6 sIgE (kUA/L) |
|---|---|---|---|---|---|---|---|---|---|---|
| **Ara h 2 sIgE >100 kUA/L** | | | | | | | | | | |
| FA031 | PA | M | 9 | Allergic | 2229 | >100 | >100 | >100 | 96.6 | >100 |
| FA042 | PA | F | 8 | Allergic | 1598 | >100 | >100 | >100 | 78.2 | 96.8 |
| FA048 | PA | F | 12 | Allergic | 2275 | >100 | >100 | >100 | >100 | >100 |
| FA051 | PA | F | 9 | Allergic | 1432 | >100 | >100 | >100 | 11 | >100 |
| FA113 | PA | F | 8 | Allergic | 1675 | >100 | >100 | >100 | 20.8 | >100 |
| FA114 | PA | M | 17 | Allergic | 1650 | >100 | >100 | >100 | 18 | >100 |
| **Ara h 2 sIgE 40–60 kUA/L** | | | | | | | | | | |
| FA027 | PA | M | 20 | Allergic | 322 | 72.2 | 37.8 | 50.4 | 12.1 | 28.9 |
| FA049 | PA | M | 17 | Allergic | 317 | 65.8 | 48.9 | 52.9 | 1.78 | 33.3 |
| FA054 | PA | F | 4 | Allergic | 1229 | 64.1 | 25.5 | 41.9 | 0.8 | 45 |
| FA088 | PA | M | 19 | Allergic | 343 | 68.1 | 8.74 | 57.3 | 4.27 | 22.9 |
| FA152 | PA | M | 10 | Allergic | 679 | 64.3 | 20.6 | 56.3 | 0.29 | 31.1 |
| FA206 | PA | M | 12 | Allergic | 428 | 61.5 | 35.3 | 45.8 | 5.96 | 14.9 |
| **Ara h 2 sIgE 10–20 kUA/L** | | | | | | | | | | |
| FA014 | PA | M | 4 | Allergic | 252 | 18.2 | 0.17 | 18.6 | <0.1 | 7.25 |
| FA022 | PA | F | 14 | Allergic | 109 | 26.4 | 10.1 | 19.5 | 0.57 | 10.5 |
| FA073 | PA | F | 27 | Allergic | 102 | 25 | 12.1 | 14.5 | 2.33 | 8.69 |
| FA128 | PA | F | 17 | Allergic | 403 | 28 | 1.55 | 14.8 | 0.4 | 26.5 |
| FA139 | PA | F | 11 | Allergic | 169 | 31.6 | 17.8 | 11.1 | 1.71 | 11.8 |
| FA223 | PA | M | 15 | Allergic | 592 | 34.7 | 19.2 | 19.5 | 1.79 | 13.3 |
| **Ara h 2 sIgE 4–8 kUA/L** | | | | | | | | | | |
| FA015 | PA | M | 4 | Allergic | 197 | 6.02 | <0.1 | 4.76 | <0.1 | 3.51 |
| FA055 | PA | M | 13 | Allergic | 977 | 9.44 | <0.1 | 6.96 | <0.1 | 6.26 |
| FA150 | PA | M | 18 | Allergic | 072 | 13.1 | 4.29 | 4.06 | 0.38 | 5.97 |
| FA156 | PA | M | 4 | Allergic | 778 | 4.23 | 0.35 | 4.71 | <0.1 | 0.23 |
| FA170 | PA | F | 22 | Allergic | 154 | 15.4 | 4.12 | 5.34 | 0.76 | 5.77 |
| FA173 | PA | F | 17 | Allergic | 412 | 15.2 | <0.1 | 5.51 | 0.99 | 5.65 |
| **Non-PA controls** | | | | | | | | | | |
| HC005 | non-PA | F | 11 | Tolerant | n.a. | n.a. | n.a. | n.a. | n.a. | n.a. |
| HC006 | non-PA | F | 16 | Tolerant | n.a. | n.a. | n.a. | n.a. | n.a. | n.a. |

PA, peanut-allergic; non-PA, non-peanut-allergic; sIgE, allergen-specific IgE; n.a., not available. PA subjects were stratified according to Ara h 2 sIgE levels for plasma IgE inhibition assays. Plasma from FA031, FA042, and FA048 was pooled for plasma-sensitized RBL-SX38 degranulation, IgG-mediated degranulation-inhibition, and PCA assays. Plasma from HC005 and HC006 was pooled for non-PA control assays.

**TABLE S2. Cryo-EM data collection, refinement and validation statistics**

| | P4EC10 – Ara h 2<br>(EMD- 76196)<br>(PDB 11YS) | P1EC5 – P1BC6 – Ara h 2<br>(EMD-77097)<br>(PDB 13JR) |
|---|---|---|
| **Data collection and processing** | | |
| Magnification | 81,000 | 81,000 |
| Voltage (kV) | 300 | 300 |
| Electron exposure (e–/Å$^2$) | 57.93 | 55.24 |
| Defocus range (μm) | -1.5, -2.5 | -1.5, -2.5 |
| Pixel size (Å) | 0.844 | 0.826 |
| Symmetry imposed | C1 | C1 |
| Initial particle images (no.) | 480,718 | 245,884 |
| Final particle images (no.) | 62,331 | 211,894 |
| Map resolution (Å) | 3.7 | 3.52 |
| FSC threshold | 0.143 | 0.143 |
| Map resolution range (Å) | 3.0-5.0 | 2.5-4.0 |
| **Refinement** | | |
| Initial model used (PDB code) | | |
| Model resolution (Å) | 3.7 | 3.6 |
| FSC threshold | 0.143 | 0.143 |
| Map sharpening *B* factor (Å$^2$) | 130.7 | 103.6 |
| Model composition | | |
| Non-hydrogen atoms | 6655 | 7323 |
| Protein residues | 870 | 960 |
| *B* factors (Å$^2$) | | |
| Protein | 77.47 | 60.74 |
| R.m.s. deviations | | |
| Bond lengths (Å) | 0.005 | 0.005 |

| | | |
|---|---|---|
| Bond angles (°) | 0.830 | 0.707 |
| Validation | | |
| MolProbity score | 0.81 | 1.04 |
| Clashscore | 0.53 | 0.49 |
| Poor rotamers (%) | 0.00 | 0.00 |
| Ramachandran plot | | |
| Favored (%) | 97.31 | 94.51 |
| Allowed (%) | 2.69 | 5.49 |
| Disallowed (%) | 0.00 | 0.00 |
