## Supplementary material for "Structural basis of human antibody recognition of linear and conformational epitopes on the peanut allergen Ara h 2": PDB validation reports

### Full wwPDB EM Validation Report (i)

May 1, 2026 – 02:26 PM EDT

| | | |
|---|---|---|
| PDB ID | : | 11YS / pdb_000011ys |
| EMDB ID | : | EMD-76196 |
| Title | : | Cryo-EM structure of Ara h 2 peanut allergen in complex with 2 P4EC10 Fab fragments |
| Deposited on | : | 2026-03-18 |
| Resolution | : | 3.70 Å(reported) |

**This wwPDB validation report is for manuscript review**

This is a Full wwPDB EM Validation Report.

This report is produced by the wwPDB biocuration pipeline after annotation of the structure.

We welcome your comments at **
A user guide is available at
https://www.wwpdb.org/validation/2017/EMValidationReportHelp
with specific help available everywhere you see the (i) symbol.

The types of validation reports are described at
http://www.wwpdb.org/validation/2017/FAQs#types.

---

The following versions of software and data (see references (i)) were used in the production of this report:

| | | |
|---|---|---|
| EMDB validation analysis | : | 0.0.1.dev132 |
| MolProbity | : | 4-5-2 with Phenix2.0 |
| Percentile statistics | : | 20250101.v01 (using entries in the PDB archive January 1st 2025) |
| EM percentile statistics | : | 202505.v01 (Using data in the EMDB archive up until May 2025) |
| MapQ | : | 1.9.13 |
| Ideal geometry (proteins) | : | Engh & Huber (2001) |
| Ideal geometry (DNA, RNA) | : | Parkinson et al. (1996) |
| Validation Pipeline (wwPDB-VP) | : | 2.49 |

For Manuscript Review

### 1 Overall quality at a glance ⓘ

The following experimental techniques were used to determine the structure:
*ELECTRON MICROSCOPY*

The reported resolution of this entry is 3.70 Å.

Percentile scores (ranging between 0-100) for global validation metrics of the entry are shown in the following graphic. The table shows the number of entries on which the scores are based.

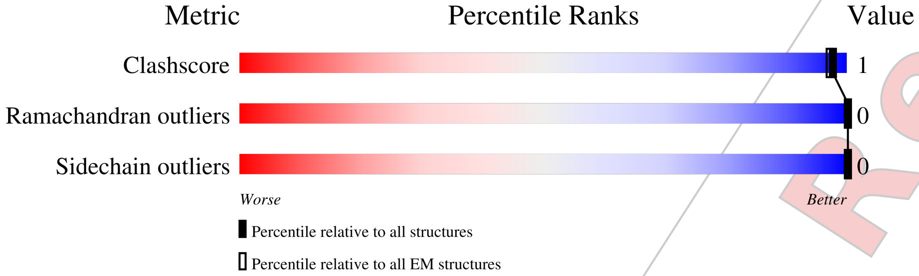


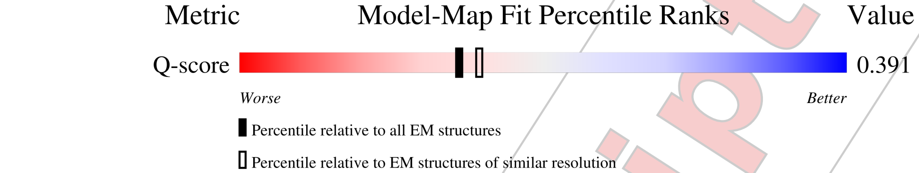


| Metric | Whole archive (#Entries) | EM structures (#Entries) | Similar EM resolution (#Entries, resolution range(Å)) |
|---|---|---|---|
| Clashscore | 229148 | 23984 | - |
| Ramachandran outliers | 224038 | 23583 | - |
| Sidechain outliers | 223484 | 23102 | - |
| Q-score | - | 25397 | 11569 ( 3.20 - 4.20 ) |

The table below summarises the geometric issues observed across the polymeric chains and their fit to the map. The red, orange, yellow and green segments of the bar indicate the fraction of residues that contain outliers for >=3, 2, 1 and 0 types of geometric quality criteria respectively. A grey segment represents the fraction of residues that are not modelled. The numeric value for each fraction is indicated below the corresponding segment, with a dot representing fractions <=5% The upper red bar (where present) indicates the fraction of residues that have poor fit to the EM map (all-atom inclusion < 40%). The numeric value is given above the bar.

| Mol | Chain | Length | Quality of chain |
|---|---|---|---|
| 1 | H | 229 | 5% / 93% • 6% |
| 1 | M | 229 | 6% / 94% • 5% |
| 2 | K | 214 | 7% / 97% •• |

*Continued on next page...*

For Manuscript Review

*Continued from previous page...*

| Mol | Chain | Length | Quality of chain |
|---|---|---|---|
| 2 | L | 214 | 5% 97% • • |
| 3 | A | 151 | • 9% 91% |

For Manuscript Review

### 2 Entry composition (i)

There are 3 unique types of molecules in this entry. The entry contains 6655 atoms, of which 0 are hydrogens and 0 are deuteriums.

In the tables below, the AltConf column contains the number of residues with at least one atom in alternate conformation and the Trace column contains the number of residues modelled with at most 2 atoms.

- Molecule 1 is a protein called P4EC10 Fab heavy chain.

| Mol | Chain | Residues | Atoms | | | | | AltConf | Trace |
|---|---|---|---|---|---|---|---|---|---|
| 1 | H | 215 | Total 1633 | C 1035 | N 278 | O 314 | S 6 | 0 | 0 |
| 1 | M | 218 | Total 1655 | C 1049 | N 282 | O 317 | S 7 | 0 | 0 |

- Molecule 2 is a protein called P4EC10 Fab light chain.

| Mol | Chain | Residues | Atoms | | | | | AltConf | Trace |
|---|---|---|---|---|---|---|---|---|---|
| 2 | L | 211 | Total 1624 | C 1018 | N 275 | O 327 | S 4 | 0 | 0 |
| 2 | K | 212 | Total 1633 | C 1023 | N 276 | O 330 | S 4 | 0 | 0 |

- Molecule 3 is a protein called Conglutin-7.

| Mol | Chain | Residues | Atoms | | | | AltConf | Trace |
|---|---|---|---|---|---|---|---|---|
| 3 | A | 14 | Total 110 | C 68 | N 16 | O 26 | 0 | 0 |

For Manuscript Review

### 3 Residue-property plots (i)

These plots are drawn for all protein, RNA, DNA and oligosaccharide chains in the entry. The first graphic for a chain summarises the proportions of the various outlier classes displayed in the second graphic. The second graphic shows the sequence view annotated by issues in geometry and atom inclusion in map density. Residues are color-coded according to the number of geometric quality criteria for which they contain at least one outlier: green = 0, yellow = 1, orange = 2 and red = 3 or more. A red diamond above a residue indicates a poor fit to the EM map for this residue (all-atom inclusion < 40%). Stretches of 2 or more consecutive residues without any outlier are shown as a green connector. Residues present in the sample, but not in the model, are shown in grey.

- Molecule 1: P4EC10 Fab heavy chain

Chain H:

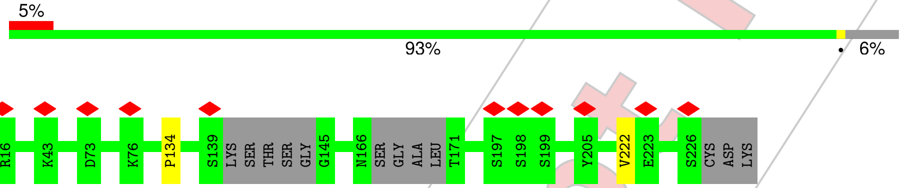


- Molecule 1: P4EC10 Fab heavy chain

Chain M:

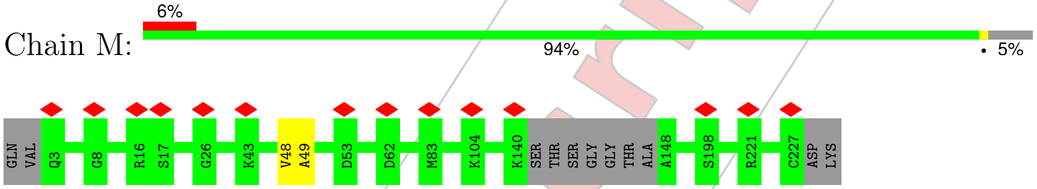


- Molecule 2: P4EC10 Fab light chain

Chain L:

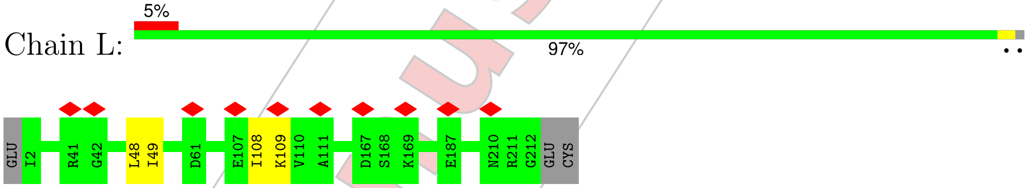


- Molecule 2: P4EC10 Fab light chain

Chain K:

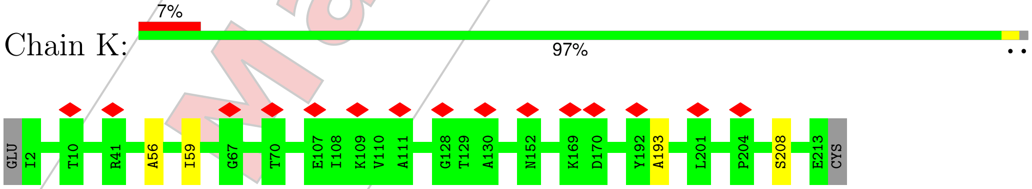


- Molecule 3: Conglutin-7

Chain A:

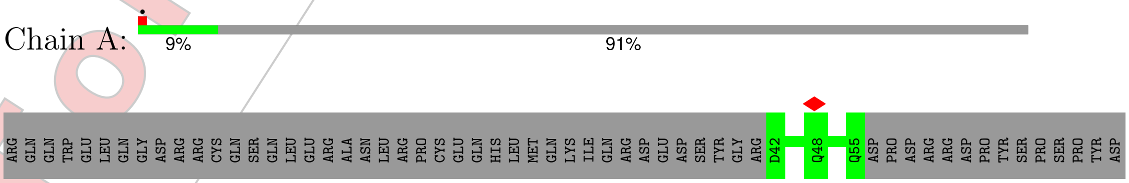


For Manuscript Review

ARG ARG GLY ALA GLY SER SER GLN HIS GLN GLU ARG CYS CYS ASN GLU LEU ASN GLU PHE GLU ASN ASN GLN ARG CYS MET CYS GLU ALA LEU GLN GLN ILE MET GLU ASN GLN SER ASP ARG LEU GLN GLY ARG GLN GLN GLU GLN GLN PHE LYS ARG GLU LEU ARG ASN LEU PRO GLN

GLN CYS GLY LEU ARG ALA PRO GLN ARG CYS ASP LEU GLU VAL GLU SER GLY GLY ARG ASP ARG TYR

For Manuscript Review

### 4 Experimental information (i)

| Property | Value | Source |
|---|---|---|
| EM reconstruction method | SINGLE PARTICLE | Depositor |
| Imposed symmetry | POINT, C1 | Depositor |
| Number of particles used | 62331 | Depositor |
| Resolution determination method | FSC 0.143 CUT-OFF | Depositor |
| CTF correction method | PHASE FLIPPING AND AMPLITUDE CORRECTION | Depositor |
| Microscope | TFS KRIOS | Depositor |
| Voltage (kV) | 300 | Depositor |
| Electron dose ($e^-/\text{Å}^2$) | 57.93 | Depositor |
| Minimum defocus (nm) | 1500 | Depositor |
| Maximum defocus (nm) | 2500 | Depositor |
| Magnification | Not provided | |
| Image detector | GATAN K3 BIOQUANTUM (6k x 4k) | Depositor |
| Maximum map value | 2.004 | Depositor |
| Minimum map value | -0.002 | Depositor |
| Average map value | 0.000 | Depositor |
| Map value standard deviation | 0.015 | Depositor |
| Recommended contour level | 0.12 | Depositor |
| Map size (Å) | 337.6, 337.6, 337.6 | wwPDB |
| Map dimensions | 400, 400, 400 | wwPDB |
| Map angles (°) | 90.0, 90.0, 90.0 | wwPDB |
| Pixel spacing (Å) | 0.84400004, 0.84400004, 0.84400004 | Depositor |

For Manuscript Review

### 5 Model quality (i)

#### 5.1 Standard geometry (i)

The Z score for a bond length (or angle) is the number of standard deviations the observed value is removed from the expected value. A bond length (or angle) with $|Z| > 5$ is considered an outlier worth inspection. RMSZ is the root-mean-square of all Z scores of the bond lengths (or angles).

| Mol | Chain | Bond lengths | | Bond angles | |
|---|---|---|---|---|---|
| | | RMSZ | #\|Z\| >5 | RMSZ | #\|Z\| >5 |
| 1 | H | 0.23 | 0/1672 | 0.46 | 0/2273 |
| 1 | M | 0.21 | 0/1695 | 0.45 | 0/2304 |
| 2 | K | 0.23 | 0/1673 | 0.43 | 0/2278 |
| 2 | L | 0.25 | 0/1664 | 0.50 | 0/2266 |
| 3 | A | 0.34 | 0/115 | 0.60 | 0/159 |
| All | All | 0.23 | 0/6819 | 0.47 | 0/9280 |

There are no bond length outliers.

There are no bond angle outliers.

There are no chirality outliers.

There are no planarity outliers.

#### 5.2 Too-close contacts (i)

In the following table, the Non-H and H(model) columns list the number of non-hydrogen atoms and hydrogen atoms in the chain respectively. The H(added) column lists the number of hydrogen atoms added and optimized by MolProbity. The Clashes column lists the number of clashes within the asymmetric unit, whereas Symm-Clashes lists symmetry-related clashes.

| Mol | Chain | Non-H | H(model) | H(added) | Clashes | Symm-Clashes |
|---|---|---|---|---|---|---|
| 1 | H | 1633 | 0 | 1595 | 2 | 0 |
| 1 | M | 1655 | 0 | 1623 | 1 | 0 |
| 2 | K | 1633 | 0 | 1567 | 2 | 0 |
| 2 | L | 1624 | 0 | 1561 | 2 | 0 |
| 3 | A | 110 | 0 | 89 | 0 | 0 |
| All | All | 6655 | 0 | 6435 | 7 | 0 |

The all-atom clashscore is defined as the number of clashes found per 1000 atoms (including hydrogen atoms). The all-atom clashscore for this structure is 1.

All (7) close contacts within the same asymmetric unit are listed below, sorted by their clash magnitude.

For Manuscript Review

| Atom-1 | Atom-2 | Interatomic distance (Å) | Clash overlap (Å) |
|---|---|---|---|
| 1:M:48:VAL:HG12 | 1:M:49:ALA:H | 1.67 | 0.60 |
| 2:L:108:ILE:HG22 | 2:L:109:LYS:HG3 | 1.94 | 0.48 |
| 2:L:48:LEU:O | 2:L:49:ILE:HD13 | 2.15 | 0.47 |
| 2:K:56:ALA:HB3 | 2:K:59:ILE:HD12 | 1.96 | 0.46 |
| 2:K:193:ALA:HB2 | 2:K:208:SER:HB2 | 1.98 | 0.45 |
| 1:H:134:PRO:HB3 | 1:H:222:VAL:HG22 | 2.00 | 0.43 |
| 1:H:134:PRO:HB2 | 1:H:222:VAL:HG13 | 2.02 | 0.40 |

There are no symmetry-related clashes.

#### 5.3 Torsion angles (i)

##### 5.3.1 Protein backbone (i)

In the following table, the Percentiles column shows the percent Ramachandran outliers of the chain as a percentile score with respect to all PDB entries followed by that with respect to all EM entries.

The Analysed column shows the number of residues for which the backbone conformation was analysed, and the total number of residues.

| Mol | Chain | Analysed | Favoured | Allowed | Outliers | Percentiles | |
|---|---|---|---|---|---|---|---|
| 1 | H | 209/229 (91%) | 205 (98%) | 4 (2%) | 0 | 100 | 100 |
| 1 | M | 214/229 (93%) | 208 (97%) | 6 (3%) | 0 | 100 | 100 |
| 2 | K | 210/214 (98%) | 206 (98%) | 4 (2%) | 0 | 100 | 100 |
| 2 | L | 209/214 (98%) | 200 (96%) | 9 (4%) | 0 | 100 | 100 |
| 3 | A | 12/151 (8%) | 12 (100%) | 0 | 0 | 100 | 100 |
| All | All | 854/1037 (82%) | 831 (97%) | 23 (3%) | 0 | 100 | 100 |

There are no Ramachandran outliers to report.

##### 5.3.2 Protein sidechains (i)

In the following table, the Percentiles column shows the percent sidechain outliers of the chain as a percentile score with respect to all PDB entries followed by that with respect to all EM entries.

The Analysed column shows the number of residues for which the sidechain conformation was analysed, and the total number of residues.

For Manuscript Review

| Mol | Chain | Analysed | Rotameric | Outliers | Percentiles | |
|---|---|---|---|---|---|---|
| 1 | H | 182/193 (94%) | 182 (100%) | 0 | 100 | 100 |
| 1 | M | 185/193 (96%) | 185 (100%) | 0 | 100 | 100 |
| 2 | K | 186/188 (99%) | 186 (100%) | 0 | 100 | 100 |
| 2 | L | 185/188 (98%) | 185 (100%) | 0 | 100 | 100 |
| 3 | A | 14/139 (10%) | 14 (100%) | 0 | 100 | 100 |
| All | All | 752/901 (84%) | 752 (100%) | 0 | 100 | 100 |

There are no protein residues with a non-rotameric sidechain to report.

Sometimes sidechains can be flipped to improve hydrogen bonding and reduce clashes. All (4) such sidechains are listed below:

| Mol | Chain | Res | Type |
|---|---|---|---|
| 2 | L | 189 | HIS |
| 1 | M | 208 | ASN |
| 2 | K | 38 | GLN |
| 2 | K | 39 | HIS |

##### 5.3.3 RNA (i)

There are no RNA molecules in this entry.

#### 5.4 Non-standard residues in protein, DNA, RNA chains (i)

There are no non-standard protein/DNA/RNA residues in this entry.

#### 5.5 Carbohydrates (i)

There are no oligosaccharides in this entry.

#### 5.6 Ligand geometry (i)

There are no ligands in this entry.

#### 5.7 Other polymers (i)

There are no such residues in this entry.

For Manuscript Review

#### 5.8 Polymer linkage issues (i)

There are no chain breaks in this entry.

For Manuscript Review

### 6 Map visualisation (i)

This section contains visualisations of the EMDB entry EMD-76196. These allow visual inspection of the internal detail of the map and identification of artifacts.

Images derived from a raw map, generated by summing the deposited half-maps, are presented below the corresponding image components of the primary map to allow further visual inspection and comparison with those of the primary map.

#### 6.1 Orthogonal projections (i)

##### 6.1.1 Primary map

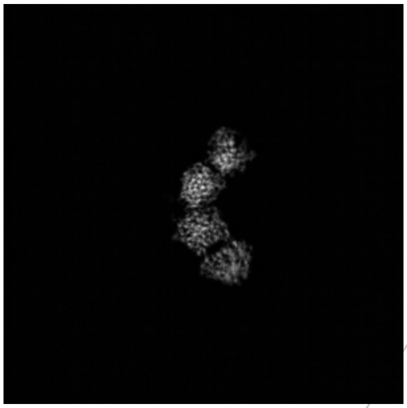

X

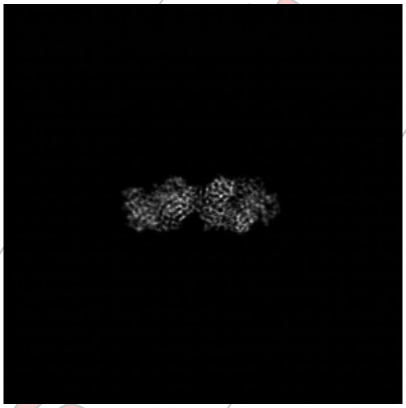

Y

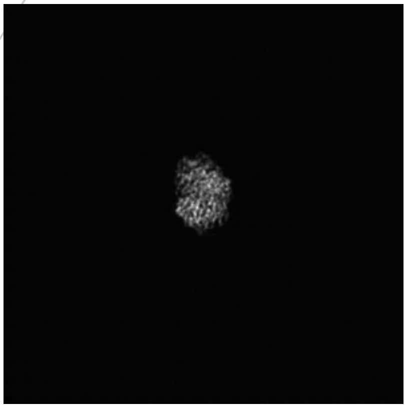

Z

##### 6.1.2 Raw map

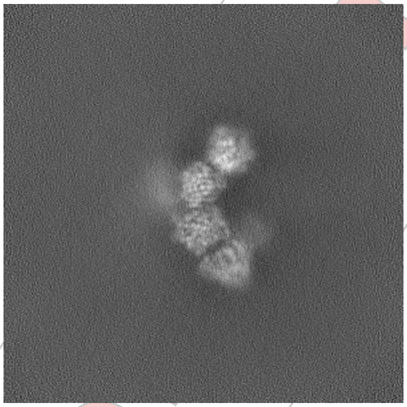

X

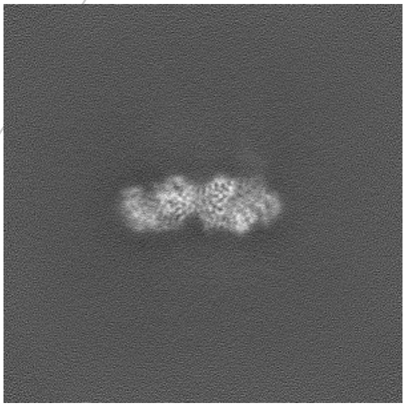

Y

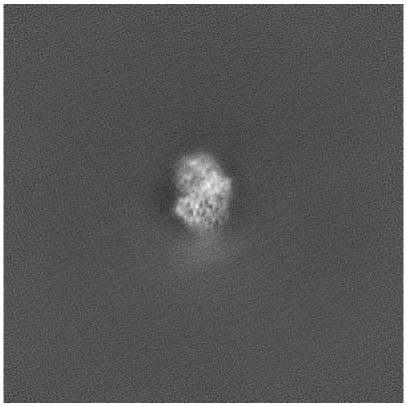

Z

The images above show the map projected in three orthogonal directions.

For Manuscript Review

#### 6.2 Central slices (i)

##### 6.2.1 Primary map

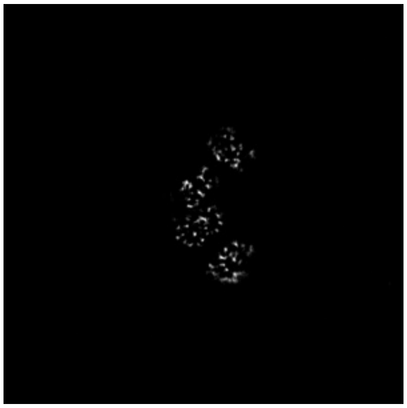

X Index: 200

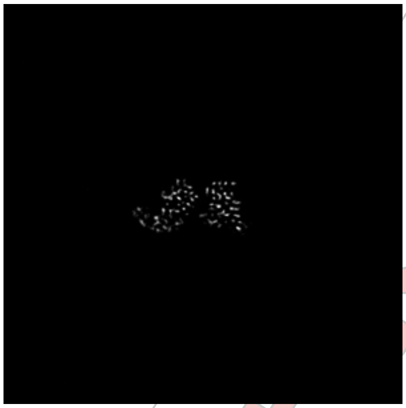

Y Index: 200

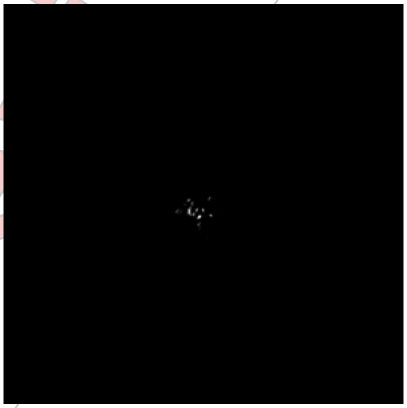

Z Index: 200

##### 6.2.2 Raw map

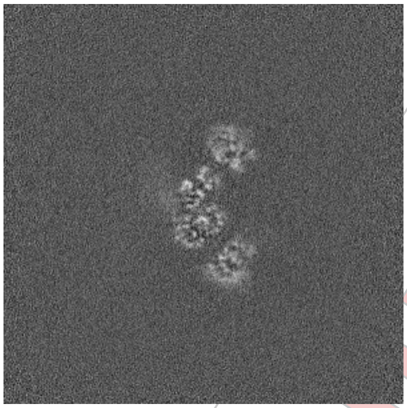

X Index: 200

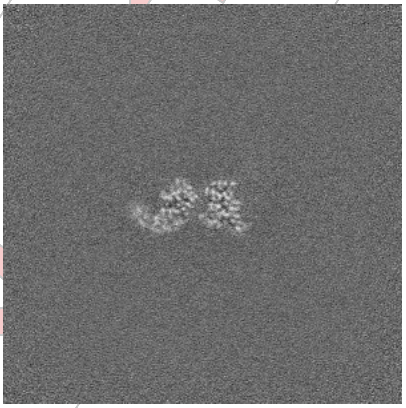

Y Index: 200

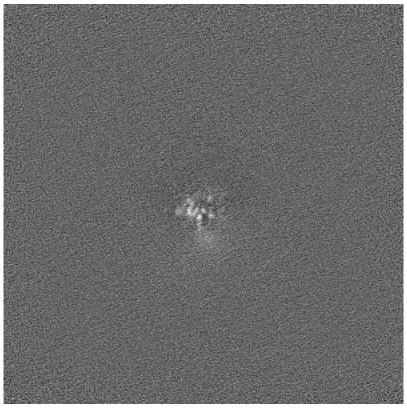

Z Index: 200

The images above show central slices of the map in three orthogonal directions.

For Manuscript Review

#### 6.3 Largest variance slices (i)

##### 6.3.1 Primary map

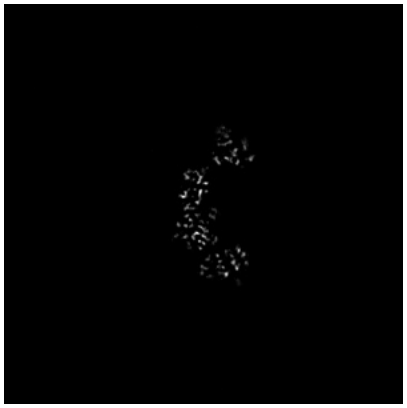
X Index: 193

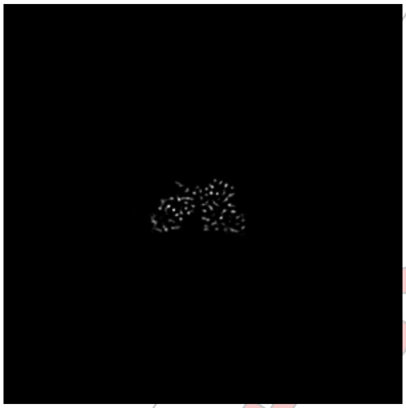
Y Index: 194

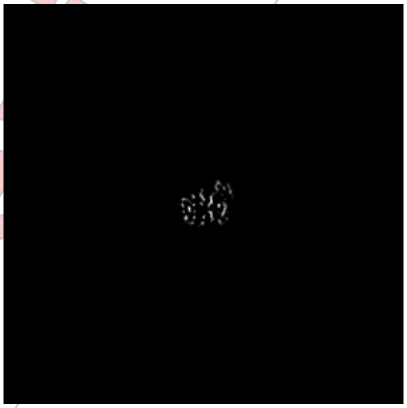
Z Index: 217

##### 6.3.2 Raw map

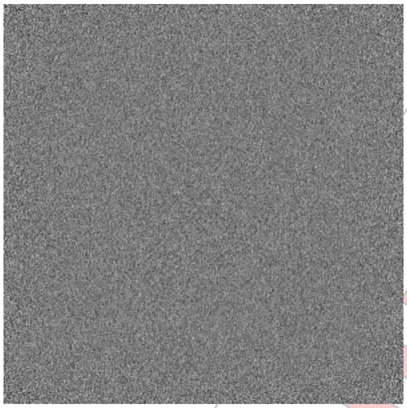
X Index: 0

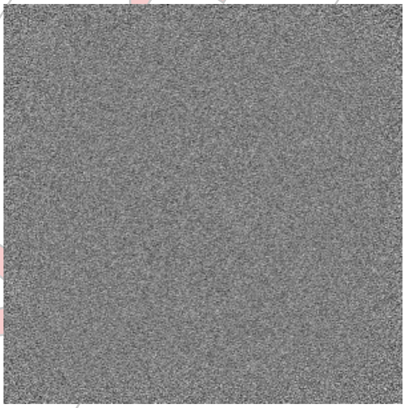
Y Index: 0

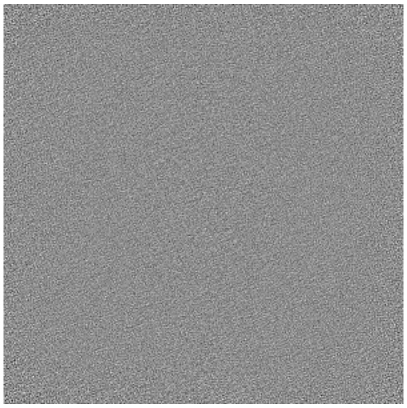
Z Index: 0

The images above show the largest variance slices of the map in three orthogonal directions.

For Manuscript Review

#### 6.4 Orthogonal standard-deviation projections (False-color) ⓘ

##### 6.4.1 Primary map

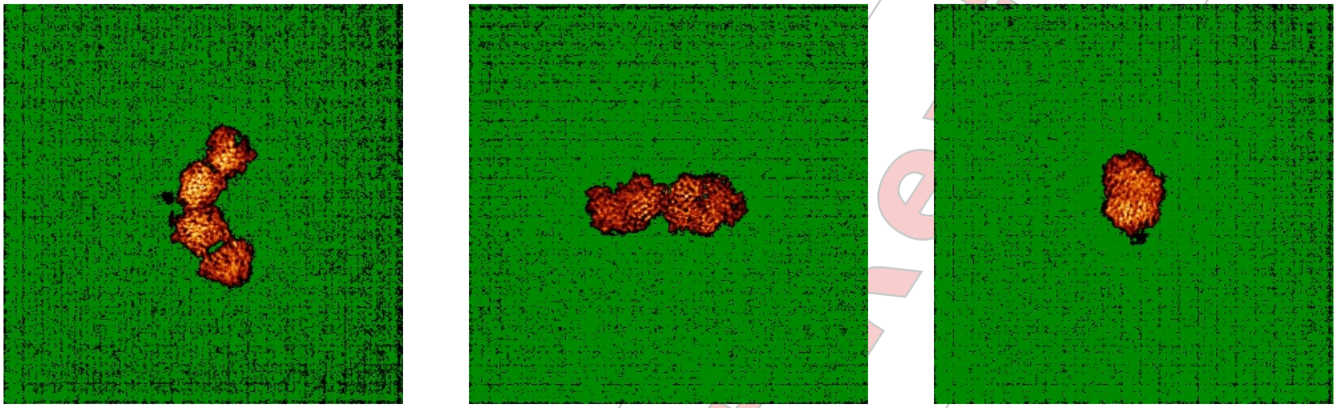

X Y Z

##### 6.4.2 Raw map

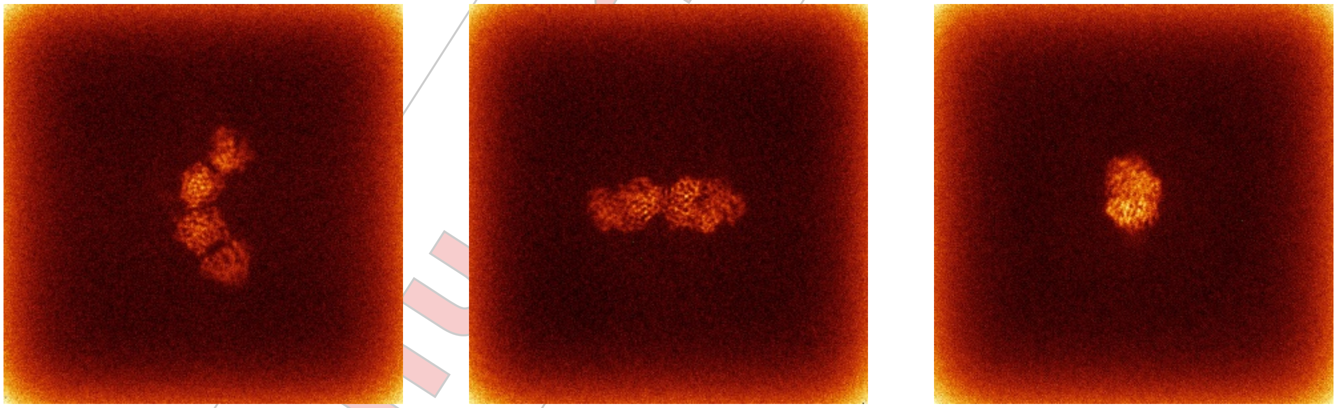

X Y Z

The images above show the map standard deviation projections with false color in three orthogonal directions. Minimum values are shown in green, max in blue, and dark to light orange shades represent small to large values respectively.

For Manuscript Review

#### 6.5 Orthogonal surface views (i)

##### 6.5.1 Primary map

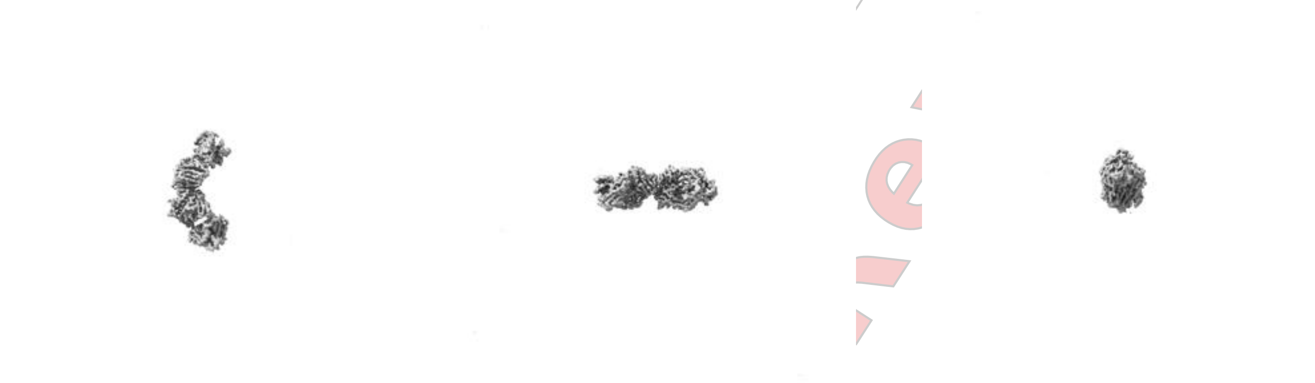

X Y Z

The images above show the 3D surface view of the map at the recommended contour level 0.12. These images, in conjunction with the slice images, may facilitate assessment of whether an appropriate contour level has been provided.

##### 6.5.2 Raw map

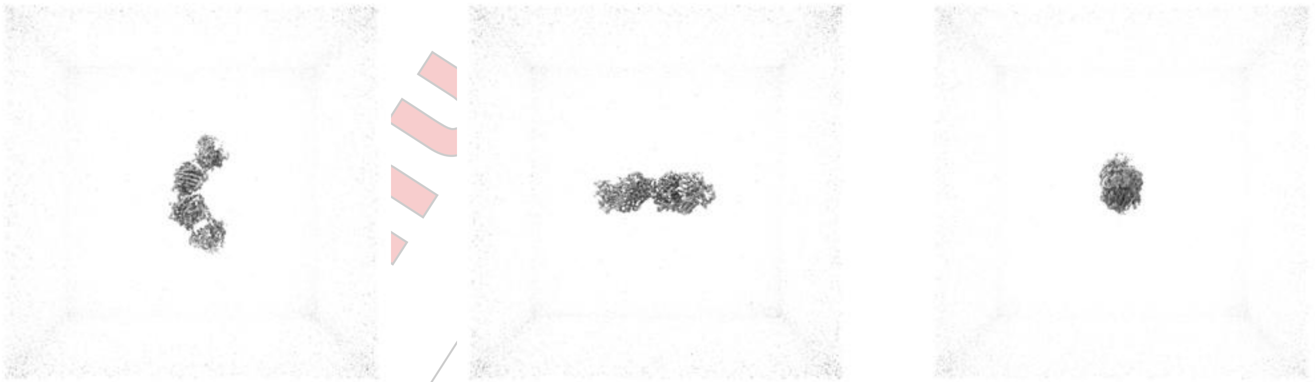

X Y Z

These images show the 3D surface of the raw map. The raw map's contour level was selected so that its surface encloses the same volume as the primary map does at its recommended contour level.

#### 6.6 Mask visualisation (i)

This section was not generated. No masks/segmentation were deposited.

For Manuscript Review

### 7 Map analysis (i)

This section contains the results of statistical analysis of the map.

#### 7.1 Map-value distribution (i)

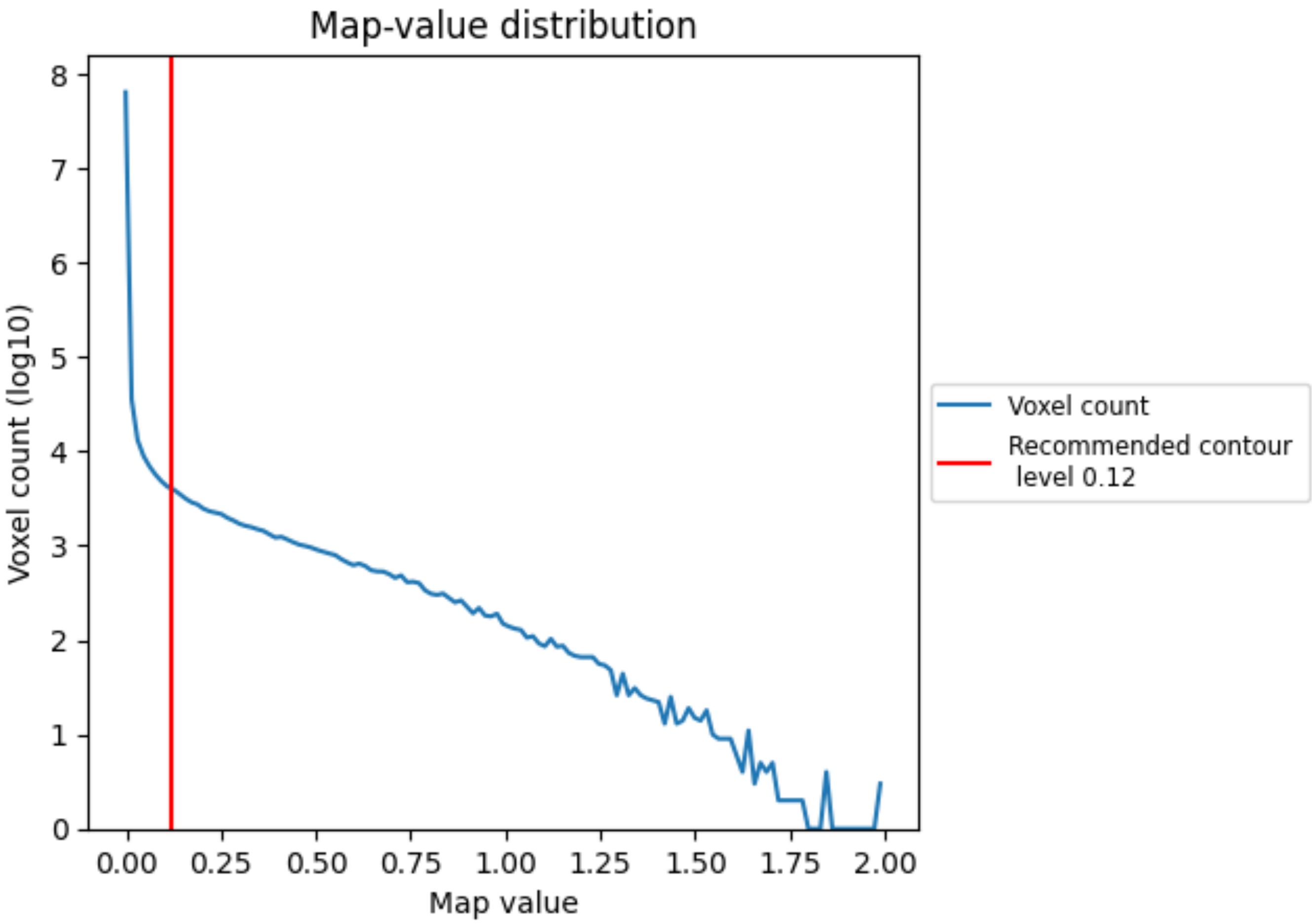


The map-value distribution is plotted in 128 intervals along the x-axis. The y-axis is logarithmic. A spike in this graph at zero usually indicates that the volume has been masked.

For Manuscript Review

#### 7.2 Volume estimate (i)

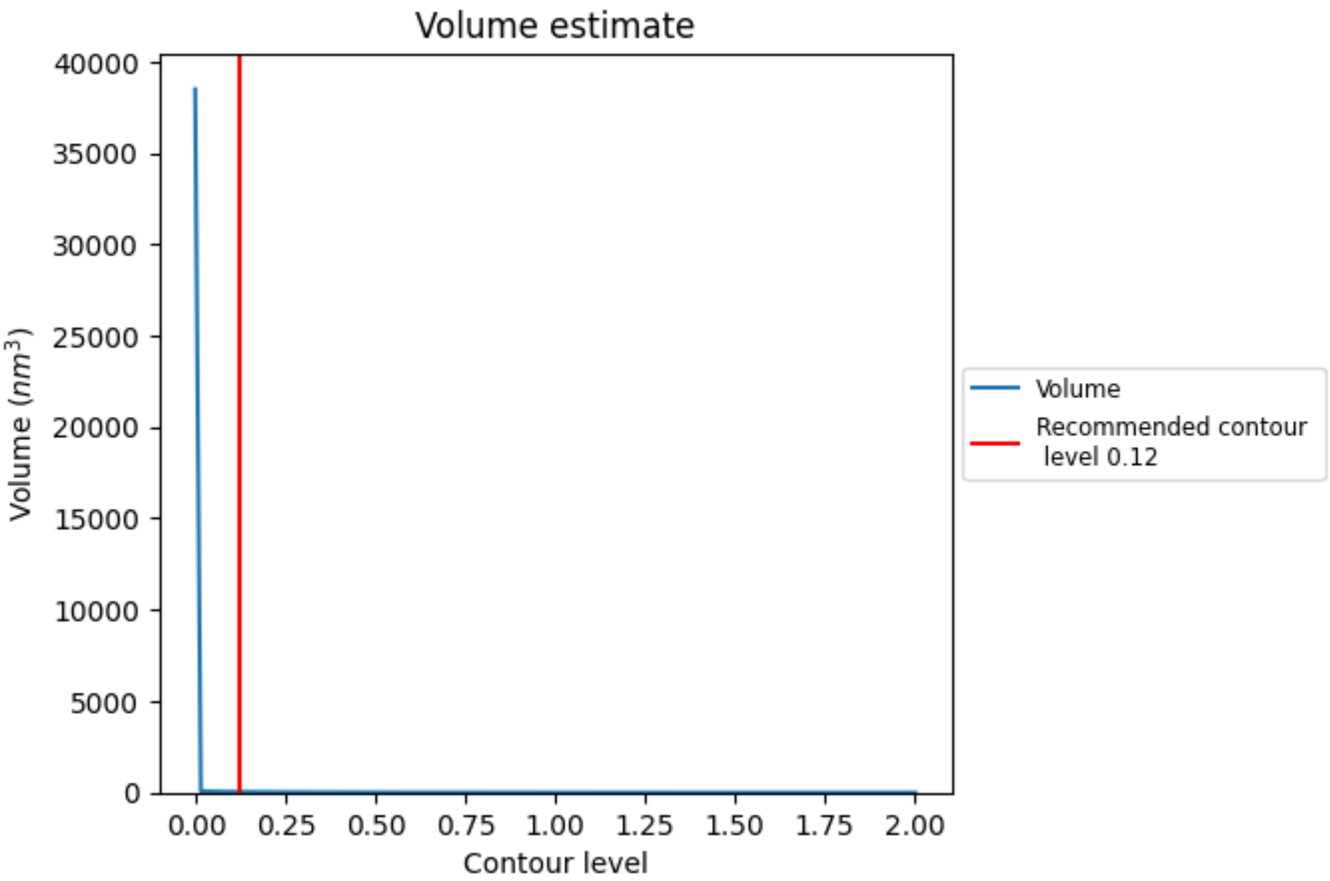


The volume at the recommended contour level is 38 $nm^3$; this corresponds to an approximate mass of 35 kDa.

The volume estimate graph shows how the enclosed volume varies with the contour level. The recommended contour level is shown as a vertical line and the intersection between the line and the curve gives the volume of the enclosed surface at the given level.

For Manuscript Review

#### 7.3 Rotationally averaged power spectrum (i)

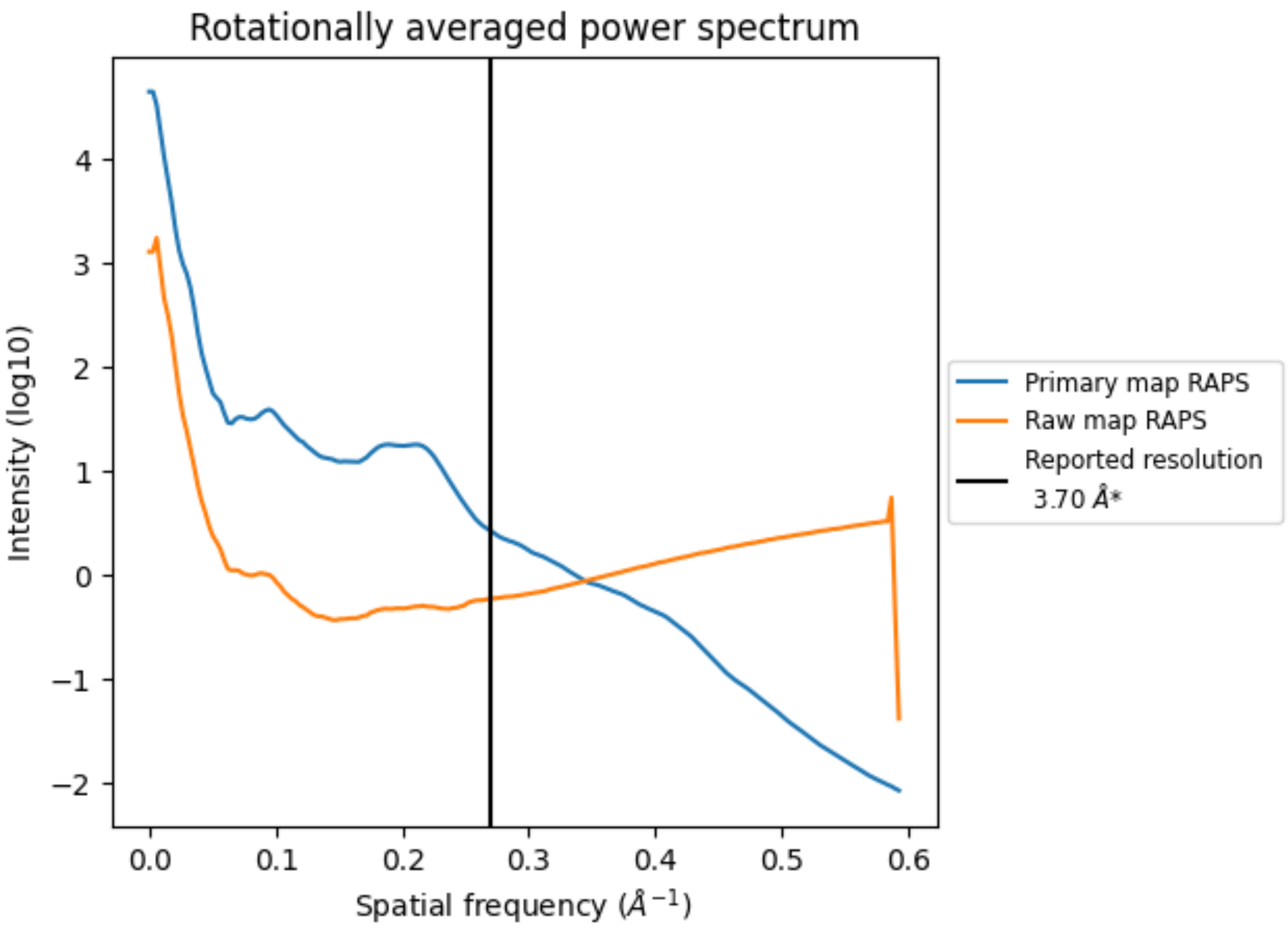


*Reported resolution corresponds to spatial frequency of 0.270 Å$^{-1}$

For Manuscript Review

### 8 Fourier-Shell correlation (i)

Fourier-Shell Correlation (FSC) is the most commonly used method to estimate the resolution of single-particle and subtomogram-averaged maps. The shape of the curve depends on the imposed symmetry, mask and whether or not the two 3D reconstructions used were processed from a common reference. The reported resolution is shown as a black line. A curve is displayed for the half-bit criterion in addition to lines showing the 0.143 gold standard cut-off and 0.5 cut-off.

#### 8.1 FSC (i)

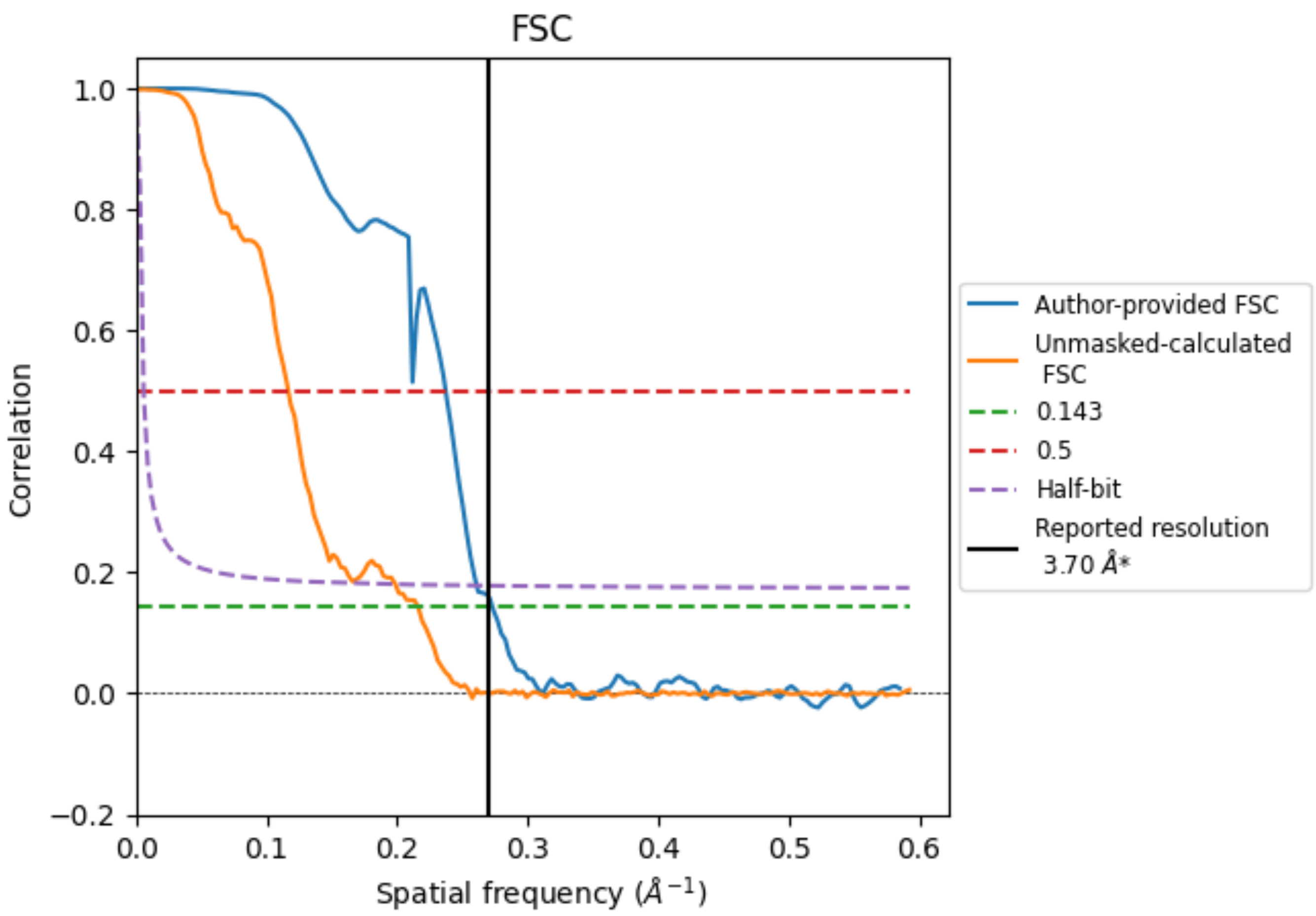


*Reported resolution corresponds to spatial frequency of 0.270 Å$^{-1}$

For Manuscript Review

#### 8.2 Resolution estimates (i)

| Resolution estimate (Å) | Estimation criterion (FSC cut-off) | | |
|---|---|---|---|
| | 0.143 | 0.5 | Half-bit |
| Reported by author | 3.70 | - | - |
| Author-provided FSC curve | 3.66 | 4.22 | 3.83 |
| Unmasked-calculated* | 4.63 | 8.58 | 5.04 |

*Resolution estimate based on FSC curve calculated by comparison of deposited half-maps. The value from deposited half-maps intersecting FSC 0.143 CUT-OFF 4.63 differs from the reported value 3.7 by more than 10 %

For Manuscript Review

### 9 Map-model fit (i)

This section contains information regarding the fit between EMDB map EMD-76196 and PDB model 11YS. Per-residue inclusion information can be found in section 3 on page 5.

#### 9.1 Map-model overlay (i)

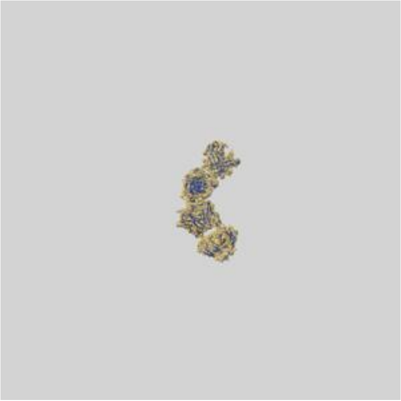

X

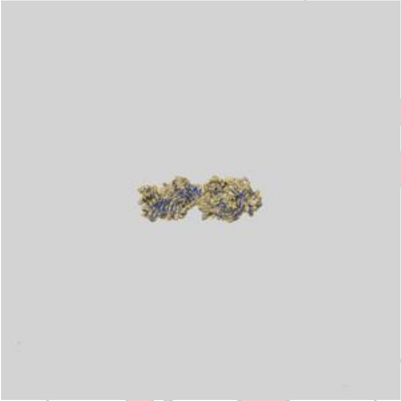

Y

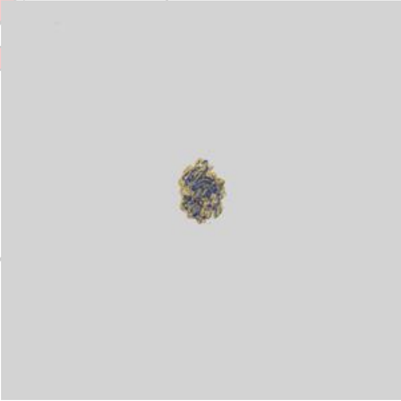

Z

The images above show the 3D surface view of the map at the recommended contour level 0.12 at 50% transparency in yellow overlaid with a ribbon representation of the model coloured in blue. These images allow for the visual assessment of the quality of fit between the atomic model and the map.

For Manuscript Review

#### 9.2 Q-score mapped to coordinate model (i)

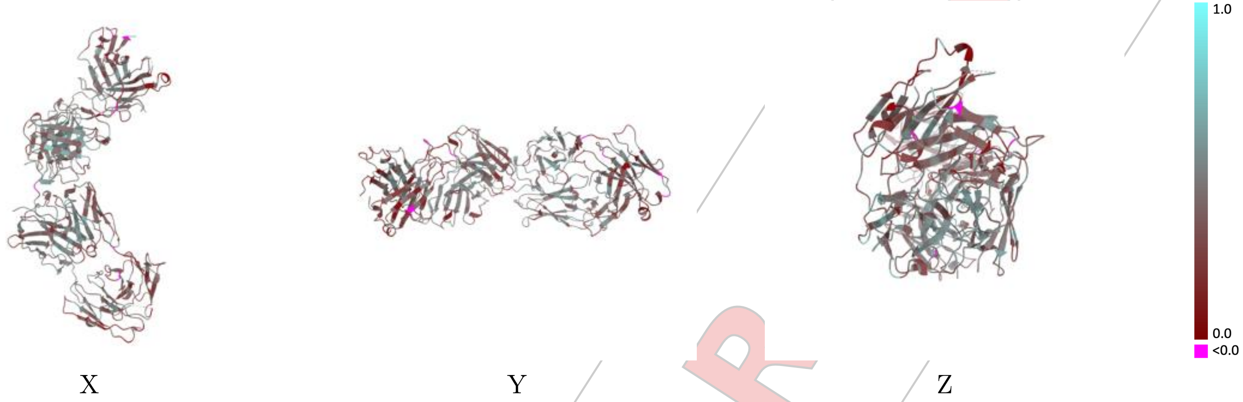


The images above show the model with each residue coloured according its Q-score. This shows their resolvability in the map with higher Q-score values reflecting better resolvability. Please note: Q-score is calculating the resolvability of atoms, and thus high values are only expected at resolutions at which atoms can be resolved. Low Q-score values may therefore be expected for many entries.

#### 9.3 Atom inclusion mapped to coordinate model (i)

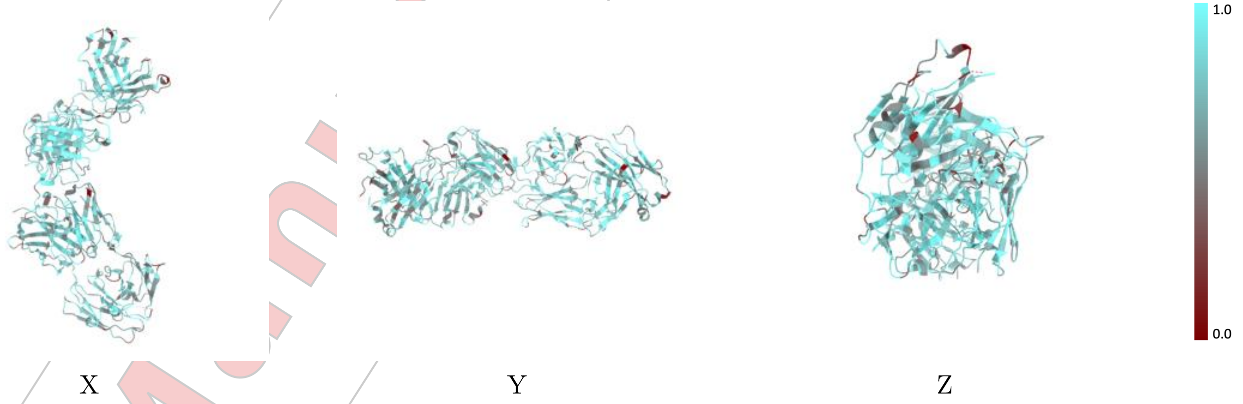


The images above show the model with each residue coloured according to its atom inclusion. This shows to what extent they are inside the map at the recommended contour level (0.12).

For Manuscript Review

#### 9.4 Atom inclusion (i)

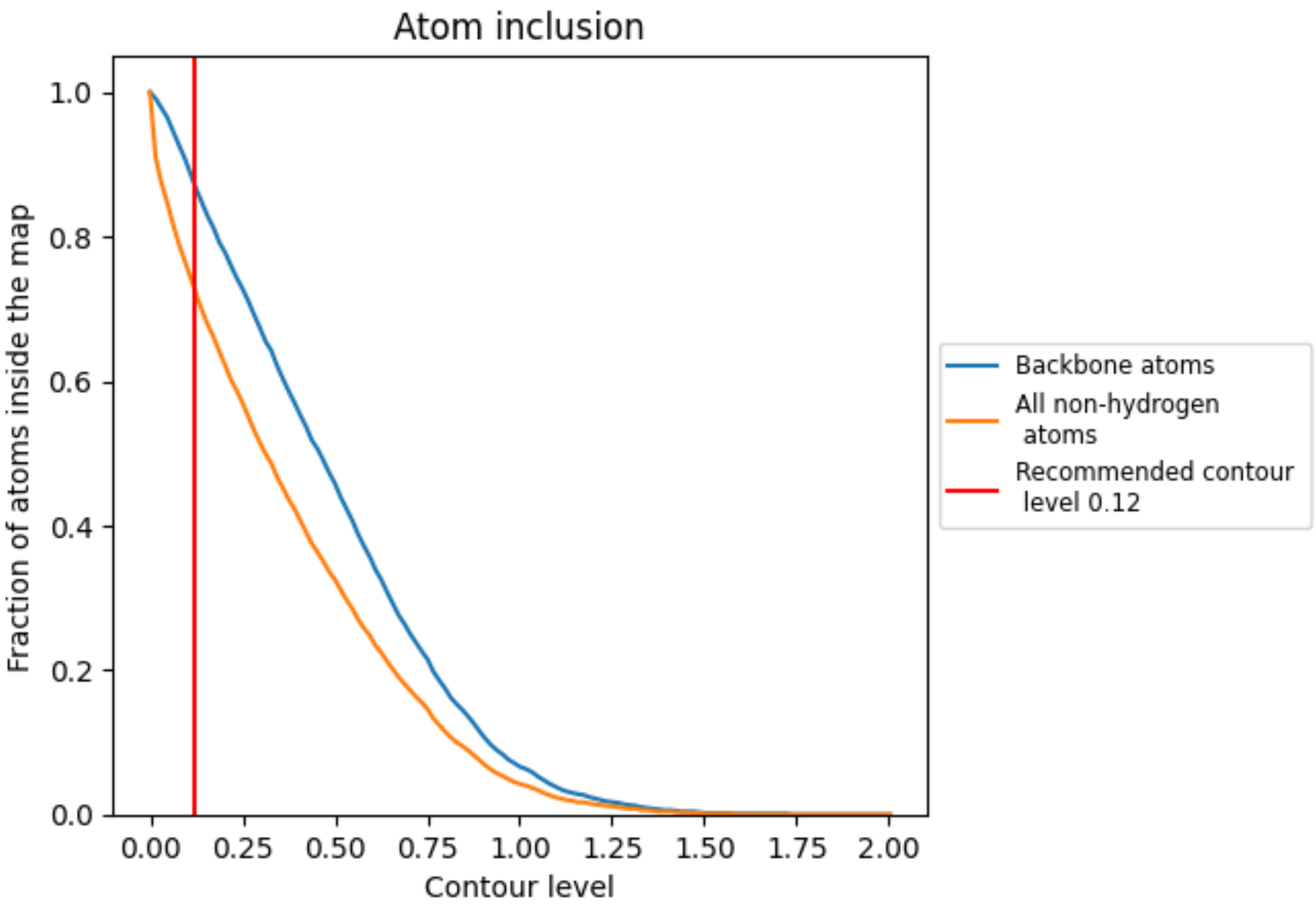


At the recommended contour level, 87% of all backbone atoms, 73% of all non-hydrogen atoms, are inside the map.

For Manuscript Review

#### 9.5 Map-model fit summary (i)

The table lists the average atom inclusion at the recommended contour level (0.12) and Q-score for the entire model and for each chain.

| Chain | Atom inclusion | Q-score |
|---|---|---|
| All | 0.7260 | 0.3910 |
| A | 0.7870 | 0.4400 |
| H | 0.7550 | 0.4170 |
| K | 0.6880 | 0.3460 |
| L | 0.7320 | 0.4010 |
| M | 0.7230 | 0.3960 |

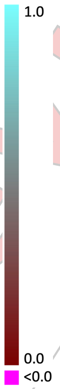


For Manuscript Review

### Full wwPDB EM Validation Report (i)

Sep 17, 2026 – 02:22 PM EDT

| | | |
|---|---|---|
| PDB ID | : | 13JR / pdb_000013jr |
| EMDB ID | : | EMD-77097 |
| Title | : | Cryo-EM structure of Ara h 2 peanut allergen in complex with P1BC6 and P1EC5 Fab fragments |
| Deposited on | : | 2026-05-11 |
| Resolution | : | 3.52 Å(reported) |

**This wwPDB validation report is for manuscript review**

This is a Full wwPDB EM Validation Report.

This report is produced by the wwPDB biocuration pipeline after annotation of the structure.

We welcome your comments at **
A user guide is available at
https://www.wwpdb.org/validation/2017/EMValidationReportHelp
with specific help available everywhere you see the (i) symbol.

The types of validation reports are described at
http://www.wwpdb.org/validation/2017/FAQs#types.

The following versions of software and data (see references (i)) were used in the production of this report:

| | | |
|---|---|---|
| EMDB validation analysis | : | 0.0.1.dev133 |
| MolProbity | : | 4-5-2 with Phenix2.0 |
| Percentile statistics | : | 20250101.v01 (using entries in the PDB archive January 1st 2025) |
| EM percentile statistics | : | 202505.v01 (Using data in the EMDB archive up until May 2025) |
| MapQ | : | 1.9.13 |
| Ideal geometry (proteins) | : | Engh & Huber (2001) |
| Ideal geometry (DNA, RNA) | : | Parkinson et al. (1996) |
| Validation Pipeline (wwPDB-VP) | : | 2.50 |

For Manuscript Review

### 1 Overall quality at a glance (i)

The following experimental techniques were used to determine the structure:
*ELECTRON MICROSCOPY*

The reported resolution of this entry is 3.52 Å.

Percentile scores (ranging between 0-100) for global validation metrics of the entry are shown in the following graphic. The table shows the number of entries on which the scores are based.

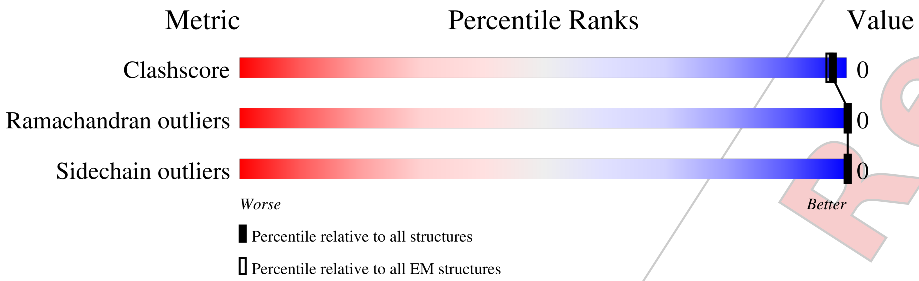


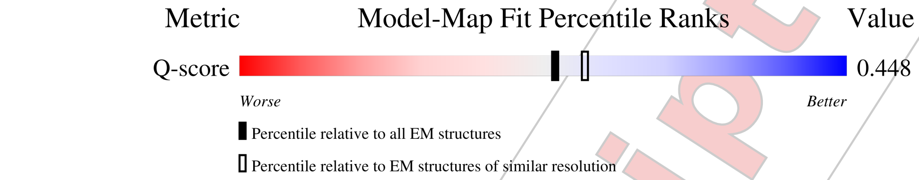


| Metric | Whole archive (#Entries) | EM structures (#Entries) | Similar EM resolution (#Entries, resolution range(Å)) |
|---|---|---|---|
| Clashscore | 229148 | 23984 | - |
| Ramachandran outliers | 224038 | 23583 | - |
| Sidechain outliers | 223484 | 23102 | - |
| Q-score | - | 25397 | 13017 ( 3.02 - 4.02 ) |

The table below summarises the geometric issues observed across the polymeric chains and their fit to the map. The red, orange, yellow and green segments of the bar indicate the fraction of residues that contain outliers for >=3, 2, 1 and 0 types of geometric quality criteria respectively. A grey segment represents the fraction of residues that are not modelled. The numeric value for each fraction is indicated below the corresponding segment, with a dot representing fractions <=5% The upper red bar (where present) indicates the fraction of residues that have poor fit to the EM map (all-atom inclusion < 40%). The numeric value is given above the bar.

| Mol | Chain | Length | Quality of chain |
|---|---|---|---|
| 1 | A | 220 | 7% / 98% • |
| 2 | B | 218 | 5% / 98% •• |
| 3 | C | 229 | 7% / 97% •• |

*Continued on next page...*

For Manuscript Review

*Continued from previous page...*

| Mol | Chain | Length | Quality of chain |
|---|---|---|---|
| 4 | D | 212 | 7% 95% • • |
| 5 | E | 151 | 7% 62% 38% |

For Manuscript Review

### 2 Entry composition (i)

There are 5 unique types of molecules in this entry. The entry contains 7322 atoms, of which 0 are hydrogens and 0 are deuteriums.

In the tables below, the AltConf column contains the number of residues with at least one atom in alternate conformation and the Trace column contains the number of residues modelled with at most 2 atoms.

- Molecule 1 is a protein called P1BC6 Fab heavy chain.

| Mol | Chain | Residues | Atoms | | | | | AltConf | Trace |
|---|---|---|---|---|---|---|---|---|---|
| | | | Total | C | N | O | S | | |
| 1 | A | 216 | 1614 | 1013 | 280 | 314 | 7 | 0 | 0 |

- Molecule 2 is a protein called P1BC6 Fab light chain.

| Mol | Chain | Residues | Atoms | | | | | AltConf | Trace |
|---|---|---|---|---|---|---|---|---|---|
| | | | Total | C | N | O | S | | |
| 2 | B | 216 | 1667 | 1046 | 278 | 339 | 4 | 0 | 0 |

- Molecule 3 is a protein called P1EC5 Fab heavy chain.

| Mol | Chain | Residues | Atoms | | | | | AltConf | Trace |
|---|---|---|---|---|---|---|---|---|---|
| | | | Total | C | N | O | S | | |
| 3 | C | 224 | 1678 | 1057 | 288 | 325 | 8 | 0 | 0 |

- Molecule 4 is a protein called P1EC5 Fab light chain.

| Mol | Chain | Residues | Atoms | | | | | AltConf | Trace |
|---|---|---|---|---|---|---|---|---|---|
| | | | Total | C | N | O | S | | |
| 4 | D | 210 | 1611 | 1002 | 271 | 332 | 6 | 0 | 0 |

- Molecule 5 is a protein called Conglutin-7.

| Mol | Chain | Residues | Atoms | | | | | AltConf | Trace |
|---|---|---|---|---|---|---|---|---|---|
| | | | Total | C | N | O | S | | |
| 5 | E | 94 | 752 | 447 | 151 | 143 | 11 | 0 | 0 |

For Manuscript Review

### 3 Residue-property plots ⓘ

These plots are drawn for all protein, RNA, DNA and oligosaccharide chains in the entry. The first graphic for a chain summarises the proportions of the various outlier classes displayed in the second graphic. The second graphic shows the sequence view annotated by issues in geometry and atom inclusion in map density. Residues are color-coded according to the number of geometric quality criteria for which they contain at least one outlier: green = 0, yellow = 1, orange = 2 and red = 3 or more. A red diamond above a residue indicates a poor fit to the EM map for this residue (all-atom inclusion < 40%). Stretches of 2 or more consecutive residues without any outlier are shown as a green connector. Residues present in the sample, but not in the model, are shown in grey.

- Molecule 1: P1BC6 Fab heavy chain

Chain A:

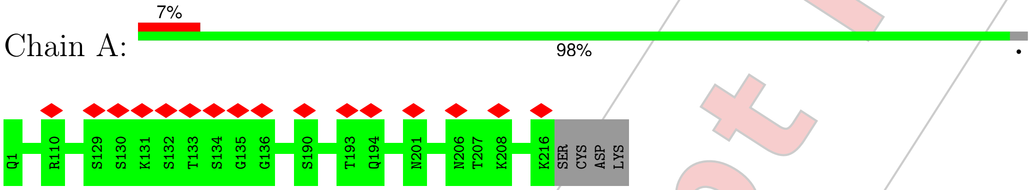


- Molecule 2: P1BC6 Fab light chain

Chain B:

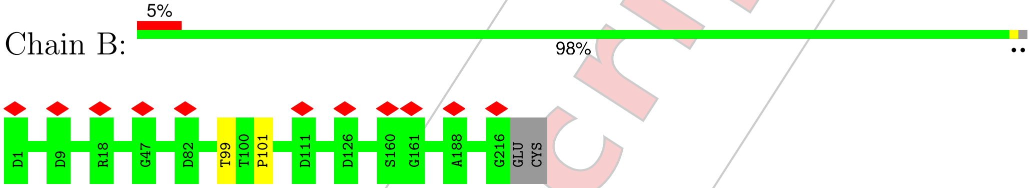


- Molecule 3: P1EC5 Fab heavy chain

Chain C:

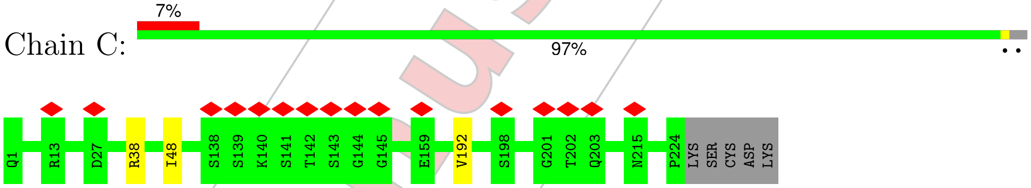


- Molecule 4: P1EC5 Fab light chain

Chain D:

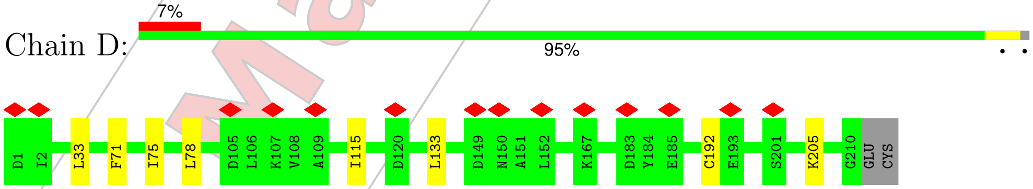


- Molecule 5: Conglutin-7

Chain E:

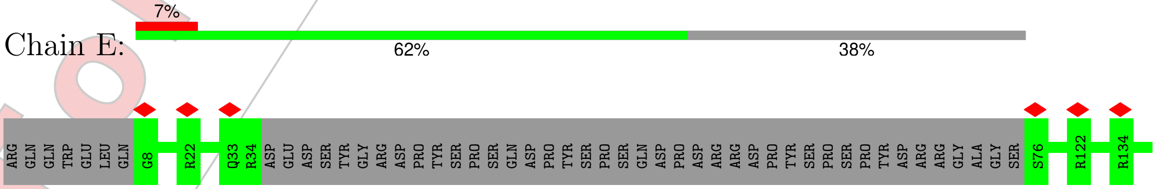


For Manuscript Review

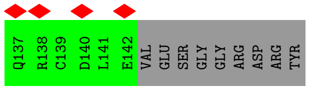


For Manuscript Review

### 4 Experimental information (i)

| Property | Value | Source |
|---|---|---|
| EM reconstruction method | SINGLE PARTICLE | Depositor |
| Imposed symmetry | POINT, C1 | Depositor |
| Number of particles used | 211894 | Depositor |
| Resolution determination method | FSC 0.143 CUT-OFF | Depositor |
| CTF correction method | PHASE FLIPPING AND AMPLITUDE CORRECTION | Depositor |
| Microscope | TFS KRIOS | Depositor |
| Voltage (kV) | 300 | Depositor |
| Electron dose $(e^-/\text{Å}^2)$ | 55.24 | Depositor |
| Minimum defocus (nm) | 1500 | Depositor |
| Maximum defocus (nm) | 2500 | Depositor |
| Magnification | Not provided | |
| Image detector | GATAN K3 BIOQUANTUM (6k x 4k) | Depositor |
| Maximum map value | 1.879 | Depositor |
| Minimum map value | -0.002 | Depositor |
| Average map value | 0.000 | Depositor |
| Map value standard deviation | 0.011 | Depositor |
| Recommended contour level | 0.12 | Depositor |
| Map size (Å) | 396.47998, 396.47998, 396.47998 | wwPDB |
| Map dimensions | 480, 480, 480 | wwPDB |
| Map angles (°) | 90.0, 90.0, 90.0 | wwPDB |
| Pixel spacing (Å) | 0.826, 0.826, 0.826 | Depositor |

For Manuscript Review

### 5 Model quality (i)

#### 5.1 Standard geometry (i)

The Z score for a bond length (or angle) is the number of standard deviations the observed value is removed from the expected value. A bond length (or angle) with $|Z| > 5$ is considered an outlier worth inspection. RMSZ is the root-mean-square of all Z scores of the bond lengths (or angles).

| Mol | Chain | Bond lengths | | Bond angles | |
|---|---|---|---|---|---|
| | | RMSZ | #$\|Z\|$ >5 | RMSZ | #$\|Z\|$ >5 |
| 1 | A | 0.20 | 0/1651 | 0.34 | 0/2247 |
| 2 | B | 0.26 | 0/1706 | 0.41 | 0/2320 |
| 3 | C | 0.21 | 0/1721 | 0.38 | 0/2347 |
| 4 | D | 0.24 | 0/1644 | 0.40 | 0/2233 |
| 5 | E | 0.22 | 0/757 | 0.38 | 0/1008 |
| All | All | 0.23 | 0/7479 | 0.38 | 0/10155 |

There are no bond length outliers.

There are no bond angle outliers.

There are no chirality outliers.

There are no planarity outliers.

#### 5.2 Too-close contacts (i)

In the following table, the Non-H and H(model) columns list the number of non-hydrogen atoms and hydrogen atoms in the chain respectively. The H(added) column lists the number of hydrogen atoms added and optimized by MolProbity. The Clashes column lists the number of clashes within the asymmetric unit, whereas Symm-Clashes lists symmetry-related clashes.

| Mol | Chain | Non-H | H(model) | H(added) | Clashes | Symm-Clashes |
|---|---|---|---|---|---|---|
| 1 | A | 1614 | 0 | 1597 | 0 | 0 |
| 2 | B | 1667 | 0 | 1596 | 1 | 0 |
| 3 | C | 1678 | 0 | 1642 | 2 | 0 |
| 4 | D | 1611 | 0 | 1555 | 5 | 0 |
| 5 | E | 752 | 0 | 694 | 0 | 0 |
| All | All | 7322 | 0 | 7084 | 7 | 0 |

The all-atom clashscore is defined as the number of clashes found per 1000 atoms (including hydrogen atoms). The all-atom clashscore for this structure is 0.

All (7) close contacts within the same asymmetric unit are listed below, sorted by their clash magnitude.

For Manuscript Review

| Atom-1 | Atom-2 | Interatomic distance (Å) | Clash overlap (Å) |
|---|---|---|---|
| 3:C:38:ARG:HB2 | 3:C:48:ILE:HD11 | 1.90 | 0.52 |
| 4:D:75:ILE:HG21 | 4:D:78:LEU:HD12 | 1.94 | 0.49 |
| 4:D:115:ILE:HD12 | 4:D:205:LYS:C | 2.41 | 0.46 |
| 4:D:115:ILE:HD11 | 4:D:192:CYS:HB3 | 1.98 | 0.45 |
| 4:D:33:LEU:HD22 | 4:D:71:PHE:CD2 | 2.52 | 0.45 |
| 3:C:192:VAL:HG21 | 4:D:133:LEU:CD2 | 2.48 | 0.43 |
| 2:B:99:THR:HG22 | 2:B:101:PRO:HD2 | 2.02 | 0.41 |

There are no symmetry-related clashes.

#### 5.3 Torsion angles (i)

##### 5.3.1 Protein backbone (i)

In the following table, the Percentiles column shows the percent Ramachandran outliers of the chain as a percentile score with respect to all PDB entries followed by that with respect to all EM entries.

The Analysed column shows the number of residues for which the backbone conformation was analysed, and the total number of residues.

| Mol | Chain | Analysed | Favoured | Allowed | Outliers | Percentiles | |
|---|---|---|---|---|---|---|---|
| 1 | A | 214/220 (97%) | 203 (95%) | 11 (5%) | 0 | 100 | 100 |
| 2 | B | 214/218 (98%) | 201 (94%) | 13 (6%) | 0 | 100 | 100 |
| 3 | C | 222/229 (97%) | 211 (95%) | 11 (5%) | 0 | 100 | 100 |
| 4 | D | 208/212 (98%) | 195 (94%) | 13 (6%) | 0 | 100 | 100 |
| 5 | E | 90/151 (60%) | 86 (96%) | 4 (4%) | 0 | 100 | 100 |
| All | All | 948/1030 (92%) | 896 (94%) | 52 (6%) | 0 | 100 | 100 |

There are no Ramachandran outliers to report.

##### 5.3.2 Protein sidechains (i)

In the following table, the Percentiles column shows the percent sidechain outliers of the chain as a percentile score with respect to all PDB entries followed by that with respect to all EM entries.

The Analysed column shows the number of residues for which the sidechain conformation was analysed, and the total number of residues.

For Manuscript Review

| Mol | Chain | Analysed | Rotameric | Outliers | Percentiles | |
|---|---|---|---|---|---|---|
| 1 | A | 180/184 (98%) | 180 (100%) | 0 | 100 | 100 |
| 2 | B | 188/190 (99%) | 188 (100%) | 0 | 100 | 100 |
| 3 | C | 189/194 (97%) | 189 (100%) | 0 | 100 | 100 |
| 4 | D | 185/187 (99%) | 185 (100%) | 0 | 100 | 100 |
| 5 | E | 78/139 (56%) | 78 (100%) | 0 | 100 | 100 |
| All | All | 820/894 (92%) | 820 (100%) | 0 | 100 | 100 |

There are no protein residues with a non-rotameric sidechain to report.

Sometimes sidechains can be flipped to improve hydrogen bonding and reduce clashes. All (6) such sidechains are listed below:

| Mol | Chain | Res | Type |
|---|---|---|---|
| 1 | A | 202 | HIS |
| 2 | B | 44 | HIS |
| 2 | B | 193 | HIS |
| 3 | C | 175 | HIS |
| 5 | E | 26 | GLN |
| 5 | E | 116 | GLN |

##### 5.3.3 RNA ⓘ

There are no RNA molecules in this entry.

#### 5.4 Non-standard residues in protein, DNA, RNA chains ⓘ

There are no non-standard protein/DNA/RNA residues in this entry.

#### 5.5 Carbohydrates ⓘ

There are no oligosaccharides in this entry.

#### 5.6 Ligand geometry ⓘ

There are no ligands in this entry.

#### 5.7 Other polymers ⓘ

There are no such residues in this entry.

For Manuscript Review

#### 5.8 Polymer linkage issues (i)

There are no chain breaks in this entry.

For Manuscript Review

### 6 Map visualisation (i)

This section contains visualisations of the EMDB entry EMD-77097. These allow visual inspection of the internal detail of the map and identification of artifacts.

Images derived from a raw map, generated by summing the deposited half-maps, are presented below the corresponding image components of the primary map to allow further visual inspection and comparison with those of the primary map.

#### 6.1 Orthogonal projections (i)

##### 6.1.1 Primary map

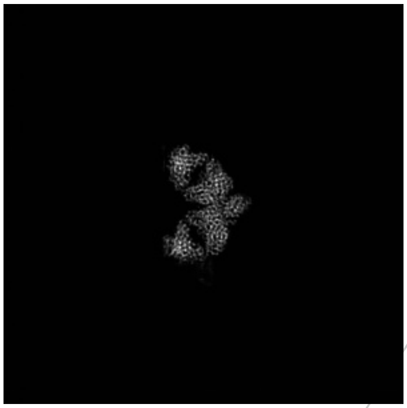

X

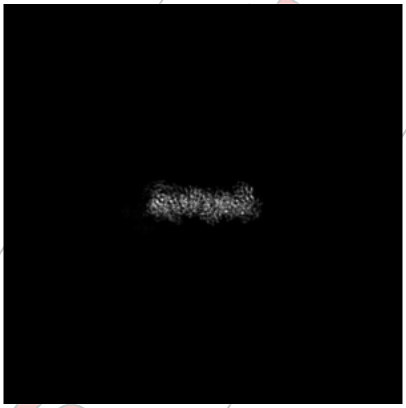

Y

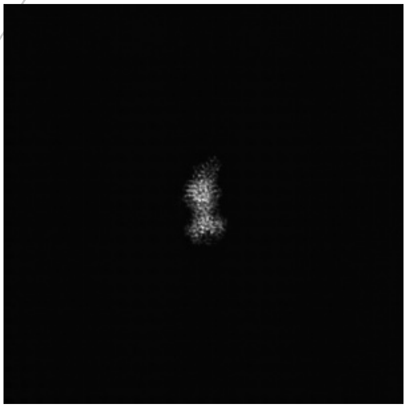

Z

##### 6.1.2 Raw map

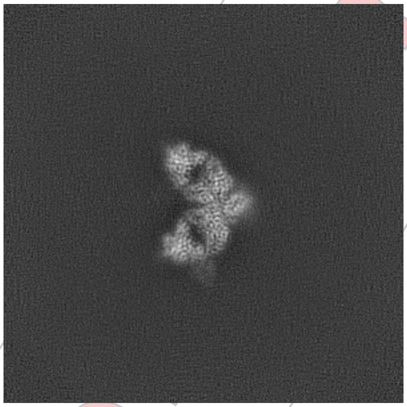

X

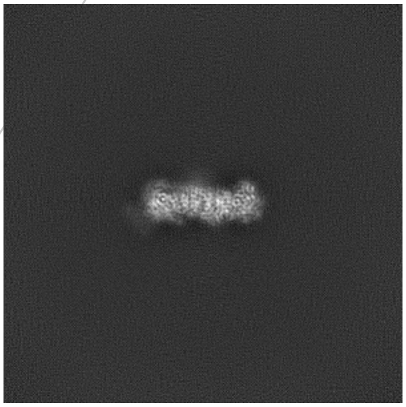

Y

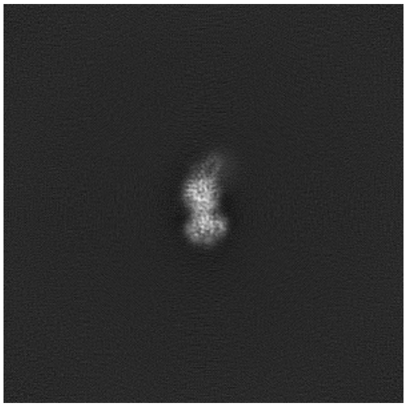

Z

The images above show the map projected in three orthogonal directions.

For Manuscript Review

#### 6.2 Central slices (i)

##### 6.2.1 Primary map

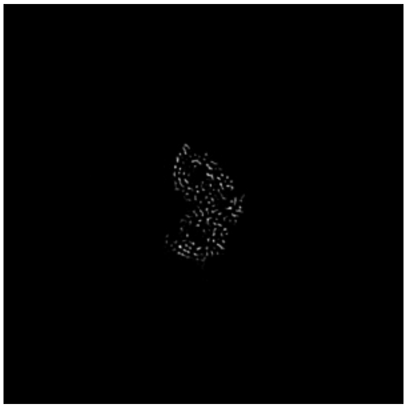

X Index: 240

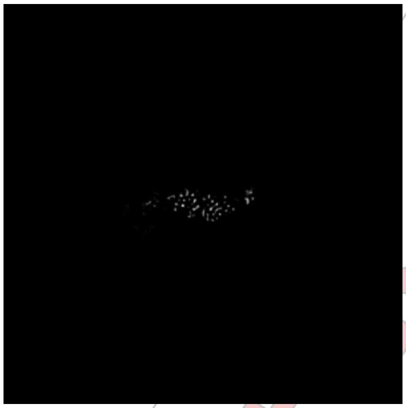

Y Index: 240

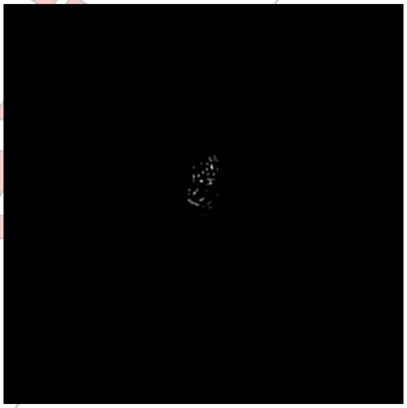

Z Index: 240

##### 6.2.2 Raw map

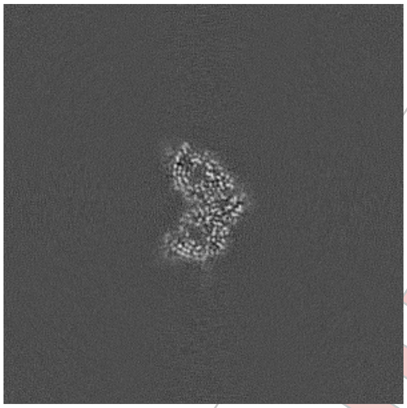

X Index: 240

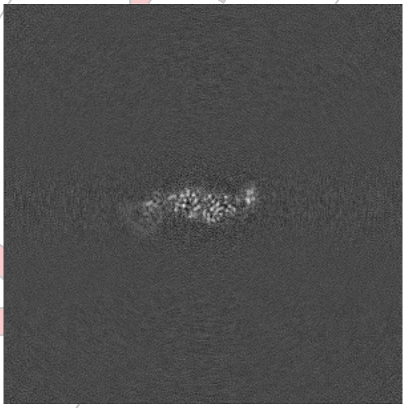

Y Index: 240

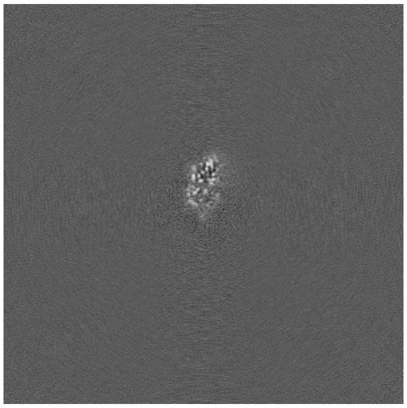

Z Index: 240

The images above show central slices of the map in three orthogonal directions.

For Manuscript Review

#### 6.3 Largest variance slices (i)

##### 6.3.1 Primary map

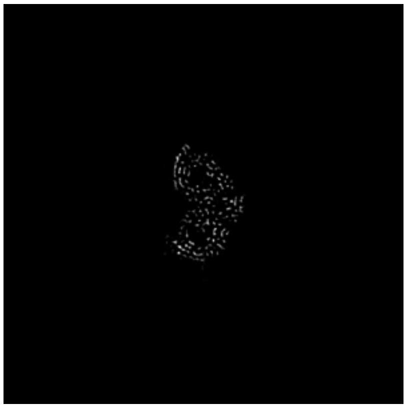
X Index: 239

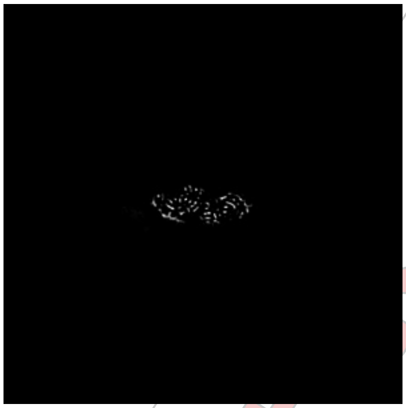
Y Index: 250

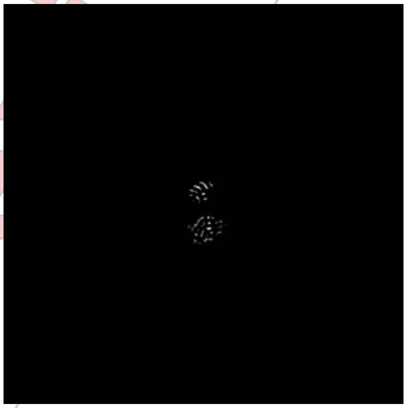
Z Index: 195

##### 6.3.2 Raw map

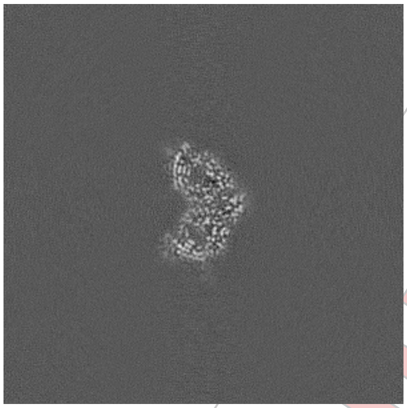
X Index: 239

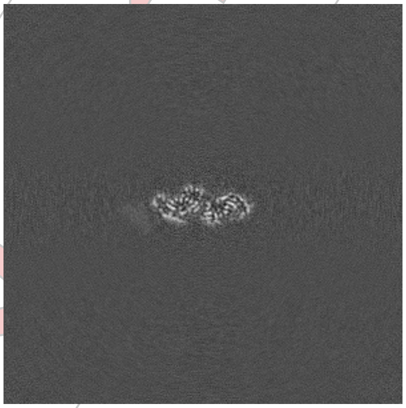
Y Index: 250

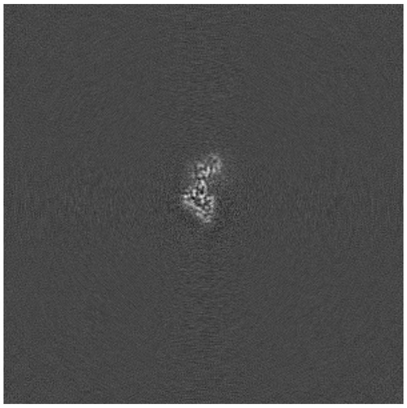
Z Index: 247

The images above show the largest variance slices of the map in three orthogonal directions.

For Manuscript Review

#### 6.4 Orthogonal standard-deviation projections (False-color)

##### 6.4.1 Primary map

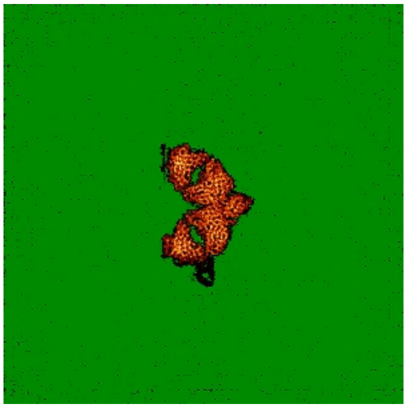

X

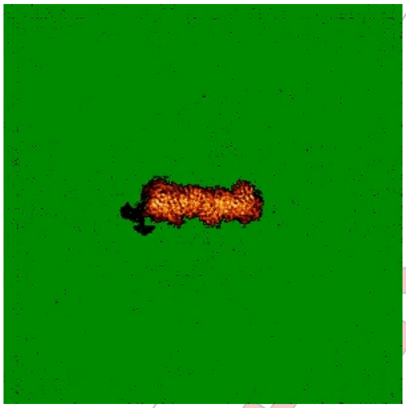

Y

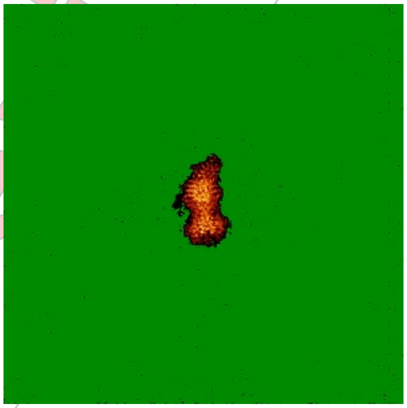

Z

##### 6.4.2 Raw map

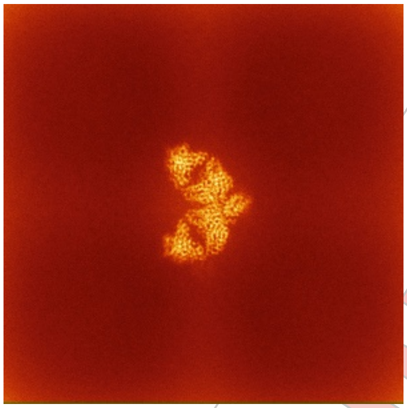

X

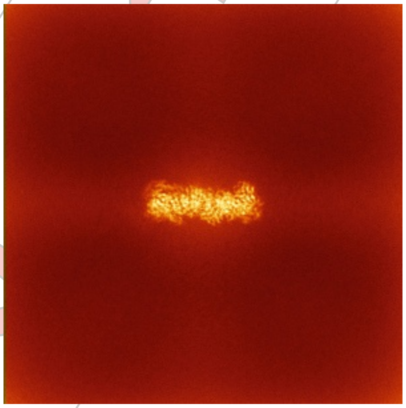

Y

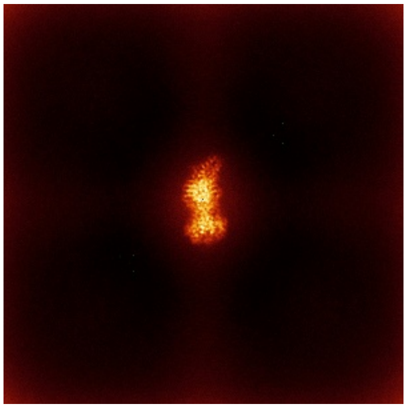

Z

The images above show the map standard deviation projections with false color in three orthogonal directions. Minimum values are shown in green, max in blue, and dark to light orange shades represent small to large values respectively.

For Manuscript Review

#### 6.5 Orthogonal surface views (i)

##### 6.5.1 Primary map

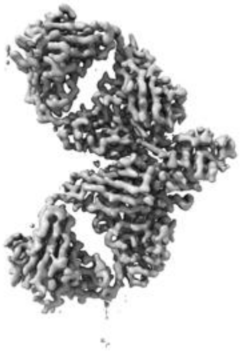

X

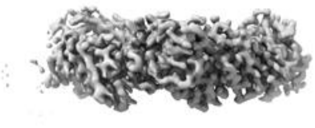

Y

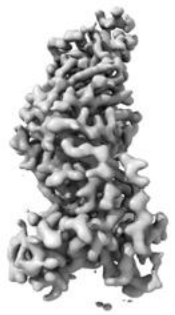

Z

The images above show the 3D surface view of the map at the recommended contour level 0.12. These images, in conjunction with the slice images, may facilitate assessment of whether an appropriate contour level has been provided.

##### 6.5.2 Raw map

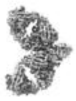

X

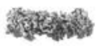

Y

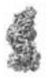

Z

These images show the 3D surface of the raw map. The raw map's contour level was selected so that its surface encloses the same volume as the primary map does at its recommended contour level.

#### 6.6 Mask visualisation (i)

This section was not generated. No masks/segmentation were deposited.

For Manuscript Review

### 7 Map analysis (i)

This section contains the results of statistical analysis of the map.

#### 7.1 Map-value distribution (i)

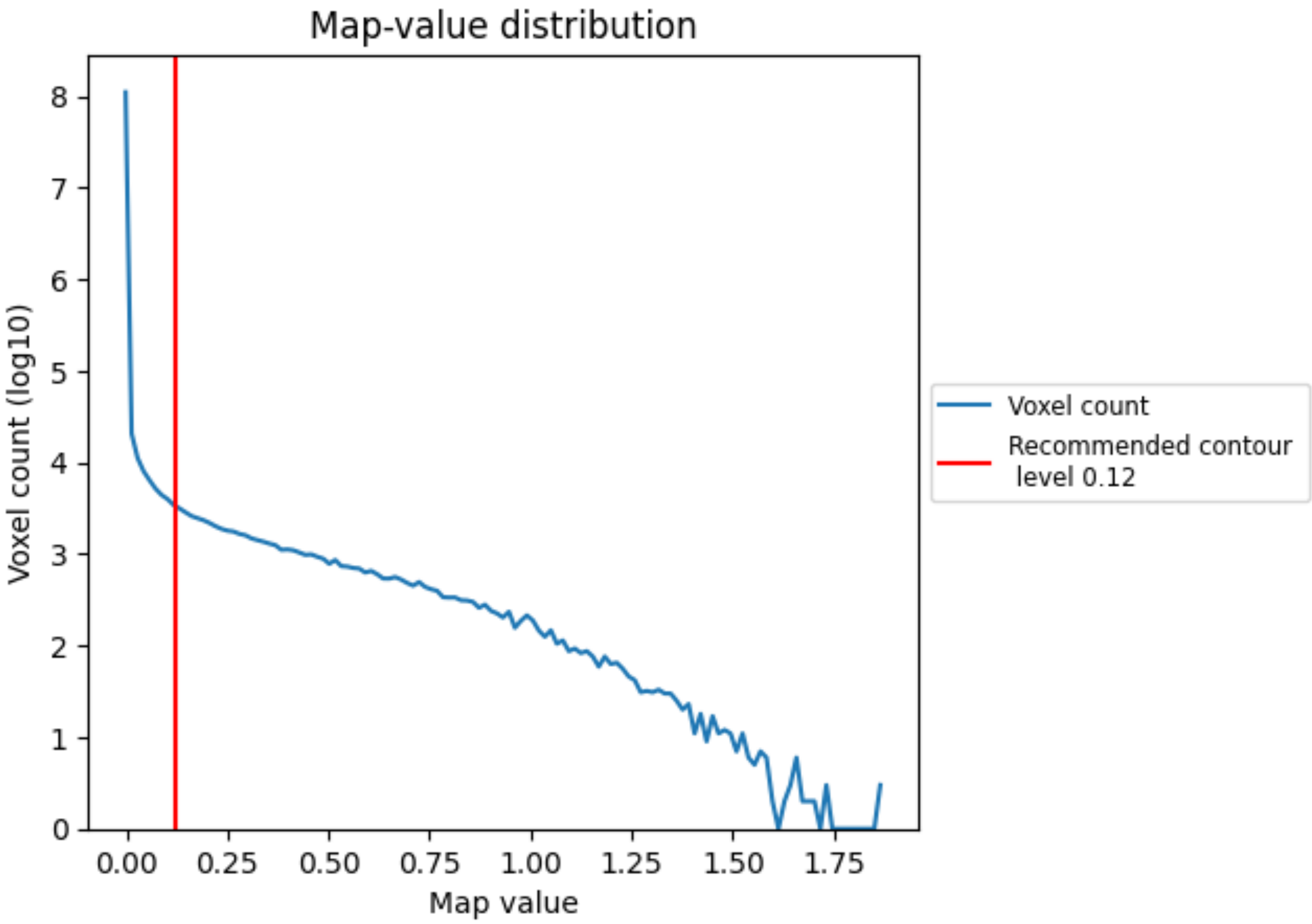


The map-value distribution is plotted in 128 intervals along the x-axis. The y-axis is logarithmic. A spike in this graph at zero usually indicates that the volume has been masked.

For Manuscript Review

#### 7.2 Volume estimate (i)

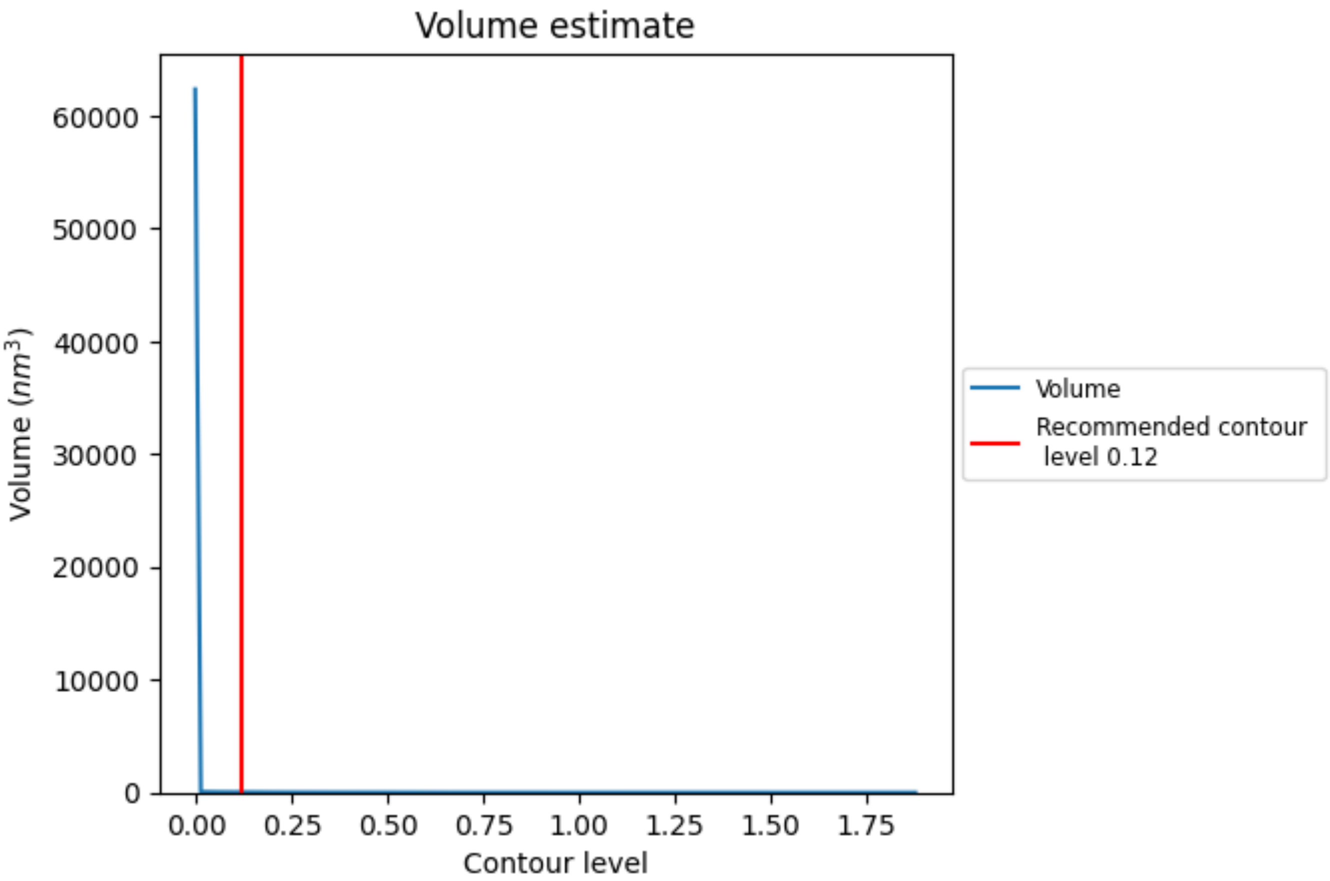


The volume at the recommended contour level is 35 $nm^3$; this corresponds to an approximate mass of 31 kDa.

The volume estimate graph shows how the enclosed volume varies with the contour level. The recommended contour level is shown as a vertical line and the intersection between the line and the curve gives the volume of the enclosed surface at the given level.

For Manuscript Review

#### 7.3 Rotationally averaged power spectrum (i)

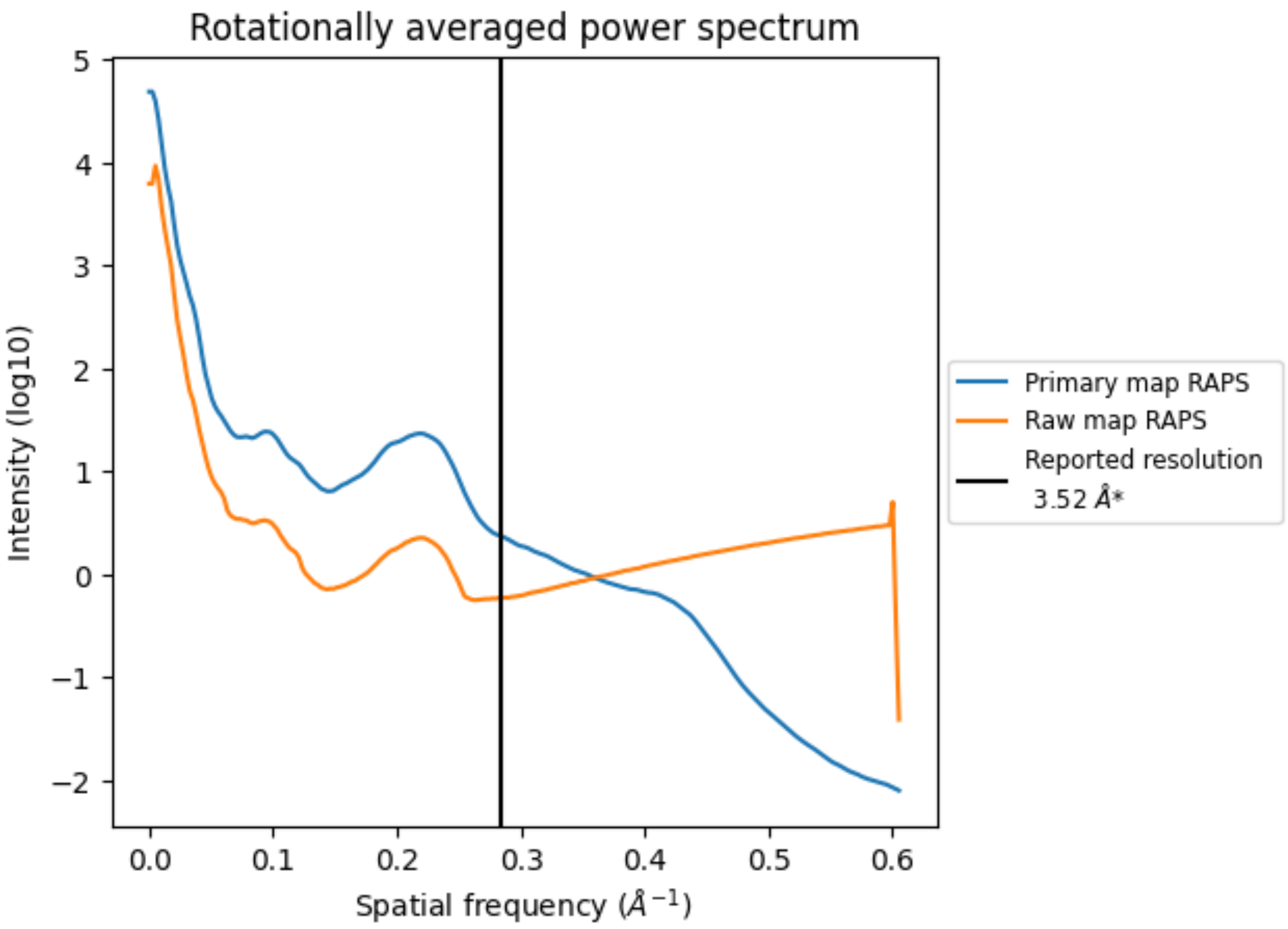


*Reported resolution corresponds to spatial frequency of 0.284 Å$^{-1}$

For Manuscript Review

### 8 Fourier-Shell correlation (i)

Fourier-Shell Correlation (FSC) is the most commonly used method to estimate the resolution of single-particle and subtomogram-averaged maps. The shape of the curve depends on the imposed symmetry, mask and whether or not the two 3D reconstructions used were processed from a common reference. The reported resolution is shown as a black line. A curve is displayed for the half-bit criterion in addition to lines showing the 0.143 gold standard cut-off and 0.5 cut-off.

#### 8.1 FSC (i)

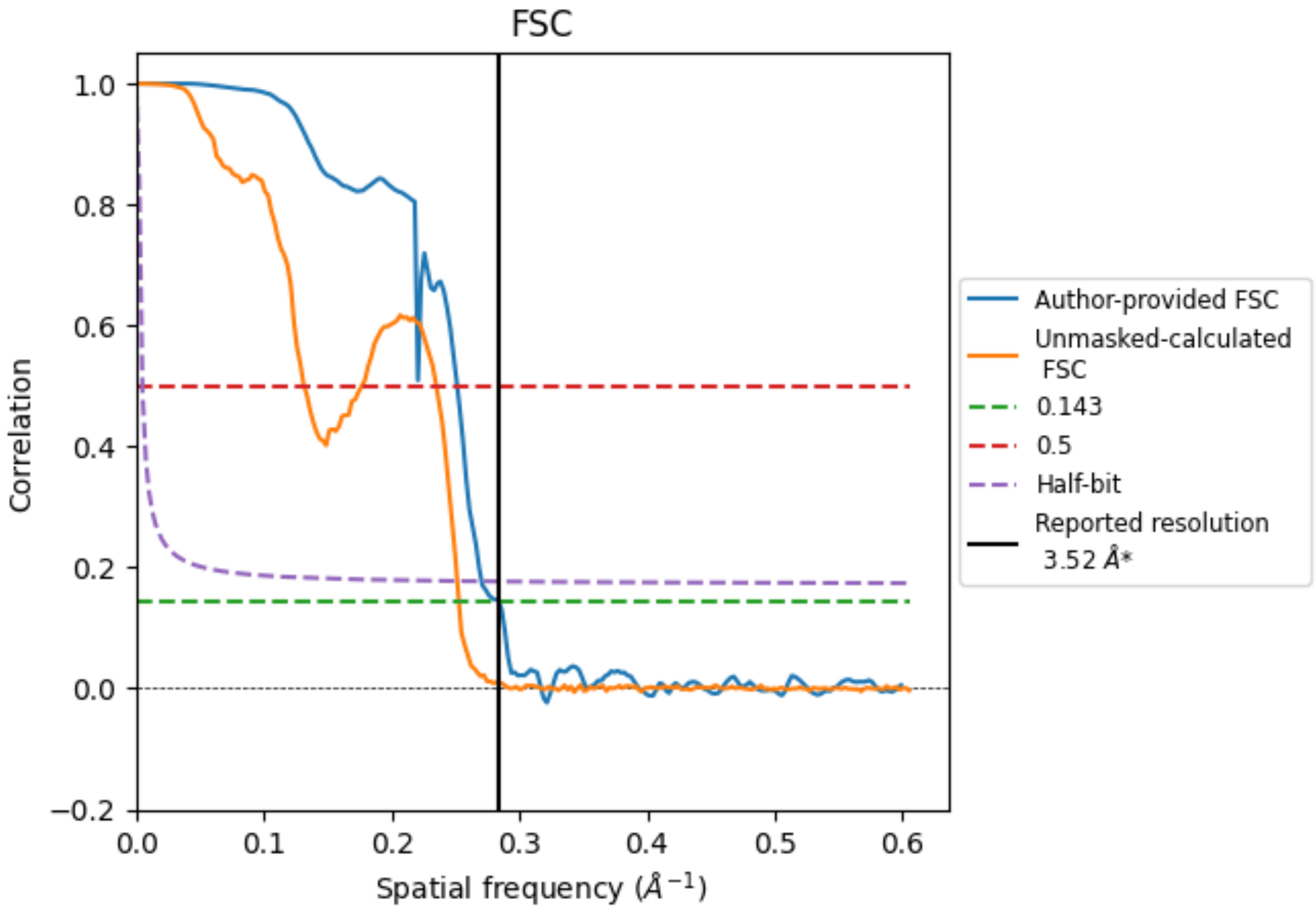


*Reported resolution corresponds to spatial frequency of 0.284 Å$^{-1}$

For Manuscript Review

#### 8.2 Resolution estimates (i)

| Resolution estimate (Å) | Estimation criterion (FSC cut-off) | | |
|---|---|---|---|
| | 0.143 | 0.5 | Half-bit |
| Reported by author | 3.52 | - | - |
| Author-provided FSC curve | 3.52 | 3.98 | 3.70 |
| Unmasked-calculated* | 3.96 | 7.60 | 3.98 |

*Resolution estimate based on FSC curve calculated by comparison of deposited half-maps. The value from deposited half-maps intersecting FSC 0.143 CUT-OFF 3.96 differs from the reported value 3.52 by more than 10 %

For Manuscript Review

### 9 Map-model fit (i)

This section contains information regarding the fit between EMDB map EMD-77097 and PDB model 13JR. Per-residue inclusion information can be found in section 3 on page 5.

#### 9.1 Map-model overlay (i)

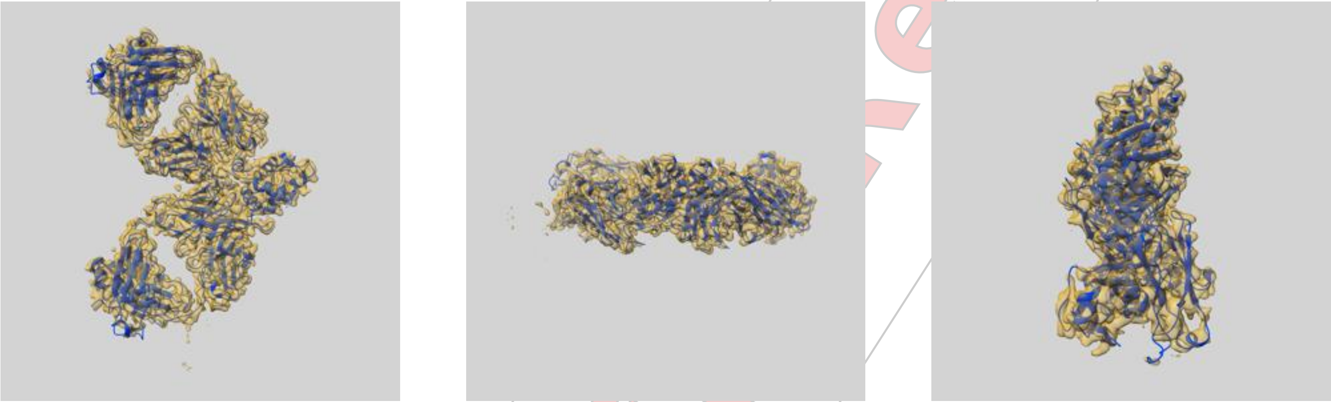

X Y Z

The images above show the 3D surface view of the map at the recommended contour level 0.12 at 50% transparency in yellow overlaid with a ribbon representation of the model coloured in blue. These images allow for the visual assessment of the quality of fit between the atomic model and the map.

For Manuscript Review

#### 9.2 Q-score mapped to coordinate model (i)

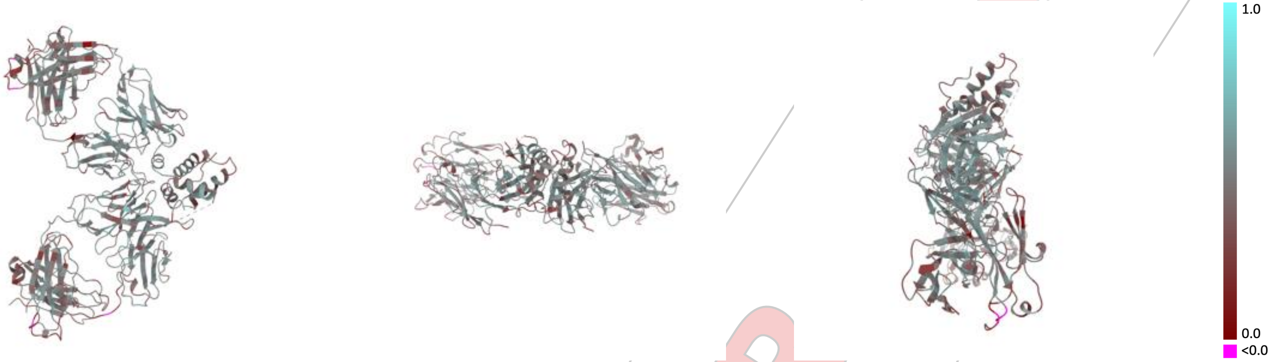


X Y Z

The images above show the model with each residue coloured according its Q-score. This shows their resolvability in the map with higher Q-score values reflecting better resolvability. Please note: Q-score is calculating the resolvability of atoms, and thus high values are only expected at resolutions at which atoms can be resolved. Low Q-score values may therefore be expected for many entries.

#### 9.3 Atom inclusion mapped to coordinate model (i)

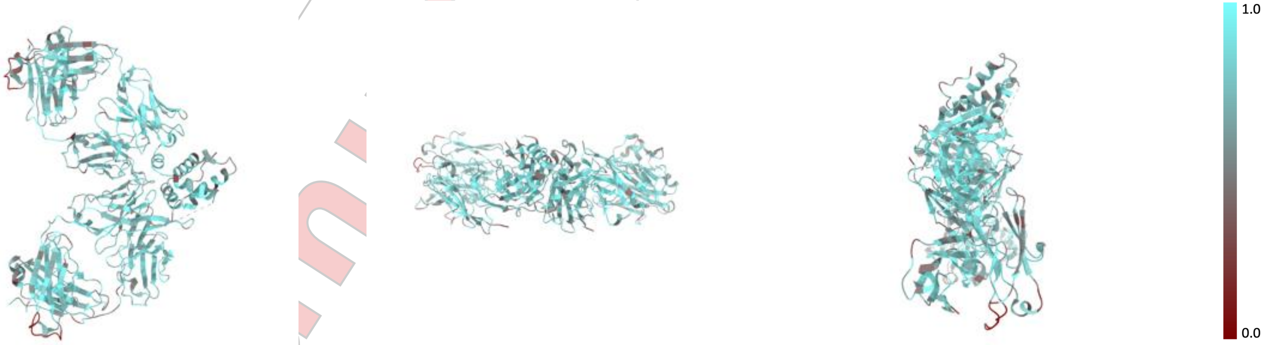


X Y Z

The images above show the model with each residue coloured according to its atom inclusion. This shows to what extent they are inside the map at the recommended contour level (0.12).

For Manuscript Review

#### 9.4 Atom inclusion (i)

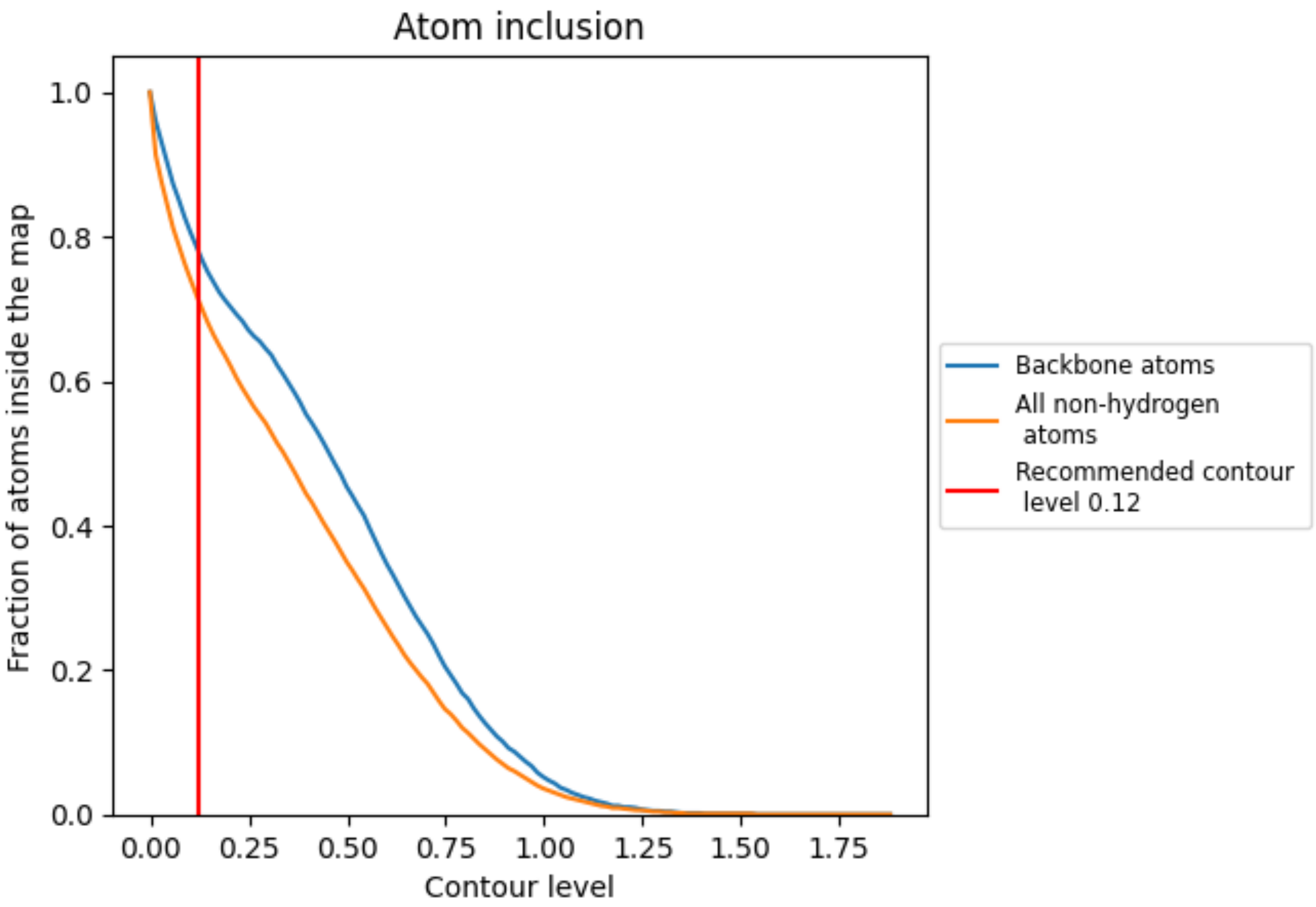


At the recommended contour level, 78% of all backbone atoms, 71% of all non-hydrogen atoms, are inside the map.

For Manuscript Review

#### 9.5 Map-model fit summary (i)

The table lists the average atom inclusion at the recommended contour level (0.12) and Q-score for the entire model and for each chain.

| Chain | Atom inclusion | Q-score |
|---|---|---|
| All | 0.7140 | 0.4480 |
| A | 0.7270 | 0.4640 |
| B | 0.7260 | 0.4550 |
| C | 0.7160 | 0.4540 |
| D | 0.6980 | 0.4270 |
| E | 0.6910 | 0.4300 |

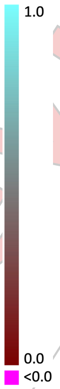


For Manuscript Review
